# Disulfiram inhibits *M. tuberculosis* growth by altering methionine pool, redox status and host-immune response

**DOI:** 10.1101/2020.09.01.277368

**Authors:** Deepika Chaudhary, Mardiana Marzuki, Andrea Lee, Rania Bouzeyen, Avantika Singh, Tannu Priya Gosain, Saqib Kidwai, Courtney Grady, Kholiswa Tsotetsi, Kiran Chawla, Foo Shihui, Josephine Lum, Sonu Kumar Gupta, Nisheeth Agarwal, Liana Tsenova, Yashwant Kumar, Bernett Lee, Pradeep Kumar, Krishan Gopal Thakur, Ramandeep Singh, Amit Singhal

## Abstract

Methionine biosynthetic pathway, essential for the growth of *Mycobacterium tuberculosis* (*Mtb*) in the host, represents an attractive target for the development of novel anti-tuberculars. Here, we have biochemically characterized homoserine acetyl transferase (HSAT viz. MetA) of *Mtb*, which catalyses the first committed step of methionine and S-adenosylmethionine (SAM) biosynthesis. High-throughput screening of a 2300 compound library resulted in identification of thiram, an anti-fungal organosulfur compound, as the most potent MetA inhibitor. Further analysis of thiram analogs led to the identification of orally bioavailable disulfiram (DIS, an anti-alcoholism FDA approved drug) as a novel inhibitor of MetA. Both thiram and DIS restricted the growth of drug-sensitive and drug-resistant *Mtb* strains in a bactericidal manner. ThermoFlour assay demonstrated direct binding of DIS with MetA. Metabolomic and transcriptomic studies showed DIS mediated perturbation of methionine and redox homeostasis, respectively, in *Mtb*. In concordance, the effect of DIS on *Mtb* growth was partially rescued by supplementation with either L-methionine as well as N-acetyl cysteine, suggesting a multi-target killing mechanism. In *Mtb*-infected mice, DIS administration restricted bacterial growth, increased efficacy of isoniazid, ameliorated lung pathology, modulated lung immune cell landscape and protective immune response. Taken together, our results demonstrate that DIS can be repurposed for designing an effective anti-tubercular therapy.

## INTRODUCTION

*Mycobacterium tuberculosis* (*Mtb*), the etiological agent of tuberculosis (TB), is a leading cause of infections in the world ^1^. The success rate of the current chemotherapy against susceptible and drug resistant (DR) *Mtb* are 85% and 55% respectively, specifically when the drug regimen is strictly adhered. Furthermore, due to prolonged duration of chemotherapy and toxicity associated with first-line TB drugs, there is always a risk of poor patient compliance resulting in emergence of DR strains. Hence, there is a need to develop drugs that possess a novel mechanism of action and that can be administered with other front-line TB drugs in order to shorten the duration of chemotherapy ^1, 2^.

Numerous studies involving either target based or phenotypic screening have identified various metabolic pathways that can be exploited to curtail DR *Mtb* ^3, 4, 5, 6^. These pathways include enzymes involved in either central carbon metabolism or lipid or amino acid or cofactor biosynthesis that have been shown to be essential for *Mtb* survival *in vivo* ^3, 5, 7^. Among these, amino acid biosynthetic pathways are corroborated targets for anti-*Mtb* drugs ^3, 8, 9^, since enzymes involved in this pathway lack human orthologs and are essential for *Mtb* growth in the host ^10, 11, 12, 13^. L-aspartate pathway, which synthesizes four essential amino acids L-threonine, L-lysine, L-methionine, and L-isoleucine plays a pivotal role in cell wall biosynthesis, translation and cellular metabolism of *Mtb* ^14^. Disruption of L-aspartate pathway suggest that both L-methionine and L-threonine are indeed required for *Mtb* to establish infection in the host, therefore, demonstrating *in vivo* essentiality of the enzymes involved in this pathway ^3^.

L-methionine is a precursor for S-adenosyl methionine (SAM), which is an essential co-factor for various enzymatic reactions. The enzyme homoserine acetyl transferase (HSAT) performs the first committed step in L-methionine biosynthesis by converting L-homoserine to O-acetyl-L-homoserine ^15, 16, 17^. The genome of *Mtb* encodes for MetA (Rv3341), which shares a 30-40% sequence identity with HSAT homologs from *C. glutamicum* ^16^, *L. interrogans* ^18^ and *H. influenzae* ^19^. *Mtb*^MetA^ strain showed auxotrophy for L-methionine during *in vitro* growth condition and was attenuated for growth in macrophages, immunocompetent and immunocompromised mice ^12^, indicating the essentiality of MetA *in vivo*.

In this study, we have performed target based screen and identified FDA-approved anti-alcoholism drug, disulfiram (DIS), as a small molecule inhibitor of *Mtb* MetA. Using an array of approaches we show that DIS (i) targets MetA enzyme and rewires L-methionine metabolism in *Mtb*; (ii) inhibits the growth of *Mtb in vitro* and in macrophages; (iii) alters redox homeostasis in *Mtb*; (iv) enhances efficacy of isoniazid (INH) in macrophages and *Mtb* infected mice; and (v) reprograms immune response, leading to reduced *Mtb* growth and pathology in lung tissues.

## RESULTS

### High through put screening identifies Thiram as MetA inhibitor that possesses whole cell activity

The genome of *Mtb* encodes for enzymes belonging to L-aspartate pathway, most of which have been predicted to be essential *in vitro* (Fig. 1a) ^7, 20^. Previous studies have shown that deletion of the *metA* gene resulted in depletion of L-methionine and SAM leading to growth defects of *Mtb in vitro*, macrophages and host tissues ^12^. Therefore, we first reassessed the essentiality of MetA by generating *metA* knock down strain using dCas9 based CRISPRi approach ^21^. The *metA* transcript levels were reduced by ∼16.0-20.0-fold in the knock down strain compared to the parental strain (Fig. S1a). The *metA* knock down strain showed reduced growth *in vitro* (Figs. 1b and S1b), reconfirming essentiality of L-methionine biosynthetic pathway for *Mtb* growth *in vitro*.

**Figure 1.**
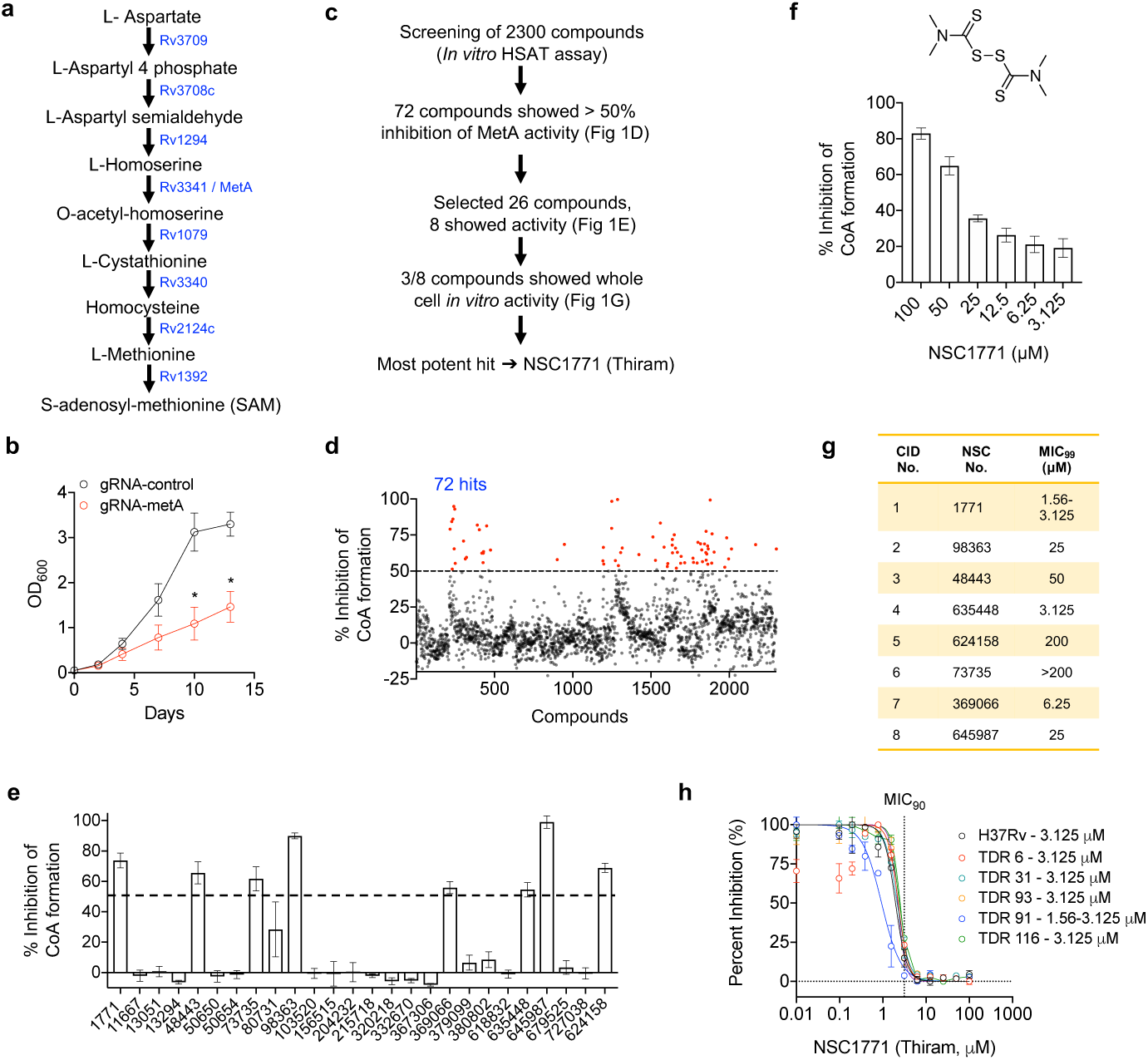
**Target based screen against MetA identifies thiram with whole cell activity. a**, The schematic showing L-methionine and SAM biosynthetic pathway in *Mtb*. The gene ID of different enzymes involved are shown. **b**, Growth curve of control and *metA* knock down *Mtb* strain was performed by measuring OD_600nm_ at regular intervals. **c**, Flow-chart depicting experimental steps in HTS for the identification of MetA inhibitors starting from the library screening to the *in vitro* MIC_99_ determination assays. **d**, Scatter plot showing average of percentage inhibition of HSAT activity (presented as % inhibition of CoA formation) in the presence of small molecules (100 μM) obtained from the 2300 compound library. 72 hits inhibiting MetA activity by atleast 50% are shown in red. **e**, Percentage inhibition of CoA formation by 26 compounds (out of 72 hits) at 100 μM concentration. **f**, MetA activity assay in the presence of different concentrations of NSC1771 (thiram, structure shown). **g**, MIC_99_ values obtained for 8 active hits identified from panel 1E against *Mtb in vitro*. The data shown is representative of three independent experiments. **h**, The effect of thiram on the growth of H37Rv and various DR clinical strains. MIC_90_ values are shown. The data in g and h is representative of two independent experiments. The data in b, e and f is mean ± SEM and is representative of three independent experiments. * P<0.05 by Mann-Whitney *U*-test.

Next, we cloned, purified and biochemically characterized recombinant histidine-tagged MetA. The existing structures from *Mtb, M. smegmatis, M. abscessus,* and *M. hassiacum* revealed that MetA predominately exists in dimeric oligomeric state ^22, 23^. We performed sedimentation velocity analytical ultracentrifugation (SV-AUC) experiments to determine the MetA oligomeric state(s) in solution. We observed three peaks with apparent molecular weights of 43 kDa, 82.5 kDa and 158 kDa corresponding to monomeric, dimeric and tetrameric oligomeric states, respectively, in solution (Fig. S1c). SV-AUC data further suggested that MetA exists predominantly as a dimer in solution, in concordance with the state observed in the crystal structures. The population of the oligomeric species remained unchanged at three different concentrations used in this study (Fig. S1c). The monomeric, dimeric and tetrameric peak fractions, showed comparable HSAT activity, resulting in the formation of 175 μM coenzyme A (CoA) in 5,5’-dithiobis-2-nitrobenzoic acid (DTNB) based assay (data not shown). Steady state kinetics revealed that the formation of O-acetyl homoserine followed Michaelis Menten kinetics with *K_m_* of 328 μM and *k_cat_* of 44.12 min^-1^ for acetyl-CoA (Fig. S1d). The catalytic efficient constant (*k_cat_/K_m_*) for MetA was 0.134 μM^-1^min^-1^. Investigation into the substrate specificity revealed that in addition to acetyl-CoA (AC), MetA could also utilize propionyl-CoA (PC) as an acyl-donor but not succinyl-CoA (SC) (Fig. S1e). Furthermore, MetA was unable to efficiently utilize L-serine or L-threonine as substrate as approximately 15% activity was observed in comparison to L-homoserine containing reactions (Fig. S1f). We observed comparable MetA activity upon inclusion of either Ca^2+^ or Mg^2+^ or Fe^3+^ in assay buffer, however, the presence of Zn^2+^ ions in assay buffer significantly inhibited the enzymatic activity (data not shown). Furthermore, MetA activity was independent of pH of the assay buffer (Fig. S1g) and maximal activity was attained within the initial 10 mins of the enzymatic reaction (Fig. S1h).

Using the above standardised parameters, we performed a high-throughput screen (HTS) in duplicates using a 2300 compound library to identify inhibitors for MetA activity *in vitro* (Fig. 1c). In our preliminary screen, 72 compounds were able to inhibit the formation of CoA by more than 50% *in vitro* at 100 μM concentration (Fig. 1d). To eliminate false positives, inhibition assays were repeated using 26 primary hits. Among these, 8 compounds, NSC1771, NSC48443, NSC73735, NSC98363, NSC369066, NSC635448, NSC645987 and NSC624158, inhibited MetA activity by more than 50% at 100 μM concentration (Fig. 1e). These active compounds also showed dose dependent inhibition of MetA enzymatic activity *in vitro* (Figs. 1f and S2). Next, we performed assays to evaluate the *in vitro* antitubercular activity of the identified MetA inhibitors. NSC1771, NSC635448 and NSC369066 were found to be the most potent compounds (Fig. 1g), with NSC1771 (also known as Thiram) displaying most potent MIC_90_ value in the range of 1.56-3.12 μM against both drug-sensitive and DR strains (Fig. 1h), thereby suggesting it targeted a metabolic pathway that was distinct from the targets of current first- and second-line TB drugs. Taken together, our target-based screen identified thiram as a MetA inhibitor with whole cell activity against *Mtb*.

### Disulfiram, an anti-alcoholism drug weakly binds MetA and alters metabolic profiles of *Mtb in vitro*

Thiram, the most active compound in our HSAT based screen is an organosulfur-based drug, used mainly as a fungicide in plants ^24^. In animals, it has few applications where it is applied directly to the skin or incorporated into sun screen or soaps ^24^. To identify an orally bioavailable thiram analogue we ordered 6 compounds from NIH-DTP (Fig. S3). All analogues except NSC1339 and NSC93058 inhibited MetA activity by >50% at 100 μM concentration (Fig. 2a). However, among the 4 active analogues, only DIS displayed an *in vitro* MIC_99_ values similar to thiram viz. 3.12 μM (Fig. 2a). DIS inhibited MetA activity *in vitro* in a dose dependent manner (Fig. 2b) and also showed *in vitro* anti-tubercular activity against DR *Mtb* strains (Fig. 2c).

**Figure 2.**
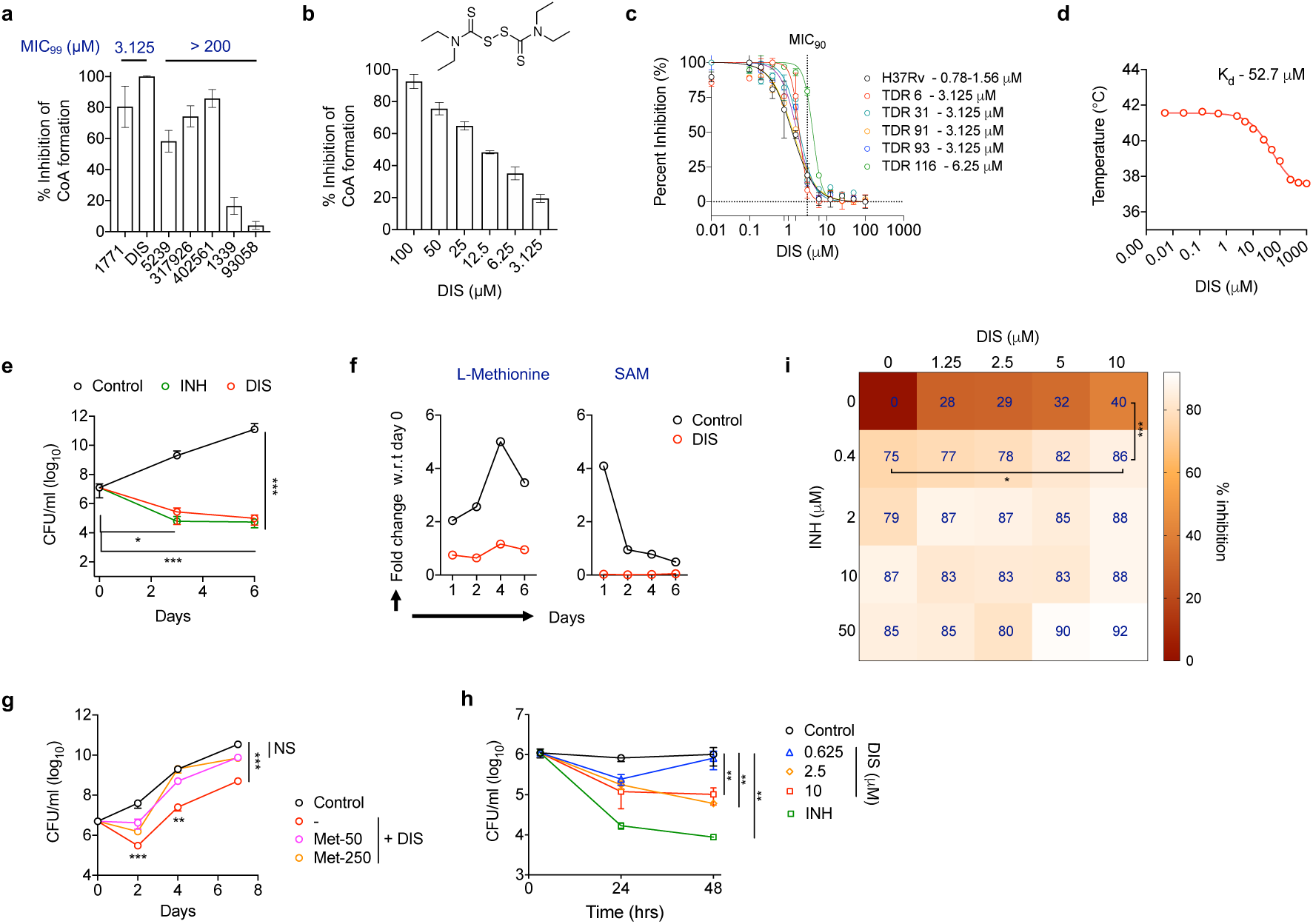
**DIS binds *Mtb* MetA, inhibits methionine biosynthesis, restricts intracellular *Mtb* growth and enhances INH efficacy. a**, MetA enzymatic activity in the presence of 6 thiram analogs was determined at 100 μM concentration. MIC_99_ values of respective analogs are depicted. **b**, MetA activity assay in the presence of different concentrations of disulfiram (DIS, structure shown). **c**, The effect of DIS on the growth of H37Rv and various DR clinical strains. MIC_90_ values are shown. **d**, Thermoflour assay demonstrating binding of DIS with MetA. The dissociation constant (K_d_) was calculated as 52.7 μM. **e**, Time kill kinetics of DIS and INH against *M. bovis* BCG. Early-log phase cultures of *M. bovis* BCG were exposed to 10x MIC_99_ concentration of either DIS or INH and cfu numbers were determined at day 3 and 6 post-exposure. **f**, Measurement of L-methionine and SAM levels in untreated and 10 μM DIS-treated *Mtb* at different time points. **g**, The early-log phase cultures of *M. bovis* BCG were exposed to 1x MIC_99_ DIS in the presence or absence of 50 μg/ml or 200 μg/ml L-methionine and bacterial numbers were determined at day 2, 4 and 7 post-exposure. **h**, Intracellular *Mtb* growth at 24 and 48 hr after treatment of primary human monocytes with vehicle control (DMSO) or INH (10 µM) or DIS (0.625-10 µM). The data in e, g and h is mean ± SEM and is representative of two independent experiments. * P<0.05, ** P<0.005, *** P<0.0005 by Kruskal Wallis test. **i**, Intracellular BCG growth after treatment of THP-1 cells with combination of various concentrations of INH and DIS for 24 hr. Representative of 2 experiments, mean of % inhibition of *Mtb* growth is indicated in each box. * P<0.05, *** P<0.0005, 2way ANOVA.

DIS is an FDA approved drug used for treatment of chronic alcoholism and sold under the trade name, Antabuse ^25^. The drug is currently under clinical investigations against certain types of cancer and HIV-1 infection ^26, 27^. Since DIS is an electrophile that can readily form disulfide with thiol-bearing molecules, we next performed ThermoFlour assays to determine the strength of DIS binding with recombinant MetA in order to eliminate the possibility of non-specific inhibition in our assay (Fig. 2a,b). A negative shift of about 4°C in MetA Tm reveals that DIS binds MetA with a K_d_ of ∼ 52 μM (Fig. 2d). No thermal shift was observed with isoniazid (INH), a negative control in the ThermoFlour assays (data not shown). Next, we performed *in vitro* killing experiments, and found that the exposure of early-log phase *M. bovis* BCG cultures to 10x MIC_99_ DIS resulted in 50- and 120-fold killing at 3- and 6-days post-exposure, respectively (Fig. 2e). The levels of killing seen in 10x MIC_99_ DIS treated samples was almost comparable to that seen upon exposure to 10x MIC_99_ INH (Fig. 2f). In order to further validate MetA to be the target for DIS, we compared the intracellular levels of L-methionine and SAM in untreated and DIS treated *Mtb*. The reduced levels of L-methionine and SAM in DIS-treated cultures in comparison to untreated cultures, indicated inhibition of MetA *in vitro* (Fig. 2f). Interestingly, supplementation of medium with L-methionine partially abrogated the killing activity of 1x MIC_99_ DIS *in vitro* (Fig. 2g). Of note, *Mtb*^MetA^ mutant strain also showed reduction in levels of L-methionine and SAM in liquid cultures ^12^.

DIS treatment also inhibited the growth of intracellular *Mtb* (∼ one log_10_ fold) in primary human monocytes in a dose-dependent manner (Fig. 2h). Further, we performed checkerboard assays to determine the synergistic effect of DIS with INH against intracellular mycobacteria. The data indicates that combination of 10 μM DIS along with 0.4 μM INH, resulted in an increased inhibition of the intracellular mycobacterial growth compared to either drug alone (Fig. 2i). Taken together, our data suggest that DIS (i) targets methionine biosynthetic pathway, (ii) inhibits *Mtb* growth in a bactericidal manner and (iii) enhances the efficacy of INH against intracellular bacteria.

### Disulfiram perturbs redox homeostasis in *Mtb*

DIS has been shown to inhibit several metabolic enzymes such as urease, aldehyde dehydrogenase and triose phosphate isomerase in various microorganisms ^28, 29, 30^. Therefore, to further delineate the mechanism behind DIS mediated killing *in vitro* we compared the transcriptomic profiles obtained from *Mtb* exposed to either 1x or 10x MIC_99_ of DIS with DMSO treated samples. Differential gene expression (DGE) analysis (FDR < 0.05 and log_2_ fold change > 1.5 or < −1.5) identified that the transcript levels of 107 genes (59 upregulated and 48 downregulated) were altered in 1x MIC_99_ DIS treated samples in comparison to control (Fig. 3a and Table S1). Majority of the identified DEGs were categorized as either conserved hypothetical proteins or regulatory proteins or the ones involved in cell wall processes or intermediary metabolism (Fig. 3b). Downregulated genes included enzymes involved in (i) cell division *(sepF*, *ftsW*), (ii) copper resistance (*mctB*) (iii) cutinase activity (*cfp21*) and (iv) respiration (type I NADH dehydrogenase; *nuoF*, *nuoH* and *nuoE*), all of which have been shown to be important for *Mtb* growth *in vitro* or *in vivo* ^31, 32^. The exposure of *Mtb* cultures to 10x MIC_99_ DIS resulted in differential expression of 119 transcripts (Fig. S4a and Table S1), 53 of which were common with the DEGs observed in 1x MIC_99_ DIS treated samples (Fig. 3c).

**Figure 3.**
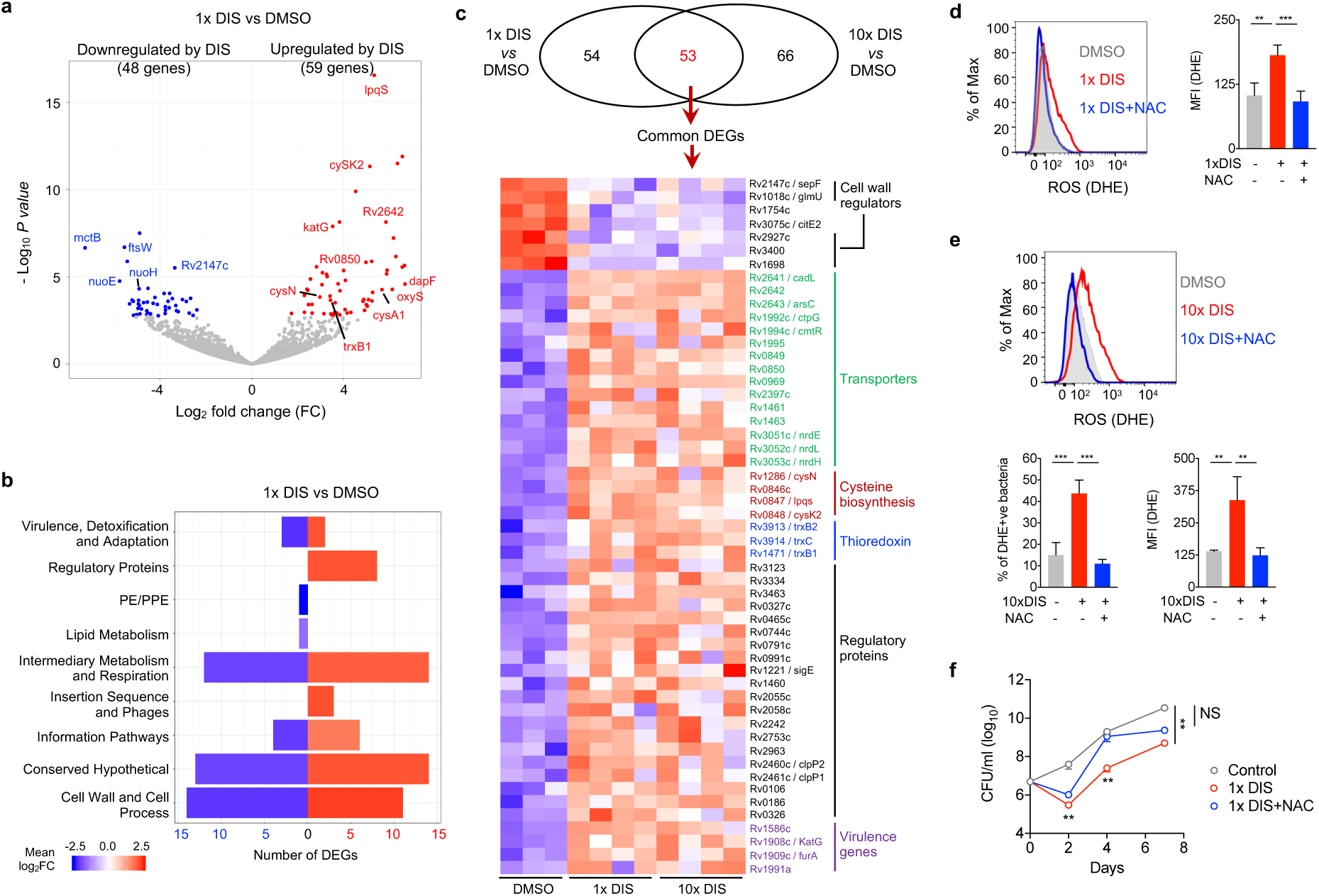
**DIS alters transcription profile and redox homeostasis of *Mtb in vitro*. a**, Volcano plot showing the gene expression comparison between 1x MIC_99_ DIS vs DMSO treated, where individual genes are represented as single dots with the -log_10_ *P*-value on the y axis and the log_2_ fold change on the x axis. The transcript shown in red and blue are significantly upregulated and downregulated, respectively. Non-significant genes are coloured in grey. Selected significant genes are shown with their gene names as the labels. **b**, The identified DEGs in panel A were categorized into various functional categories as listed in Tuberculist. The total number of downregulated (blue) and upregulated (red) genes in each category are depicted as bars. The color intensity reflects the magnitude of the log_2_ fold change (FC) for the set of genes in that category. **c**, Venn diagram demonstrating common *Mtb* genes, expression of which was altered upon exposure to either 1x MIC_99_ or 10x MIC_99_ DIS in comparison to DMSO treated samples. The heat map corresponding to the 53 common genes is shown, where each column represents a single sample. For each gene expression in log_2_ RPKM were *z*-score scaled and shown as red for increased expression and blue for reduced expression. The genes are grouped by their annotated functions labelled on the right. **d and e**, *M. bovis* BCG cultures were exposed to different combinations and intracellular ROS was measured by flow cytometry using 5μM DHE. **f**, The killing curves of *Mtb* were performed upon exposure to DIS in the presence or absence of 10 mM NAC and bacterial numbers were determined at day 2, 4 and 7 post-exposure. The data in d-f is mean ± SEM and is representative of two independent experiments. ** P<0.005, *** P<0.0005 by Kruskal Wallis test.

In agreement with 1x MIC_99_ DIS DEGs, 10x MIC_99_ DIS DEGs were also categorized as either conserved hypothetical proteins or regulatory proteins or the ones involved in cell wall processes or intermediary metabolism (Fig. S4b). Various transcripts such as *espR*, *hupB*, *mctB*, *senX3*, *drrC*, *rpsC*, *dapB*, *Rv2927c*, *murB*, *lgt*, *dnaB*, *ilvC*, etc which are essential for *Mtb* growth or virulence were found to be downregulated upon exposure to 10x MIC_99_ DIS (Table S1). The more detailed assessment of common 53 DEGs, showed upregulation of genes involved in (i) sulfur metabolism and cysteine biosynthesis (*cysA1*, *cysK2* and *cysN*), (ii) DNA replication and repair (*nrdE*, *nrdH* and *nrdL*), (iii) oxidative stress (*trxC*, *trxB1* and *trxB2*), (iv) transporters (*arsC*, *ctpG*, *cmtR*, *cadL*) and (v) regulatory proteins (*sigE*, *Rv3334*, *Rv3463*, *Rv1086*) (Fig. 3c). The observed downregulation of transcripts belonging to cell wall regulators such as *sepF*, *glmU*, *Rv3400*, *Rv1698* indicated that DIS might effect *Mtb* cell wall integrity (Fig. 3c).

The increased expression of thioredoxins, cysteine biosynthesis enzymes and type I NADH dehydrogenases genes (Fig. 3a,c), suggested that exposure to DIS may result in the alteration of redox homeostasis and generation of reactive oxygen species (ROS), which has been shown to restrict *Mtb* growth ^33, 34^. In agreement, DIS treatment (either by 1x MIC_99_ or 10x MIC_99_) increased the ROS production in *Mtb* as early as 6h (Fig. 3d,e), which was maintained till 24h post-treatment (Fig. S4c,d). The addition of ROS-quenching agent N-acetyl Cysteine (NAC) in liquid cultures abolished the ability of DIS to induce ROS (Fig. 3d,e and Fig. S4b,c) and partially abrogated 1x MIC_99_ DIS mediated killing of *Mtb* (Fig. 3f). Taken together our data suggest that DIS exposure alters the redox homeostasis of *Mtb* and in addition to inhibition of MetA and copper homeostasis ^35^, ROS generation is another mechanism for DIS mediated killing *in vitro*.

### Disulfiram reduces *Mtb* burden and tissue pathology in mice

We next evaluated the *in vivo* efficacy of DIS in acute TB murine model where *Mtb*-infected C57BL/6 mice were treated with DIS starting from day 7 post-infection (Fig. 4a). DIS treated mice showed decreased bacillary load in the lung and spleen compared to untreated mice at 4 weeks (wk) post-treatment (p.t.) or 35 days post-infection (p.i) (Fig. 4b,c). In order to significantly improve the control of TB, drugs should also promote resolution of tissue pathology in addition to accelerating bacillary clearance ^36^. Therefore we performed histopathologic examination of the lungs from *Mtb*-infected mice treated with DIS at 4 wk p.t. Evaluation of the lungs of uninfected mice showed completely intact lung parenchyma (Fig. 4d). Examination of lungs from untreated *Mtb*-infected mice revealed peribronchial diffuse coalescent lesions with large numbers of infiltrating macrophages and lymphocytes, and numerous intracellular acid-fast-bacilli (AFB) (Fig. 4d,e). Interestingly, DIS treatment was associated with increased lymphocyte infiltration and formation of aggregates in the infected tissues (Fig. 4d), and reduced numbers of AFB (Fig. 4e). These properties have been associated with improved *Mtb* control in mice ^37^. Taken together, our results indicate that DIS treatment controls *Mtb* growth and ameliorates TB pathology, suggesting the potential of DIS as an adjunct to current TB therapy.

**Figure 4.**
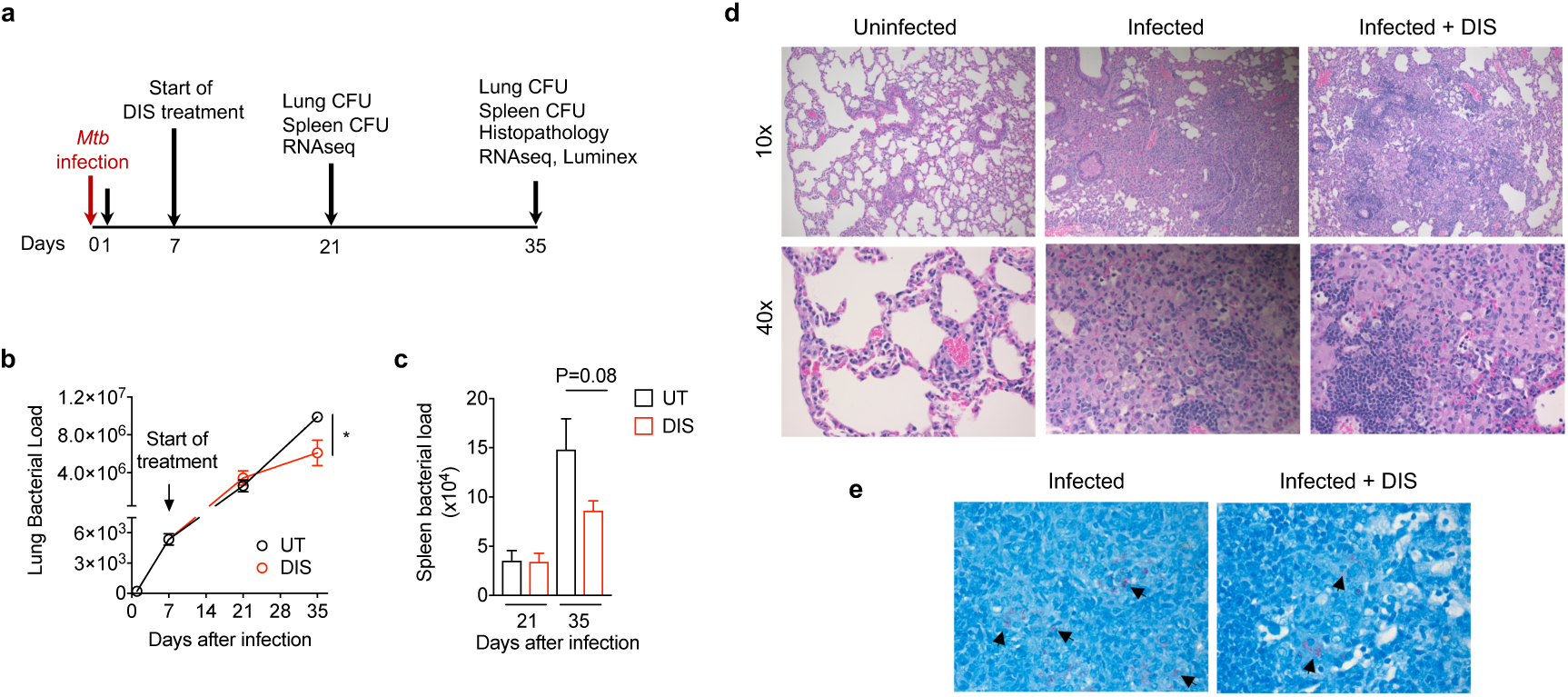
**DIS inhibits proliferation of *Mtb* and TB-associated pathology in mouse lungs. a**, Experimental scheme of acute *Mtb* C57BL/6 mouse model. **b and c**, *Mtb*-infected mice were treated with DIS (160mg/kg) starting 7 days post-infection. Bacterial loads were enumerated in the lungs (b) and spleens (c) at different time points. UT, Untreated *Mtb*-infected mice; DIS, DIS-treated *Mtb*-infected mice. The data shown in this panel is mean ± SEM obtained from 4-6 mice per time point. \**P* < 0.05 by Mann-Whitney *U*-test. **d**, Light micrographs of representative H&E stained lung sections of mice, 4 weeks post-treatment. Uninfected, *Mtb*-infected and DIS-treated *Mtb*-infected lungs were observed at 10x and 40x magnification. **e**, Light micrographs of AFB staining of representative lung sections of mice, 4 wk p.t. 60x magnification. AFB is indicated by arrow. *n* = 5 mice per group per time point.

### Disulfiram treatment enhances protective innate immune response in lung tissues

To further investigate the mechanism by which DIS restricts *Mtb* growth in the mouse lung, a genome-wide transcriptional analysis using total RNA isolated from uninfected or untreated or or DIS-treated *Mtb*-infected mouse lungs was performed. At 4 wk p.t 2846 transcripts (FDR < 0.05) were differentially expressed in response to infection (*Mtb*-infected vs uninfected, Table S2). Treatment of infected mice with DIS altered the expression of 169 genes (54 decreased and 115 increased) as compared to the untreated group (Fig. S5a and Table S2). Ingenuity pathway analysis (IPA) of 169 differentially expressed genes (DEGs) revealed that DIS treatment (i) increased the transcript levels of genes associated with mTOR (mechanistic target of rapamycin) signalling and MIF (Macrophage Migration Inhibitory Factor)-mediated innate immunity; and (ii) decreased the expression of genes associated with oxidative phosphorylation (OXPHOS) and Wnt pathway (Fig. S5b). mTOR is crucial for restricting *Mtb* growth by inducing autophagy ^38^. MIF is a critical mediator of the innate immune response to *Mtb* ^39^. MIF deficient mice are susceptible to *Mtb* infection with increased lung pathology and decreased innate cytokine production such as TNF-α ^39^.

To further dissect the effect of DIS on lung immune response and TB pathogenesis, we next performed a time-course analysis using the RNA-seq data from uninfected or untreated or DIS-treated *Mtb*-infected mouse lungs at 21 and 35 days p.i. This resulted in an identification of five distinct gene clusters (Cluster 1 – 5; Fig. 5a and Table S3) that are differentially expressed between different groups. Cluster 3 and 5 included the genes which were upregulated by *Mtb* as early as 21 days p.i, however DIS treatment had no effect on the expression of these genes at either 21 and 35 days p.i (Fig. 5b). The transcript levels of genes in Cluster 1 initially increased in response to infection and subsequently decreased to the levels observed in uninfected animals (Fig. 5b). DIS treatment, however, resulted in constant increase in the expression of these 61 genes till 35 days p.i (Fig. 5b). The genes in the cluster 1 included mostly innate response genes such as *vim*, *mif*, *card9*, and *nod1* (Table S3), which are important for the control of *Mtb* infection in host cells/tissues ^39, 40, 41, 42^. We also noticed enrichment of MIF-related innate immunity pathways among cluster 1 genes (Fig. 5c). Cluster 2 included the genes associated with OXPHOS (*cox4I1* and *uqcr11*), expression of which was continuously decreased in infected tissues (Fig. 5b,d). DIS treatment resulted in reducing the expression of cluster 2 genes back to the level noticed in uninfected animals (Fig. 5b). The transcript levels of genes belonging to Cluster 4 were reduced in DIS-treated mice in comparison to untreated mice at 35 days p.i (Fig. 5b). Importantly pathways such as Wnt signaling that promotes TB infection were enriched among cluster 4 genes (Fig. 5e) ^43^.

**Figure 5.**
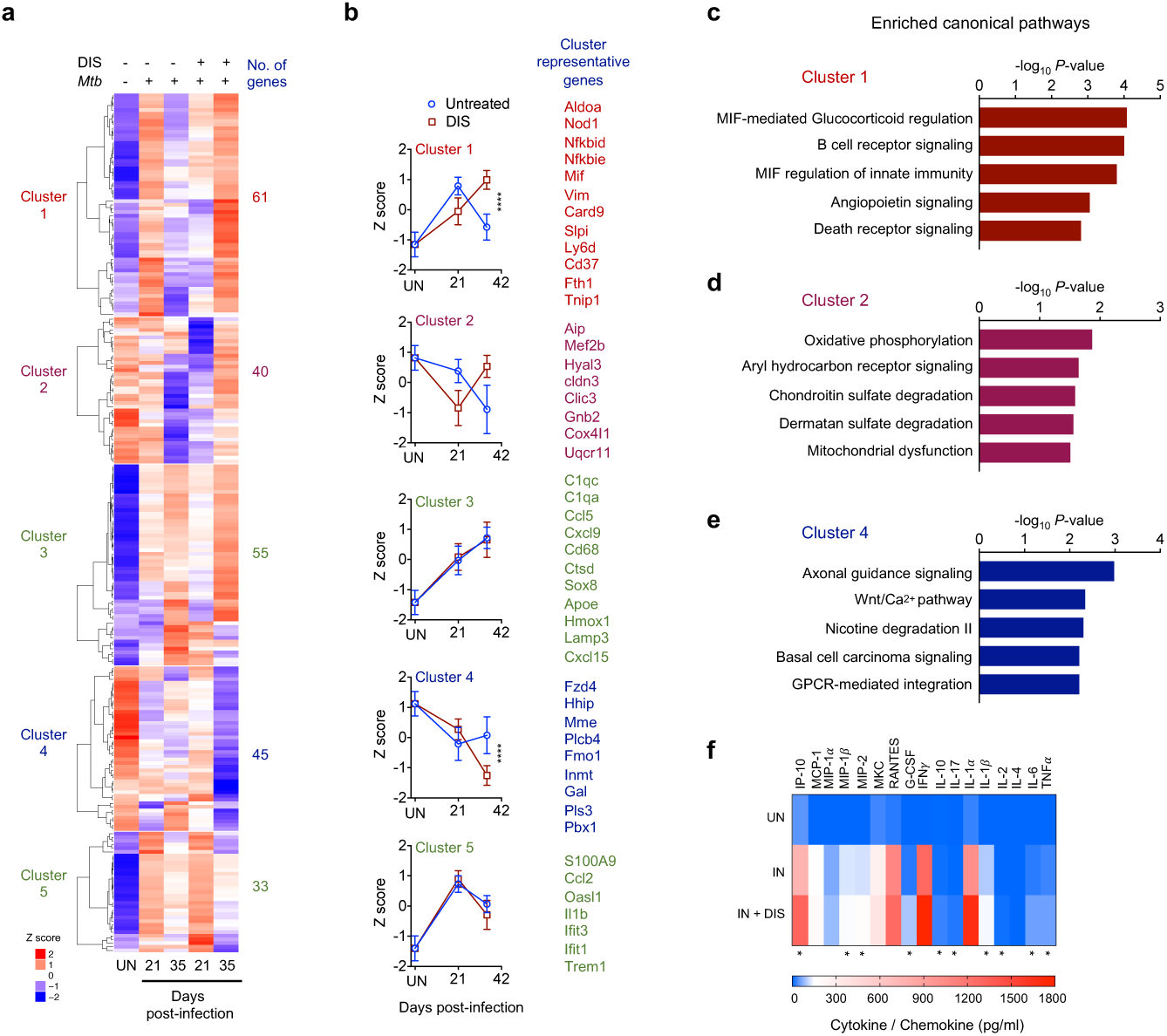
**DIS enhances lung immune response in *Mtb*-infected mice. a**, Heat map of genes (N=234) modulated by time, infection and DIS treatment (genes identified as significantly changed during infection as well as being significantly changed during time or DIS treatment). Gene expression in log_2_ RPKM values were averaged for each group of treatment and time point and then *z*-score normalized prior to visualization by heat map. The columns represents the average of the treatment and time point combination for individual genes. The genes were clustered using complete linkage hierarchical clustering using Pearson correlation as the distance metric and 5 clusters were observed and are presented. The number of genes in each of the clusters is shown on the right of the cluster. **b**, Line plots of the averaged *z*-scores computed in panel A for DIS treated and untreated *Mtb*-infected samples showing the differences in the profiles. Data shown in this panel is mean ± SD. A list of representative genes for each cluster are shown on the right of the line plots. Trend in cluster 1 and 4 is significant. ****P < 0.0001, 2way ANOVA. **c-e**, Ingenuity pathway analysis (IPA) of genes in cluster 1, 2 and 4, respectively. **f**, Chemokines and cytokines levels in the lung homogenates of mice, on day 35 after *Mtb* infection, measured by ELISA. UN, Uninfected; IN, *Mtb*-infected; IN + DIS, DIS-treated *Mtb*-infected. \**P* < 0.05, by Mann-Whitney *U*-test. N=4 mice per group per timepoint.

MIF has been shown to upregulate TLR4 expression by macrophages resulting in enhanced production of cytokine (including MIF), and other chemokine mediators ^44^. Hence, we measured cytokines and chemokines in the lung homogenates of DIS treated *Mtb*-infected mice. At 4 wk p.t, in comparison to infected mice, DIS-treatment resulted in increased levels of cytokines that constitute innate immune response such as TNF*α*, IL-6, IL-1*β*, and IL-17 in the lung homogenates (Fig. 5f). We also noticed increased levels of chemokines such as IP-10, MIP-1*β* and MIP-2 in DIS treated *Mtb*-infected mice (Fig. 5f). These chemokines/cytokines are essential for survival following *Mtb* infection and known to potentiate anti-TB immunity^45^. Taken together, our results indicates that DIS regulates immunoregulatory mechanisms in the lungs of infected mice leading to the enhancement of protective immune response.

### Disulfiram enhances *Mtb* clearance by anti-TB drugs in mice and modulates lung immune cell landscape

We next investigated the adjunctive efficacy of DIS with INH in the chronic mouse model experiment using C57BL/6 mice ^46^, where treatment was initiated at day 30 p.i. (Fig. 6a). After 4 wk of therapy at 60 days p.i mice treated with INH in combination with DIS showed reduced lung bacillary loads in comparison to INH treated mice (Fig. 6a). To further explore the differential effect of DIS on host immune response, pulmonary immune cells were phenotyped using flow cytometry. Analysis of macrophages at 4 wk p.t. deciphered three unique population *viz*. alveolar macrophages (AMs, CD64^+^F4/80^+^CD24^-^CD11b^-^CD11c^+^MHC-II^-^), interstitial macrophages (IMs, CD64^+^F4/80^+^CD24^-^CD11b^+^CD11c^-^MHC-II^hi^) and monocyte-derived macrophages (MDMs, CD64^+^F4/80^+^CD24^-^CD11b^+^CD11c^mid^MHC-II^lo^) (Fig. S6a,b). An increased frequency of AMs, the first line of immune defense against respiratory pathogens such as *Mtb* ^47^, was observed in the INH+DIS treated mice in comparison to both untreated and INH-treated groups (Fig. 6b). Lungs of INH and INH+DIS treated mice also had an increased (i) Ly6C^mid^CD43^+^ intermediate monocytes (int-Mo), and (ii) Ly6C^lo^CD43^+^ non-classical monocytes (nc-Mo, known as patrolling tissue repair monocytes), at 4 wk p.t (Fig. 6c). However, no differences on the infiltration of Ly6C^hi^CD43^-^ classical monocytes (c-Mo) was observed between different groups (Fig. 6c). Lungs of both INH and INH+DIS treated mice showed decreased conventional type 1 and 2 dendritic cells (cDC1 and cDC2, respectively), and neutrophils in comparison to untreated groups (Fig. S6c). No effect on the frequency of eosinophils was observed (Fig. S6c). Profiling of lung lymphoid cells (Fig. S7a), which are essential to control primary *Mtb* infection ^48^, showed no effect of treatment on the frequency of total CD3^+^, CD4^+^, CD8^+^ T cells; and NK cells in lung tissues (Fig. S7b). However, dissection of T cells based on CD44 and CD62L expression indicated an increased frequency of lung CD8^+^CD44^-^CD62L^+^ naïve T cells (T_N_) and decreased CD8^+^CD44^+^CD62L^-^ effector T cells (T_E_) in INH+DIS treated mice compared to the untreated or INH treated mice (Fig. 6d). This was paralleled with a similar trend in CD4^+^ T cell subsets (Fig. 6e), thus confirming the dominance of T_N_ over T_E_ cells among CD4^+^ and CD8^+^ T cells in lungs of INH+DIS treated mice (Fig. 6f). However, the frequency of CD8^+^ or CD4^+^ central memory T cells (T_CM_) remained unchanged between different groups (Fig. 6d,e). Overall, our results suggest that apart from direct anti-bacterial effect, DIS also results in the reprograming of protective immune response, which must be tightly regulated to maintain disease tolerance and host survival ^49, 50, 51^.

**Figure 6.**
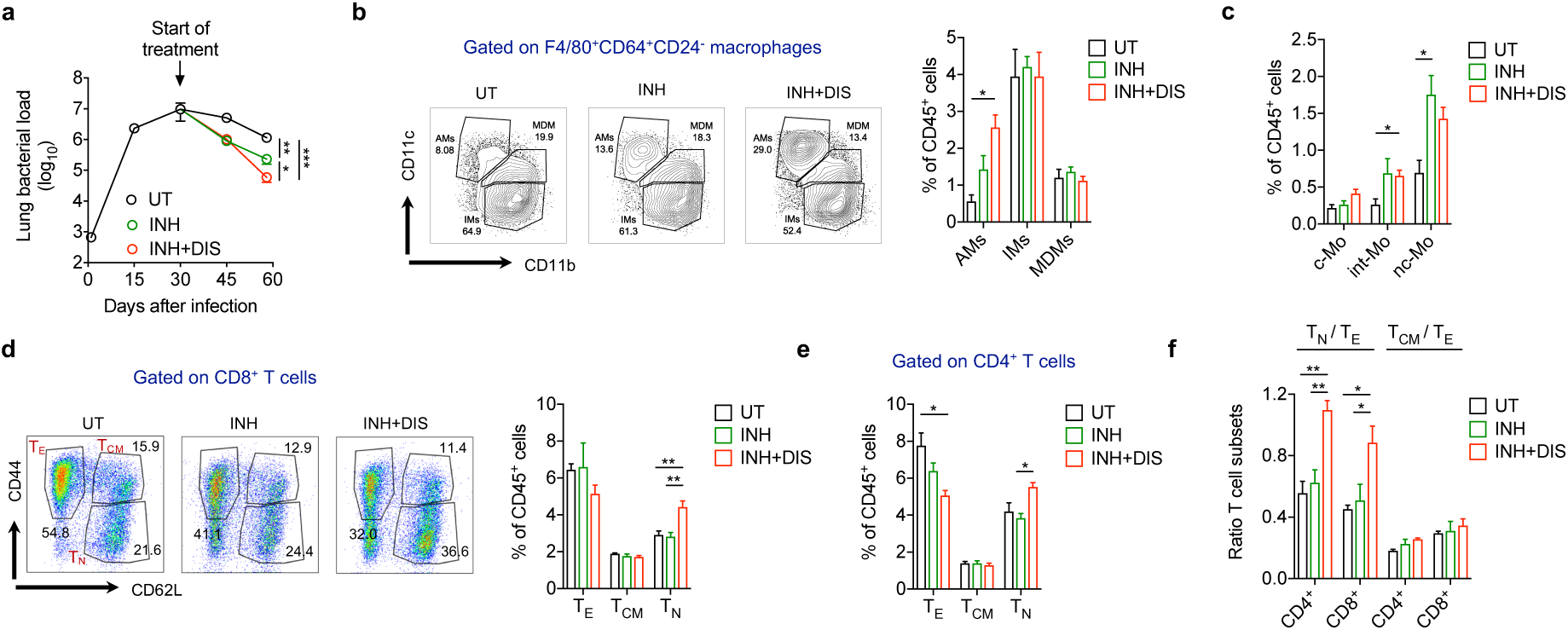
**DIS enhances INH efficacy *in vivo* and rewire lung immune cells landscape. a**, *Mtb*-infected mice were treated with INH (10 mg/kg) or INH+DIS (160 mg/kg) starting 30 days post-infection. Bacterial loads were enumerated in the lungs at different time points. UT, Untreated *Mtb*-infected mice; INH, INH treated *Mtb*-infected mice; INH+DIS, INH and DIS treated *Mtb*-infected mice. Statistical differences observed at 4 wk p.t has been indicated. **b**, Flow cytometric analysis of lung macrophages from mice at 4 wk p.t. Representative dot plots and compiled data is shown. AMs, Alveolar macrophages; IMs, interstitial macrophages; MDMs, monocyte-derived macrophages. **c**, Compiled flow cytometric analysis of lung monocytes from mice at 4 wk p.t. c-Mo, classical monocytes; int-Mo, intermediate monocytes; nc-Mo, nonclassical monocytes. **d**, Flow cytometric analysis of lung CD8^+^ T cell subsets from mice at 4 wk p.t. Representative dot plots and compiled data is shown. T_E_, effector T cells; T_CM_, central memory T cells; T_N_, naïve T cells. **e**, Compiled flow cytometric analysis of lung CD4^+^ T cell subsets from mice at 4 wk p.t. **f**, T_N_/T_E_ and T_CM_/T_E_ ratio among lung CD4^+^ and CD8^+^ T cells at 4 wk p.t. The data in A-F is mean ± SEM obtained from 4-6 mice per time point. *P < 0.05, **P < 0.005, ***P < 0.001 by one-way ANOVA.

## DISCUSSION

Methionine biosynthesis pathway is (i) involved in several biological processes such as translation initiation, synthesis of SAM, DNA and sulphur containing compounds, (ii) absent in mammalian cells, and (iii) predicted to be essential for the growth of microorganisms ^12, 52^. Indeed, *Mtb* needs a functional methionine/SAM biosynthesis for survival in the host ^12^, suggesting that the enzymes belonging to this pathway are attractive targets for designing new anti-*Mtb* therapy. Here, using a high-throughput target based screen, we identified 8 compounds including thiram (a plant fungicide) and its analogue disulfiram (DIS, a FDA approved anti-alcoholism drug, Antabuse) as inhibitors of *Mtb* homoserine acetyl transferase (HSAT, known as MetA), an enzyme that catalyzes the conversion of homoserine to O-acetyl-L-homoserine in L-methionine biosynthesis pathway ^15, 16, 17^. The identified scaffolds inhibited *Mtb* MetA’s acetyl transferase activity *in vitro* and were structurally different from the known fungal-HSAT inhibitors, CTCQC (a nucleotide substrate analogue) and Ebelactone A ^52, 53^. All initial hits except NSC635488, NSC369066 and thiram displayed anti-tubercular activity of greater than10μM in whole-cell based assay, which could be attributed to their ineffective transport and/or intracellular metabolism. Among thiram analogues, DIS was the most potent compound displaying comparable anti-tubercular against both drug-sensitive and DR *Mtb* strains. In agreement, DIS treatment resulted in the reduced levels of L-methionine and SAM *in vitro*. Supplementation of liquid cultures with L-methionine partially restored DIS mediated killing of *Mtb*, which phenocopied the effect of L-methionine supplementation on the growth of *Mtb*^MetA^ strain ^12^. Since methionine and SAM together play an important role in translation initiation and one-carbon metabolism ^54^, our finding suggest that DIS treatment might lead to a metabolic shutdown in *Mtb*. Importantly, SAM is also required for the biosynthesis of mycolic acid that is essential for viability, drug resistance and cell wall integrity of *Mtb* ^55^. However, DIS exposure did not affect the mycolic acid composition of *Mtb* (data not shown). Of note, DIS has demonstrated anti-bacterial activity against various *Mtb* clinical isolates including DR strains ^56^, methicillin-resistant *Staphylococcus aureus* ^57^ and *Pseudomonas aeruginosa* ^58^.

In addition to targeting methionine biosynthesis pathway, analysis of *Mtb* transcriptome suggested that DIS might alter redox homeostasis by inducing endogenous oxidative stress in *Mtb*. Indeed, we found DIS elevates endogenous ROS in *Mtb* and supplementation of liquid cultures with NAC partially restored DIS meditated killing *in vitro*. ROS is also accumulated in L-arginine deprived *Mtb* where it leads to oxidative damage of DNA, RNA and proteins resulting in cell death ^59, 60^. Similarly, drugs such as clofazimine, INH and vitamin C induce rapid death by ROS generation leading to oxidative damage ^61, 62, 63^. Since redox stress could lead to the generation of drug tolerant *Mtb* and high rates of TB therapy failure ^64^, targeting redox metabolism by FDA approved drugs might enhance mycobactericidal activities of first-line TB drugs ^64, 65^. Indeed DIS enhanced *Mtb* clearance by INH *in vitro* and *in vivo*, which might be attributed to the ROS generation property of both drugs. DIS has earlier been shown to enhance the efficacy of rifampicin, which also generate ROS in *Mtb* ^66^, in mice lungs ^56^. Importantly, maintaining an appropriate redox balance is critical to the clinical outcome because (i) several anti-TB prodrugs are only effective upon bioreductive activation and (ii) proper homeostasis of oxido-reductive systems is essential for *Mtb* survival, persistence and subsequent reactivation.

In the lungs of infected mice DIS ameliorated TB pathology and augmented anti-*Mtb* protective immune responses ^45^. Specifically, transcriptomic analysis of lung cells revealed that DIS treatment resulted in induction of MIF-mediated innate immunity, which has been shown to be critical for host immunity against TB ^39^. Mice deficient in MIF had increased lung pathology and decreased innate cytokine production (IL-6, TNF-α and IL-17) ^39^, all of which were increased in the lung of DIS-treated *Mtb*-infected mice. In addition, INH+DIS treated mice showed increased recruitment of AMs, cells which are the first immune cells to encounter invading *Mtb* and are responsible for the outcome of TB disease ^49^. AMs are known to influence responses of local T cells in the lung ^67^, which though are important to contain *Mtb* infection ^48^. However, when dysregulated, these cells can promote TB pathology and susceptibility ^50^, suggesting that a tightly regulated T cell response is fundamental for host to achieve better control of TB. By rewiring the effector T cell frequency DIS might be strengthening the host immune repertoire involved in combating *Mtb*. Of note, host-directed therapies targeting pathologic effector T cells have been proposed ^68^.

Taken together, this study expands our knowledge about the plausible anti-mycobacterial activities of DIS by highlighting that in addition to rewiring methionine metabolism, redox homeostasis, DIS also modulates host protective immunity against *Mtb*. We further speculate that this drug would also be able to clear intracellular bacteria that might acquire mutation during the course of chemotherapy. Lastly, since alcohol consumption increase the risk for TB by three-four fold ^69^, greater than that by diabetes and smoking, our study may entice the development of TB treatment strategies for this emerging co-morbidity.

## METHODS

### Chemicals, Strains and Growth conditions

The chemicals used in the present study were procured from Sigma, Merck unless mentioned. *Escherichia coli* XL-1 Blue and BL-21 (*λ*DE3, plysS) were used for cloning and protein expression studies, respectively. *Mtb* H37Rv strain was used for MIC_99_ determination and growth inhibition experiments. In the present study, various *E.coli* and mycobacterial strains were cultured in Luria Bertani and Middlebrook medium, respectively as previously described ^70^. The *metA* knock down strain of *Mtb* were constructed using CRISPRi approach as previously described ^21^. The growth of parental and *metA* knocked down *Mtb* strains was compared by measuring OD_600nm_ at regular intervals and spotting of 10.0 fold serial dilutions at designated time points. The genotypes for various *Mtb* clinical strains used for MIC_90_ determination are the following; TDR91: RIF^r^, EMB^r^, TDR116: INH^r^, PAS^r^, EMB^r^; TDR31: INH^r^, RIF^r^, EMB^r^, KAN^r^, SM^r^, CAP^r^; TDR6: RIF^r^, INH^r^, EMB^r^ and TDR93: INH^r^, EMB^r^

### Expression and Purification of HSAT protein

For expression studies, *metA* was PCR amplified and cloned into prokaryotic expression vector, pET28b. pET28b-*metA* was transformed into BL-21 (*λ*DE3, plysS) and protein expression was induced by the addition of 0.3 mM isopropyl *β*-D-1-thiogalactopyranoside (IPTG) at O.D_600nm_ ∼ 0.5 at 18°C, overnight. The induced cultures were lysed by sonication and recombinant protein was purified from clarified lysates using nickel-nitrilotriacetic acid (Ni-NTA) resin as per manufacturer’s recommendation. The elution fractions were concentrated and (His)_6_-Rv3341 was further purified by gel exclusion chromatography using Superdex 200 Increase 10/300 GL column (GE Healthcare). The purified fractions were pooled, dialysed, concentrated and stored as aliquots in enzyme storage buffer (50 mM NaH_2_PO_4_, 300 mM NaCl, 10% glycerol). The amount of protein in purified concentrated fractions was quantified using UV-Vis spectroscopy and Bradford reagent as per standard protocols.

### Analytical ultracentrifugation experiments

In order to determine the oligomeric state of MetA, AUC experiments were performed using Beckman-Coulter XL-A analytical ultracentrifuge equipped with a TiAn50 eight hole rotor. The protein samples at three different concentrations (10, 20 and 30 µM in 20 mM Tris-Cl, pH 7.5 and 150 mM NaCl buffer) were prepared, centrifuged at 40,000 rpm and the absorbance scans were recorded at 280 nm every 3 or 4 mins. To fit multiple scans at regular intervals, SEDFIT continuous distribution c(s) model was used ^71^. SEDNTERP (http://www.rasmb.bbri.org/) was used to find the solvent viscosity (η) and density (ρ).

### Biochemical characterization of MetA enzyme

To optimise the assay condition for MetA enzyme, various parameters such as buffer pH, ion concentration, substrate specificity, and reaction time were standardised. The enzymatic activity was measured spectrophotometrically at 412 nm using 0.2 mM Ellman’s reagent ^72^. For *K_m_, V_max_, k_cat_/K_m_* determination, initial velocities in enzyme reactions (rate of CoA released in μM) were plotted against different concentration of acetyl-CoA using non-linear regression to the Michaelis-Menten equation using Prism software (GraphPad software, version 8.0). The substrate specificity of MetA was also determined by performing enzymatic reactions in the presence of 1.0 mM L-homoserine or L-serine or L-threonine. The enzymatic reactions were also performed using 1.0 mM L-homoserine in the presence of either 1.0 mM acetyl-CoA or propionyl-CoA or succinyl-CoA.

### High-throughput screening to identify MetA inhibitors

An endpoint HSAT activity based assays was performed using the 2300 structurally diverse compounds belonging to small molecule library of the National Cancer Institute Developmental Therapeutic Program (NCI-DTP) in 96 well format. High through put screening assays were performed in 30 μl reaction volume containing 100 mM Tris-Cl, pH-7.4, 5.0 mM MgCl_2_, 500 nM (His)_6_-MetA and the corresponding compound of the chemical library at a single concentration of 100 μM. The enzyme-scaffold mix was incubated at room temperature for 10 mins and the reaction was initiated by the addition of 1.0 mM L-homoserine and 1.0 mM acetyl-CoA. The reaction was further incubated at 37°C for 10 mins, the amount of CoA released in enzymatic reaction was determined upon DTNB addition and subsequently, total activity and percentage inhibition was calculated. All reaction plates contained appropriate enzyme only, substrate only and assay buffer control. The activity assays were also performed in the presence of 3.125 μM to 100 μM concentration of various compounds. The final DMSO concentration in each reaction was 10% and each reaction was performed in triplicates.

### Characterization of the binding of DIS with MetA using ThermoFlour assays

The SYPRO Orange dye was used to monitor MetA folding in the absence or presence of DIS. The protein samples (5 μM) were heated from 20°C to 95°C with an increment of 1°C per min cycle. The increase in temperature results in protein unfolding, SYPRO Orange dye binds to unfolded protein and fluorescence intensity is measured at an excitation and emission wavelength of 372/472 nm and 570 nm, respectively. The data was acquired from step-one software and plotted to calculate melting temperature (T_m_) of MetA in the absence or presence of DIS.

### MIC_99_ determination and *in vitro* killing experiments

The antimicrobial activity of hits identified from high-throughput screening was measured as previously described ^70, 73^. Briefly, the small molecules were serially diluted 2.0-fold with a final volume of 50 μl. Subsequently early-logarithmic mycobacterial cultures (OD_600nm_∼0.2) were serially diluted 1:1000, 50 μl of diluted cultures were added to each well and plates were incubated at 37°C for 14 days. The minimum inhibitory concentration (MIC_99_) was the lowest concentration of compound that completely inhibited *Mtb* growth in 96 well plates. For *in vitro* killing experiments, early-logarithmic cultures (OD_600_ ∼ 0.2) of *M. bovis* BCG were exposed to 10x MIC_99_ concentration of drugs for different time points. Further, *in vitro* killing experiments were also performed in the absence or presence of 50 μg/ml or 200 μg/ml L-methionine or 10 mM N-acetyl-cysteine. At designated time point, 10.0-fold serial dilutions were prepared and 25 μl was plated on MB7H11 supplemented with oleic acid–ADC (OADC) at 37°C for 3-4 weeks.

### Metabolomics experiments

*M. bovis* BCG cultures (OD_600nm_ ∼ 0.4 - 0.6) were exposed to either DMSO or 10 μM DIS. At designated time points, cultures were harvested and samples were processed as previously described ^12^. Briefly, the data was acquired on the orbitrap fusion mass spectrometer equipped with heated electrospray ionization (HESI) source. Data was acquired on positive and negative mode at 120,000 resolution in MS1 mode and 30000 resolution in data-dependent MS2 scan mode. The spray voltage of 4000 and 35000 volt for positive and negative modes was used respectively. Sheath gas and auxiliary gas were set to 42 and 11 respectively. Mass scan range was 60-900 m/z, AGC (automatic gain control) target at 100000 ions and maximum injection time was 50 ms for MS and AGC target was 20000 ions and maximum injection time 60 ms for MS/MS was used. The extracted metabolites were separated on UPLC ultimate 3000 using XBridge BEH Amide column (100 x 2.1 mm i.d, 2.5 micrometre) maintained at 35°C temperature. The mobile phase A was 20 mM ammonium acetate in the water of pH 9.0 and mobile phase B was 100% acetonitrile. The elution gradient is used as follows: 0 min, 85% B, 2 mins, 85% B, 12 mins, 10% B, 14 mins, 10% B, 14.1mins, 85% B, and 16 mins, 85% B. The flow rate of 0.35 mL/min and the sample injection volume was 5 microliter. The acquired data was processed using Progenesis QI for metabolomics software using the default setting. The untargeted metabolomics workflow of Progenesis QI was used to perform retention time alignment, feature detection, deconvolution, and elemental composition prediction. Metascope plug of Progenesis QI was used for the in-house library with accurate mass, fragmentation pattern and retention time for database search. The available online polar compounds spectral library was used for further confirmation of identification. Samples from three biological replicates in duplicates were harvested at day 1, 2, 4 and 6, metabolites extracted and subjected to UPLC-MS analysis. The data shown is fold change observed relative to the untreated samples.

### Flow cytometry for ROS estimation

Reactive oxygen species production in untreated and DIS-treated *M. bovis* BCG samples was evaluated using dihydroethidium (DHE) as probe. Briefly, mid-log phase *M. bovis* BCG cultures (OD_600nm_ ∼ 1) were diluted to an OD_600nm_ ∼ 0.02 and treated with either DMSO (vehicle control), 1x MIC_99_ or 10x MIC_99_ DIS concentration alone or in the presence of 10 mM NAC. At different timepoints, the cultures were harvested, washed twice with 1x PBS (pH 7.4) and stained with DHE (5 µM) at 37°C for 30 mins. The fluorescence intensity was detected by FACS calibur (BD Biosciences) and analyzed by FlowJo software v10.

### Bacterial RNA-seq and analysis

To assess the effect of DIS on *Mtb* transcriptome, early log phase cultures (OD_600nm_ ∼ 0.2) and treated with 1x- and 10x-MIC_99_ concentration of DIS for 24 h. The samples were then processed for total RNA isolation by TRIzol reagent as per manufacturer’s protocol. *Mtb* total RNAs were analyzed using an Agilent Bioanalyser for quality assessment with a RNA Integrity Number (RIN) range of 6.2 to 9.7 and a median of 8.1. Ribosomal RNA (rRNA) were depleted from 500ng of bacterial RNA using RiboMinus^™^ Bacteria Transcriptome Isolation Kit (Invitrogen Thermo Fisher Scientific), according to manufacturer’s protocol. cDNA libraries were prepared from rRNA depleted mRNA using Lexogen SENSE Total RNA-Seq Library Prep Kit according to manufacturer’s protocol. The length distribution of the cDNA libraries was monitored using a DNA High Sensitivity Reagent Kit on the Perkin Elmer Labchip. All samples were subjected to an indexed paired-end sequencing run of 2 x 151 cycles on an Illumina HiSeq 4000 system. FASTQ files were mapped to the *Mtb* strain H37Rv (GenBank accession number AL123456.3) using bowtie2. Gene counts were then counted using feature Counts (part of Subread package) using annotations from same GenBank record. The DEG comparisons between 1x MIC_99_ DIS vs DMSO and 10x MIC_99_ DIS vs DMSO were performed using DESeq2. Heat maps and volcano plots were generated in R version 3.3.3. The gene type annotations from the GenBank record was used to classify the genes for analysis.

### THP-1 cell culture

Human monocyte THP-1 cells (ATCC^®^, TIB-202^™^) were cultured in RPMI 1640, complete with the following supplements; 10% heat-inactivated fetal bovine serum (FBS), 1% sodium pyruvate, 1% L-glutamine, 1% non-essential amino acids, and 1% kanamycin, at 37°C in a 5% CO2 humidified atmosphere. During infection, cells were seeded at a density of 3 x 10^6^ per millilitre and no antibiotic was used.

### Isolation of human primary monocytes

Peripheral blood mononuclear cells (PBMCs) were isolated from total blood using a Ficoll gradient. Monocytes were enriched from PBMCs using CD14 magnetic selection. Blood collection from healthy volunteers was approved by the Institutional Review Board (IRB), NHG Domain, Singapore.

### Infection of cells with mycobacteria

For infection studies, *Mtb* H37Rv and *M, bovis* BCG were grown till an OD_600nm_ 0.4 – 0.5. The mycobacterial cells were pelleted, resuspended in fresh Middlebrook 7H9 medium with 20% glycerol, and stored at −80°C. A vial of the bacterial stock was thawed to estimate CFU per millilitre. Frozen mycobacterial strains were thawed, washed, resuspended in antibiotic-free complete RPMI 1640, and used to infect cells with a multiplicity of infection of 1:5. The infected cells were incubated at 37°C in 5% CO_2_ for 3 hours, subsequently washed twice with antibiotic-free medium, counted and seeded in triplicate or replicate of five, either left treated (by different doses of INH and DIS either alone or in combination) or untreated. At designated time points, treated cells were washed once with 1x PBS, lysed with 0.06% SDS and 10.0-fold serial dilutions were plated on Middlebrook 7H11 agar supplemented with OADC at 37°C for 3-4 weeks.

### Mouse models of infection with *Mtb*

The *in vivo* anti-tubercular activity of DIS was evaluated in an acute infection model of C57BL/6 mice. Adjunctive activity of DIS with INH was evaluated in a chronic infection model of C57BL/6 mice. Female C57BL/6 mice (6 to 8 weeks old), were anesthetized then infected intratracheally with *Mtb* at a dose of ∼1000 CFU. Three to four animals were sacrificed at day 1 to determine the number of bacteria embedded in the lungs, which is 150-300 CFU. For the acute model, DIS treatment was initiated 7 days after infection, and mice were orally administered with the drug once a day, 6 days a week, for 4 wk. Two treatment groups: (i) infected mice receiving DIS (160 mg/kg), diluted in poly-ethylene glycol (PEG); (ii) infected mice receiving vehicle only (PEG). For the chronic model, treatment was initiated 30 days after infection, and the drugs were orally administered once a day, 6 days a week, for 4 wk. Three treatment groups: (i) infected mice receiving INH (10 mg/kg), diluted in PEG; (ii) infected mice receiving INH (10 mg/kg) and DIS (160 mg/kg), diluted in PEG; and (iii) Infected mice receiving vehicle only (PEG). All mice were housed in a biosafety level 3 (BSL3) laboratory, and treated humanely using procedures described in animal care protocols. The proposed animal experiments were approved by the institutional animal ethics committee of Defence Science Organization (DSO), Singapore.

### Enumeration of *Mtb* CFU in infected mice

The mycobacterial loads in lungs and spleens of infected mice at different time points was quantified by plating tissue homogenates on Middlebrook 7H11 agar supplemented with OADC. At scheduled time points, mice were euthanized, organs were aseptically collected, washed and homogenised in 1x PBS using a MACS tissue dissociator. Tissue homogenate were serially diluted and plated in triplicate on Middlebrook 7H11 agar supplemented with OADC at 37°C for 3-4 weeks. CFU obtained from two or three dilutions were used to calculate the total number of CFU per tissue per mouse.

### Histology

The lower left lobe of the mouse lung was fixed in 10% buffered formalin and paraffin-embedded. Sections were stained with haematoxylin and eosin (H&E) and Ziehl-Neelsen acid-fast stain as described previously ^46^. The histologic examination was performed after coding of the samples. The images were photographed with Olympus Microscope and captured at either 10x or 40x or 60x.

### Cytokine and Chemokine Analysis

Lung homogenates from various groups were assayed by Luminex 100 using Exponent 3.2 software and Milliplex MAP for Luminex xMAP Technology Assay (MCYTOMAG-70K Mouse Cytokine 32-Plex) as per manufacturers’ instructions.

### Lung RNA-seq and analysis

Total RNA from lung tissue samples from different groups was extracted following the double extraction protocol: RNA isolation by acid guanidinium thiocyanate-phenol-chloroform extraction followed by a Qiagen RNeasy Micro clean-up procedure. Total RNAs were analyzed on Agilent Bioanalyser for quality assessment with RNA Integrity Number (RIN) range from 4.2 to 6.7 and median of RIN 5.1. Ribosomal RNA (rRNA) were depleted from 500ng of RNA using *Ribo*-*Zero*™ Gold (Human/Mouse/Rat) according to manufacturer’s protocol. cDNA libraries were prepared from the resultant rRNA depleted RNA using Lexogen SENSE Total RNA-Seq Library Prep Kit according to manufacturer’s protocol for degraded RNA except with 21 PCR cycles and an additional cleanup with 0.9X Ampure XP beads. The length distribution of the cDNA libraries was monitored using a DNA High Sensitivity Reagent Kit on the Perkin Elmer Labchip. All samples were subjected to an indexed paired-end sequencing run of 2×151 cycles on an Illumina HiSeq 4000 system. FASTQ files were mapped to the mouse genome build GRCm38 using STAR based on GENCODE M16 annotations. Gene counts were determined using feature counts (part of the Subread package). DESeq2 was used to identify differentially expressed genes (DEG) from the gene counts in R version 3.3.3. Infection modulated genes were defined as those whose expression was significantly altered in comparison to untreated and uninfected samples using DESeq2. Time and treatment modulated genes were determined using DESeq2 as an interaction model with the main terms being time and treatment and as interaction term time : treatment. Multiple testing correction was performed using the method of Benjamini and Hochberg. Heat maps and volcano plots were generated in R version 3.3.3.

### Flowcytometry for characterizing lung immune cells

The lung tissues of chronically infected mice at 60 days p.i were aseptically isolated and incubated at 37°C in 10% FBS RPMI media containing 1.4 mg/ml collagenase A and 30 µg/ml DNase I for 60 mins. The treated lung tissue was dissociated over the 70 µm cell strainer. Strainer was washed to collect single cell suspension. RBC were lysed with ACK lysing buffer (Lonza) for 5 mins following by washing of cells with culture media. Cells were counted and adjusted to 1 × 10^6^ cells/ml. Live dead staining was performed for 15 mins at room temperature or for 20 mins at 37°C. Non-specific antibody binding was blocked by adding anti-CD16/32 (Fc-block) (BD Biosciences). The surface markers were stained for 20 mins at 4°C with two separate antibody panels. Myeloid panel - anti-CD45-FITC, anti-CD11c-APC-Cy7, anti-MHCII-Qdot-655, anti-CD11b-Qdot-605, anti-CD64-Pacific blue, anti-F4/80-PE-Cy7, anti-Ly6C-APC, anti-CD43-BUV737, anti-CD24-PerCP and anti-CD3/CD19/Ly6G-BUV395 (Lineage mark). Lymphoid panel - anti-CD45-Pacific blue, anti-CD3-FITC, anti-CD4-PerCP, anti-NK1.1-APC, anti-CD44-Qdot-655, and anti-CD62L-PE-Texas Red. The stained cells were acquired using a LSRII cytometer with 5 lasers and flow data was analysed with FlowJo software.

### Statistics

All values are expressed as the mean ± SEM or mean ± SD of individual samples. Samples were analysed using Mann-Whitney test for two groups and Kruskal Wallis test or 2way ANOVA for multiple groups. Graphs were generated using Graph Pad prism.

## Author contributions

R.S. and A.S. conceived the idea, designed and oversaw the study. D.C. and M.M performed the experiments, and analyzed the data. A.L. and R.B. performed flow cytometry. T.P.G. and S.K. performed MIC experiments. Ava.S. and K.G.T. performed protein purification and SV-AUC experiments. F.S. and J.L. performed RNAseq studies. B.L. performed bioinformatics analysis. K.C and N.A. provided reagents. S.K.G. and Y.K performed metabolomic experiments. L.T. performed histopathological analysis. C.G., K.T. and P.K. performed experiments with the DR strains. R.S and A.S. wrote the manuscript. All authors discussed results and commented on the manuscript.

## Acknowledgements

We thank DSO, Singapore and THSTI, Infectious Disease Research Facility personnel for assistance. The research fellowship to D.C. and T.P.G. from Department of Biotechnology, Govt. of India is acknowledged. Ava.S. fellowship was supported by the University of Grants commission. R.B. acknowledges Department of Science and Technology India and FICCI for providing C.V. Raman International fellowship. R.S. acknowledge THSTI, and Department of Biotechnology, Govt. of India (BT/PR29075/BRB/10/1699/2018) for funding. This research was supported by SIgN A*STAR, A*STAR JCO-CDA grant (#1518251030), and Singapore-India Joint grant (#1518224018) to A.S.

## Supplemental Information

### Supplementary Figures

**Figure S1.**
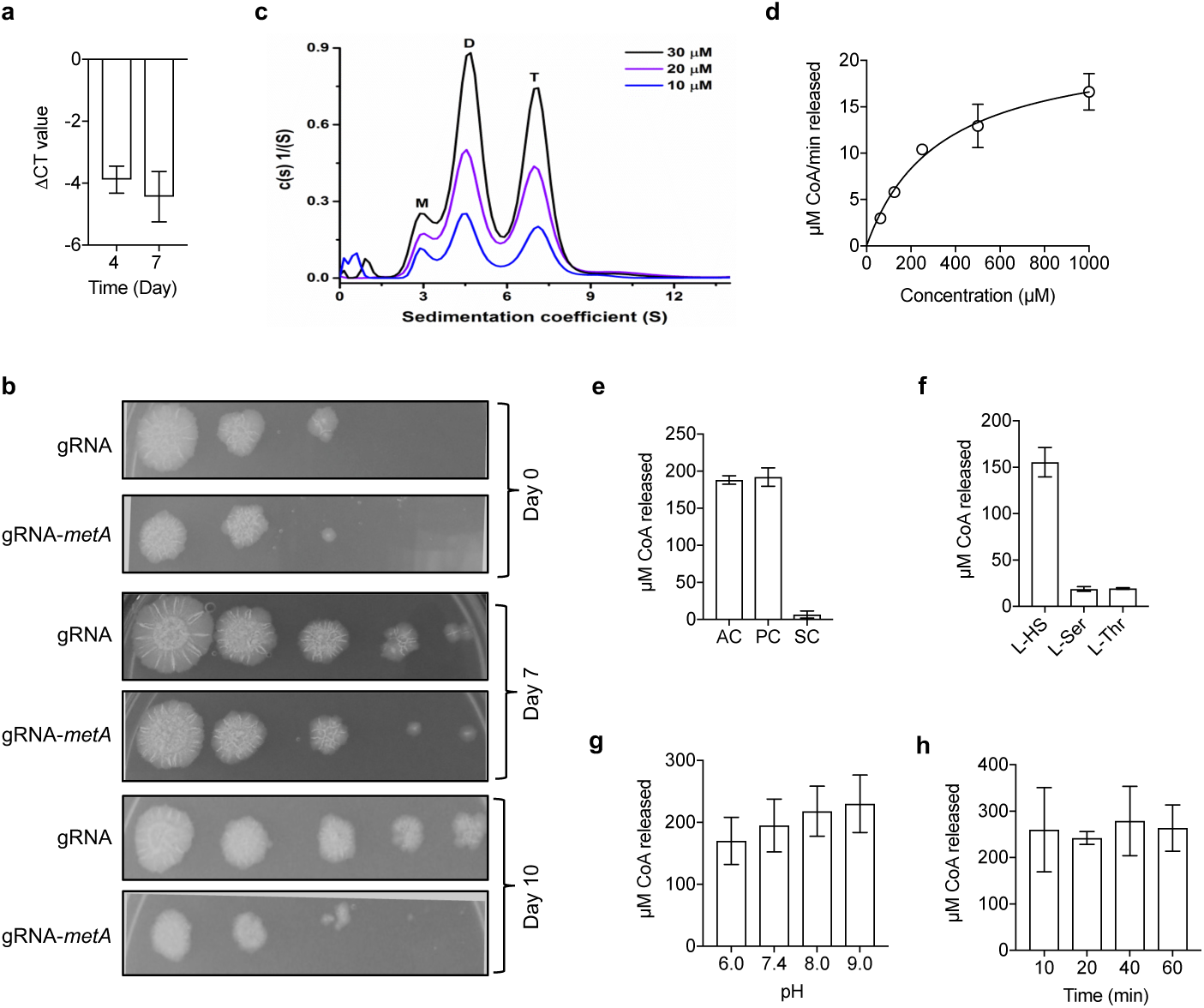
**Functional and biochemical characterization of MetA from *M. tuberculosis*. a**, Silencing of MetA expression in *Mtb* using CRISPRi approach. The expression of *metA* was quantified in knock down strains after normalization to the transcript levels of house-keeping gene, *sigA*. The data shown in this panel is representative of two different experiments. **b**, The effect of conditional repression of MetA on *in vitro* growth of *Mtb* was determined by spotting serial dilutions of various strains at different time points post-induction. The data shown in this panel is representative of two independent experiments. **c**, Oligomeric state of MetA enzyme. SV-AUC was performed using purified MetA protein at 10 μM, 20 μM and 30 μM concentration. **d**, Michaelis Menten plot for MetA enzymatic activity. The data shown on y-axis is μM CoA released in enzymatic reaction. **e**, Substrate specificity of MetA enzyme. Enzymatic reactions were performed in the presence of either 1.0 mM acetyl-CoA (AC) or propionyl-CoA (PC) or succinyl-CoA (SC). f, The specificity for L-homoserine (L-HS) was determined by performing enzymatic reactions in the presence of either 1.0 mM L-HS or L-serine (L-Ser) or L-threonine (L-Thr). **g**, Effect of buffer pH on MetA activity *in vitro*. MetA activity assays were performed in buffers of pH in the range of 6.0 – 9.0. **h**, Time kinetics analysis of MetA activity. The MetA activity was calculated after 10 mins, 20 mins, 40 mins and 60 mins post-incubation. The in panels d-h is mean ± SEM of CoA released obtained from three independent experiments performed in duplicates.

**Figure S2.**
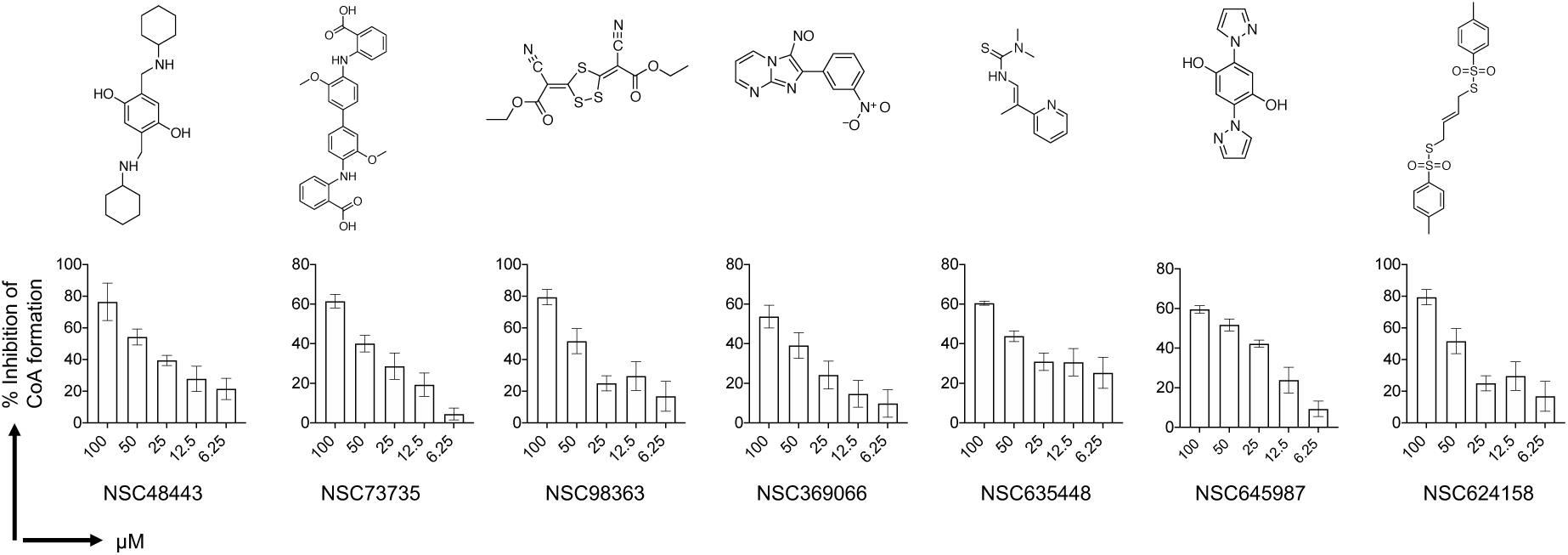
**The chemical structure of various MetA inhibitors identified from the high-throughput screening.** The enzymatic activity was performed in the presence of different concentrations of various identified preliminary hits. The y-axis depicts mean ± S.E. of percentage inhibition of enzymatic activity obtained from three independent experiments.

**Figure S3.**
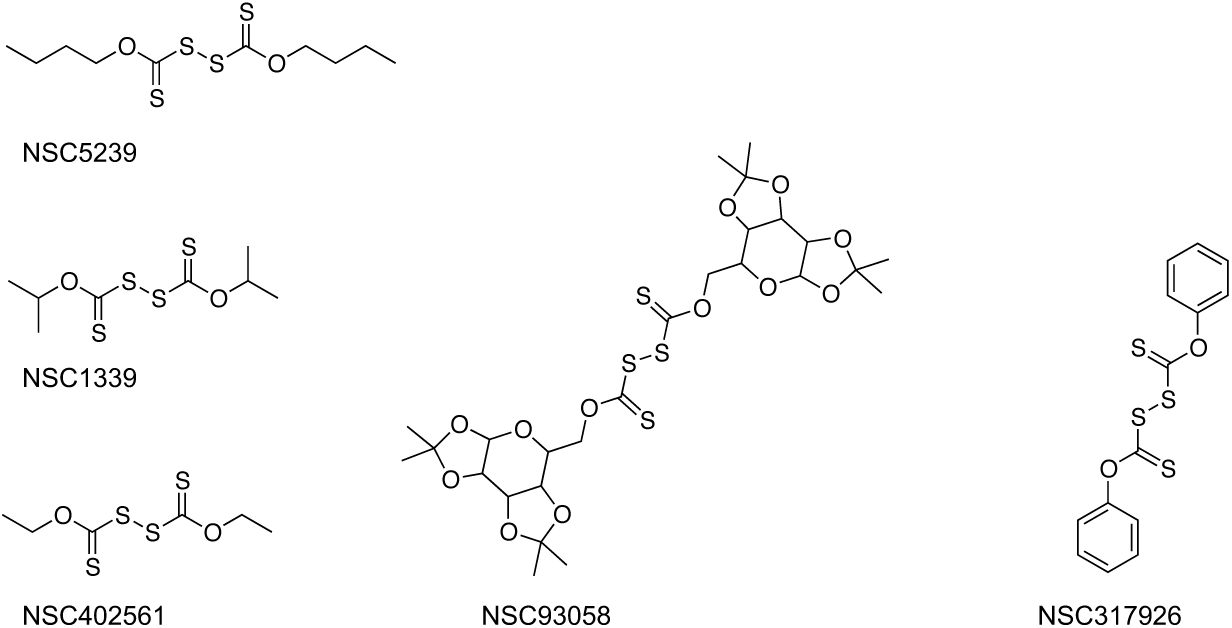
**The chemical structures of various Thiram analogs used in the study are shown.**

**Figure S4.**
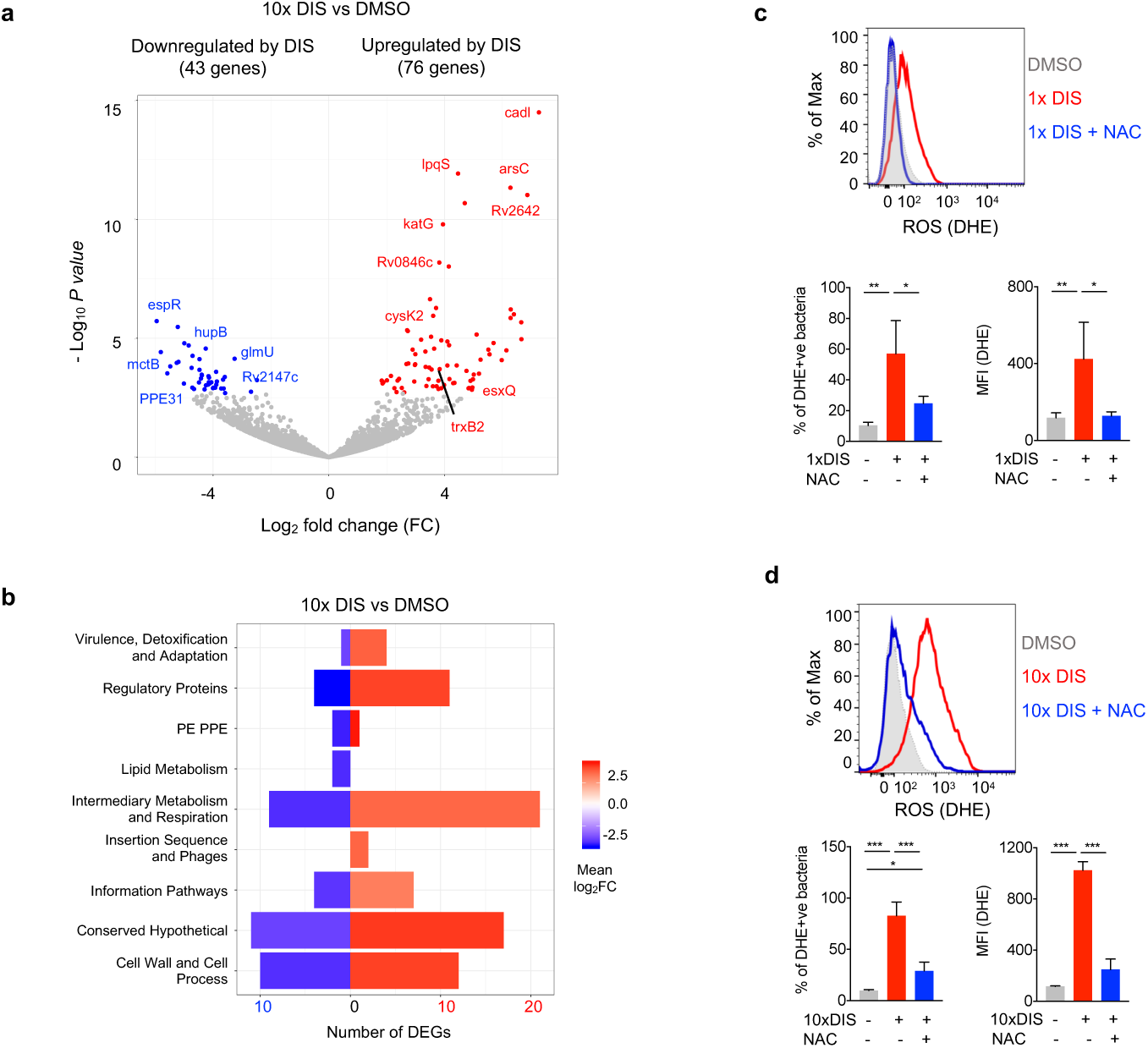
**DIS alters transcription profile and redox homeostasis of *Mtb*. a**, *Mtb* was treated with 10x MIC_99_ DIS and at 24 h total RNA was extracted. Differentially expressed genes (DEGs) upon 10x MIC_99_ DIS treatment are shown in the volcano plot (blue – downregulated, red - upregulated). b, The DEGS identified in panel A were categorized into 9 functional categories as listed in Tuberculist. The number of downregulated (43 DEGS) and upregulated (76 DEGS) transcript in each functional category are depicted as bars. The bar color is proportional to the average mean log_2_ fold change (FC) of all DEGs together for a particular functional category. **c and d**, *M. bovis* BCG cultures were exposed to different combinations and intracellular ROS was measured by flow cytometry using 5μM DHE. The data shown in panels c and d is mean ± SEM and is representative of two independent experiments. *P<0.05, ** P<0.005, *** P<0.0005 by one-way ANOVA.

**Figure S5.**
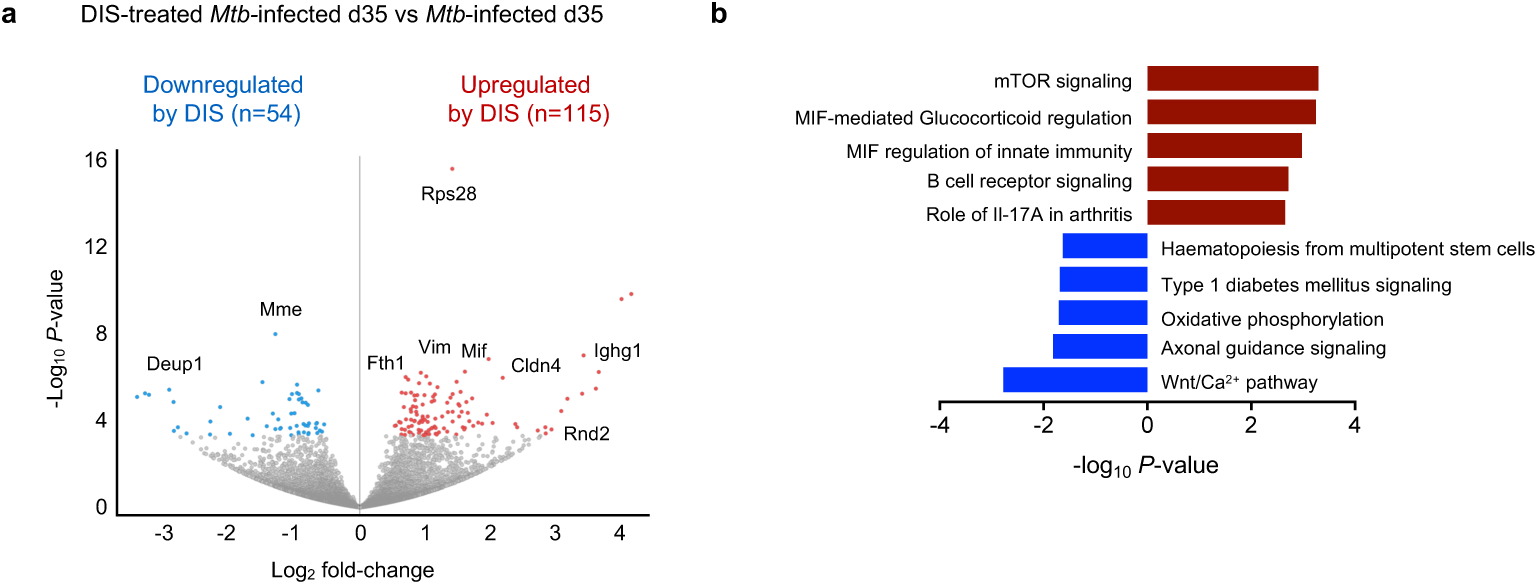
**DIS treatment alters transcription profiles of lung tissues. a,** Volcano plot of the effects of DIS at 4 weeks p.t. Significantly differentially expressed transcripts in the lung of *Mtb*-infected mice upon DIS treatment are shown in the volcano plot (blue – downregulated, red - upregulated). Genes in grey are not significant. The x axis shows the log_2_ fold changes while the y-axis show the -log_10_ nominal P-value. **b**, Ingenuity pathway analysis (IPA) of downregulated (blue bar) or upregulated (red bar) genes by DIS treatment by 4 weeks p.t. x-axis, - log_10_ *P-value* of the enriched pathway from IPA.

**Figure S6.**
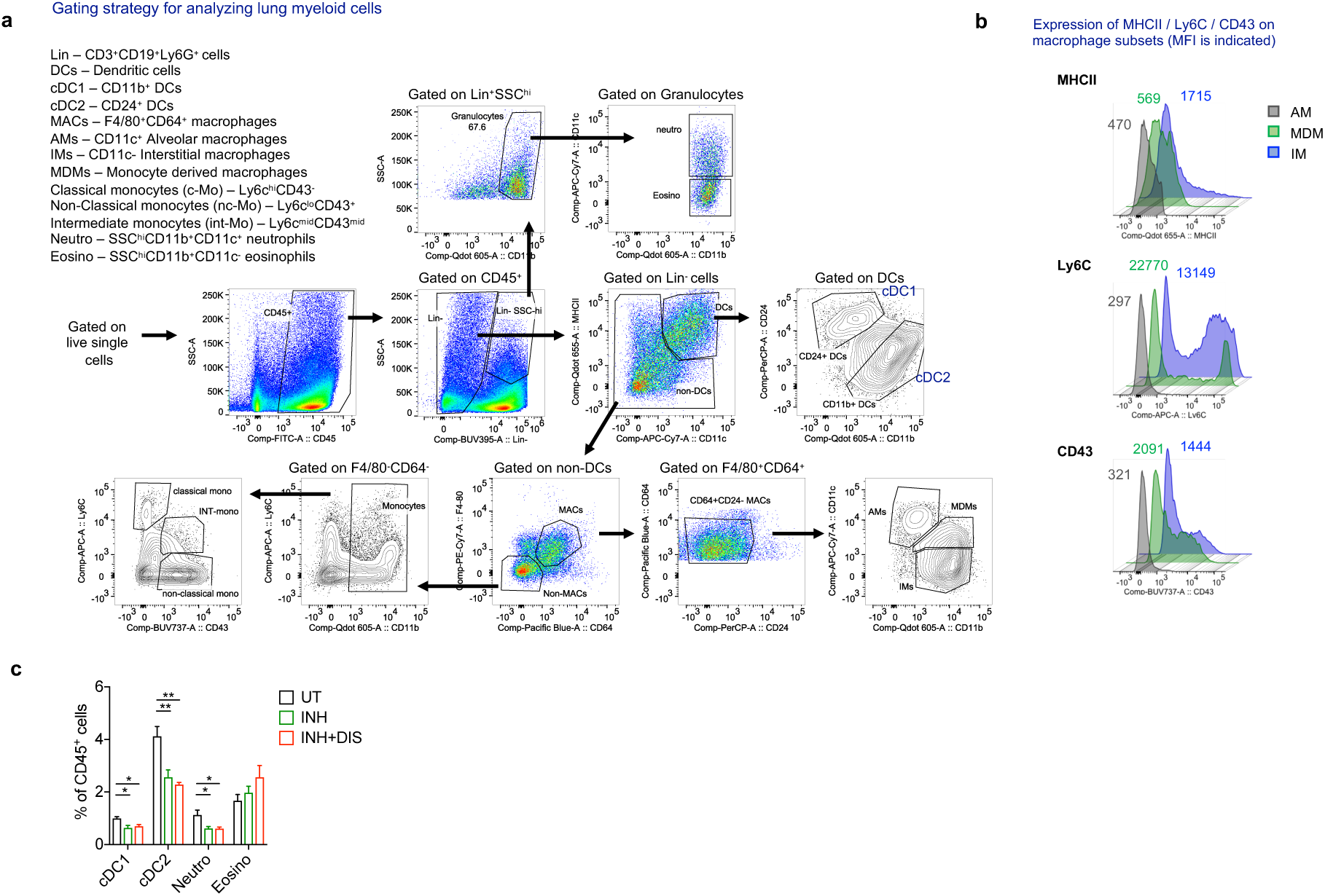
**Phenotyping of lung myeloid cells in uninfected, *Mtb*-infected and INH/DIS treated infected mice. a**, Flow cytometry gating strategy to phenotype lung myeloid cells of untreated, INH-treated or INH+DIS-treated mice at 4 wk p.t. in chronic infection model **b**, Histogram plot demonstrating expression (Median fluorescence intensity, MFI) of MHCII, Ly6C and CD43 on three different macrophage populations. AM, Alveolar macrophages; IM, interstitial macrophages; MDM, monocyte-derived macrophages. Respective MFI has been indicated. **c**, Compiled flow cytometry analysis of lung conventional type 1 dendritic cell (cDC1), conventional type 2 dendritic cell (cDC2), neutrophils (Neutro) and eosinophils (Eosino) in untreated or INH-treated or INH+DIS-treated mice. The data is mean ± SEM obtained from 4-6 mice. *P < 0.05, **P < 0.005, one-way ANOVA.

**Figure S7.**
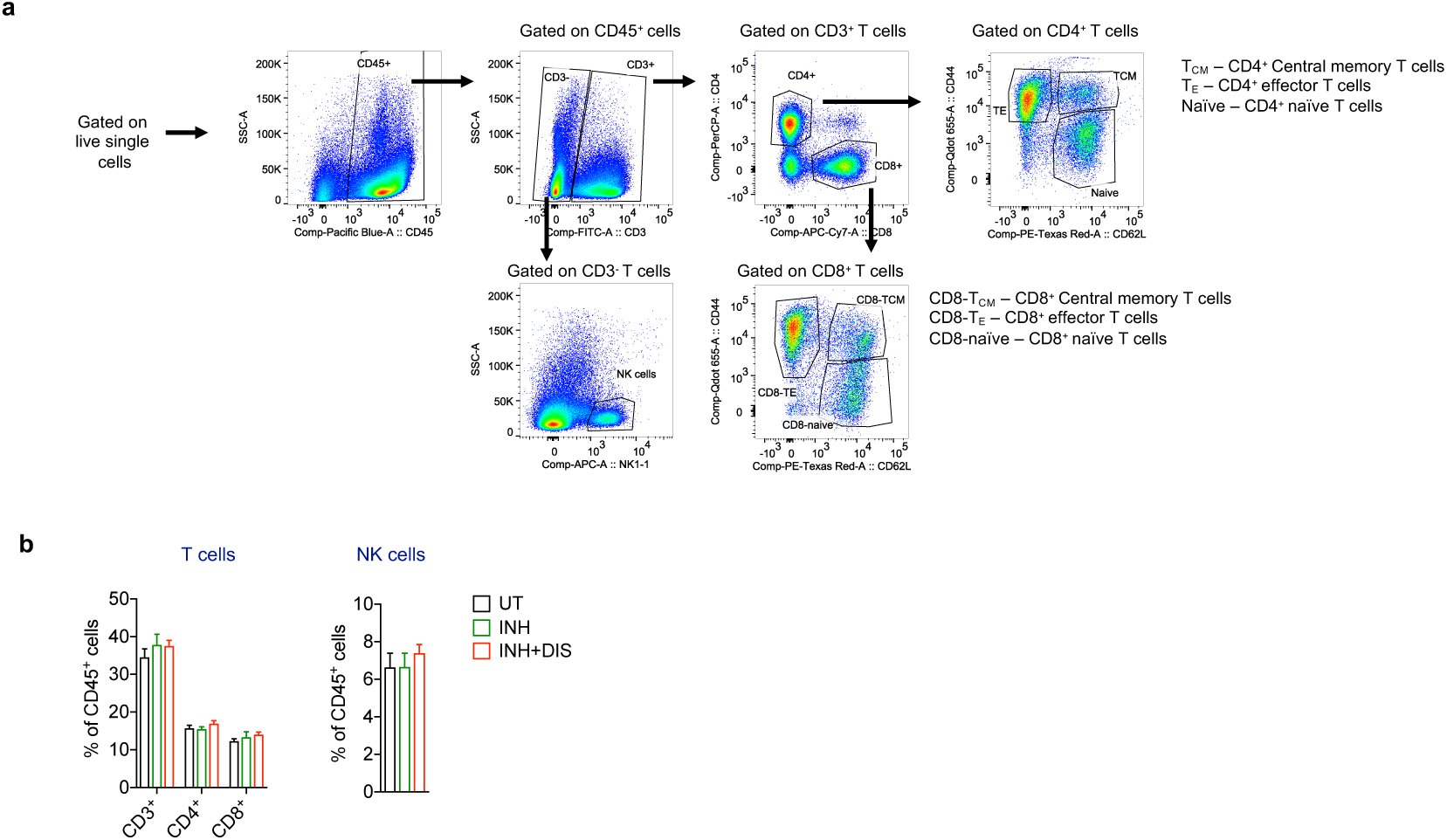
**Phenotyping of lung lymphoid cells in untreated and *Mtb*-infected INH/INH+DIS treated mice. a**, Flow cytometry gating strategy to phenotype lung T and NK cells at 4 wk p.t. in chronic infection model. **b**, Compiled flow cytometry analysis of lung CD3^+^, CD4^+^, CD8^+^ T cells and NK cells.

### Supplementary Tables

**Table S1:**
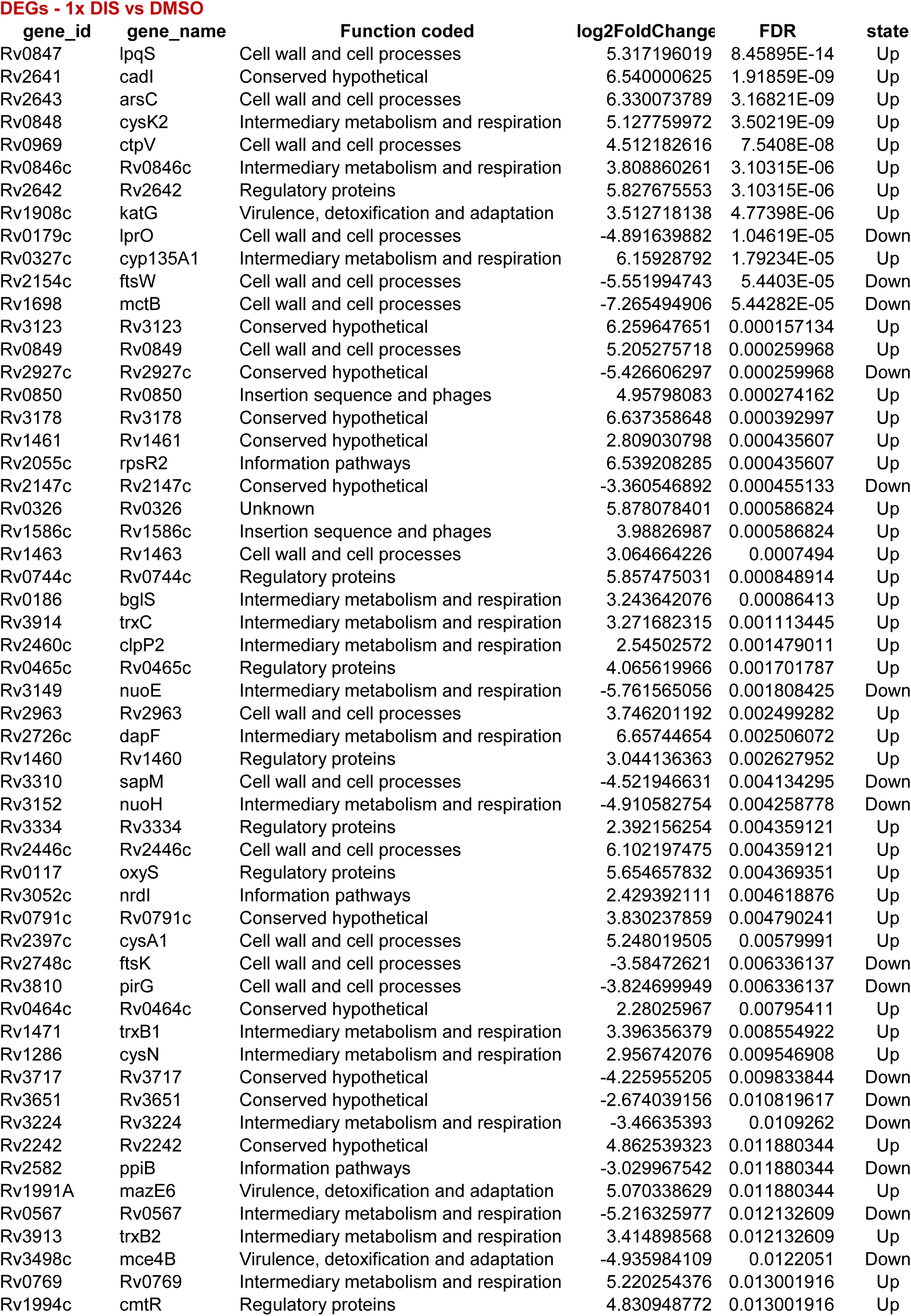
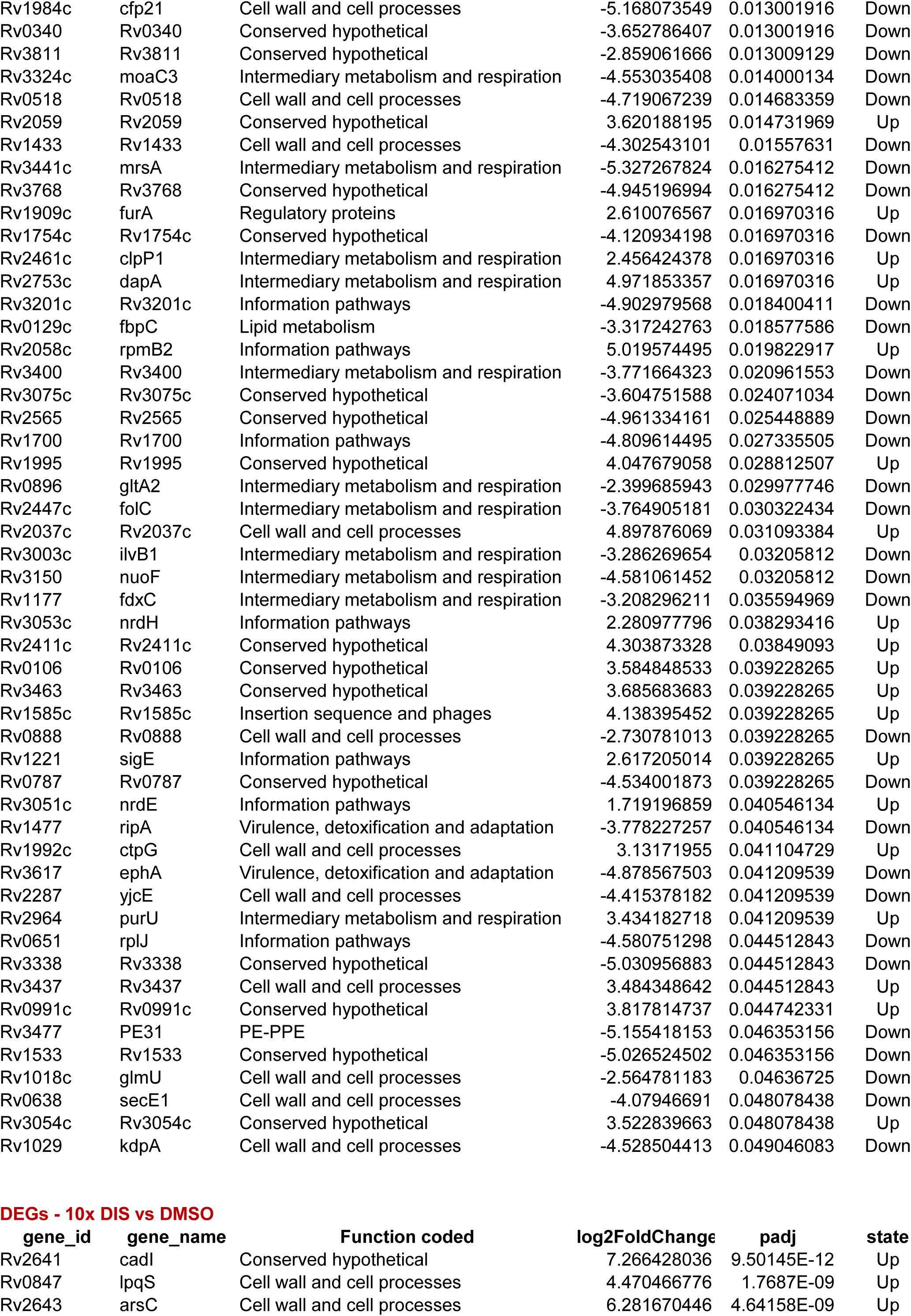
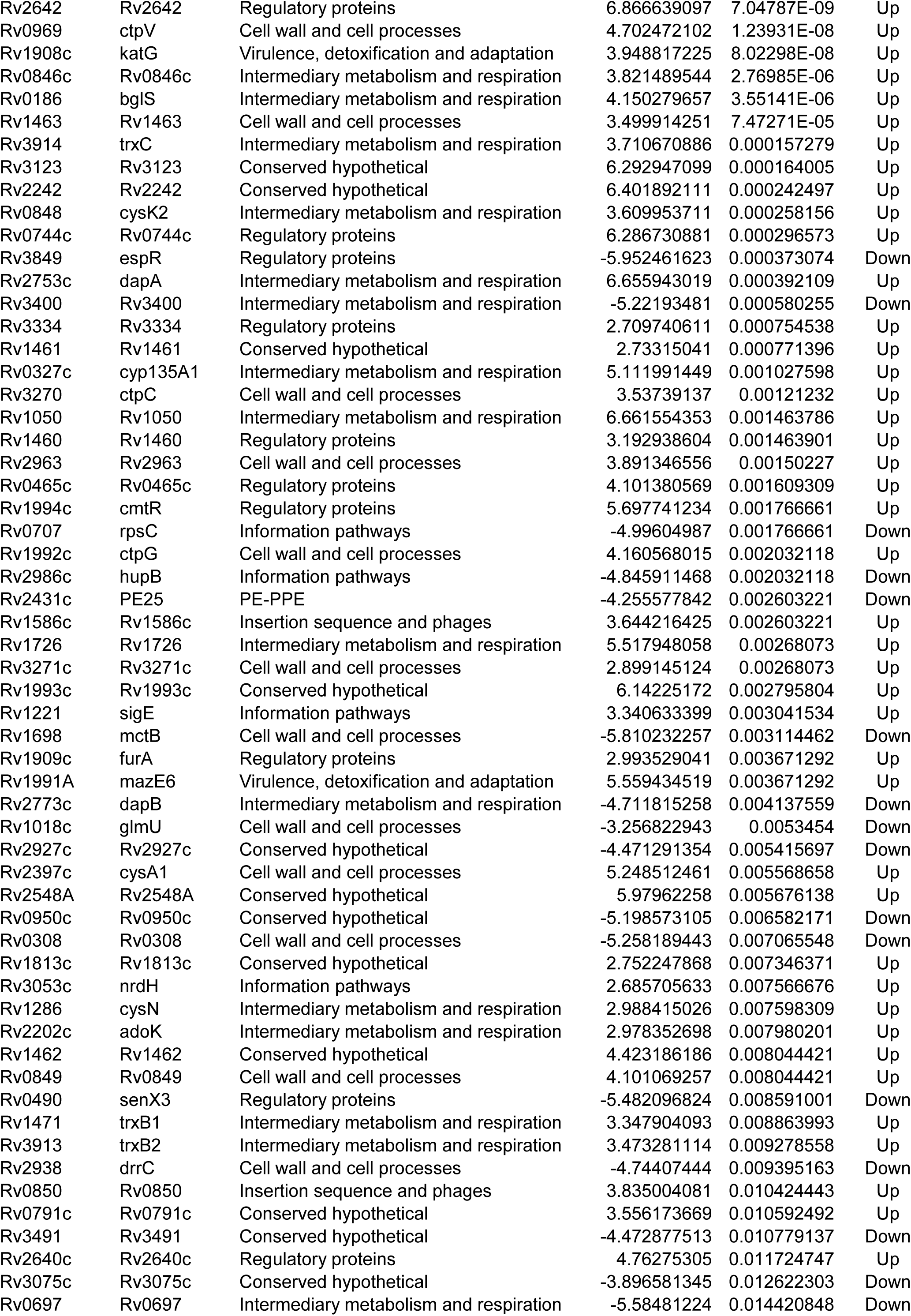
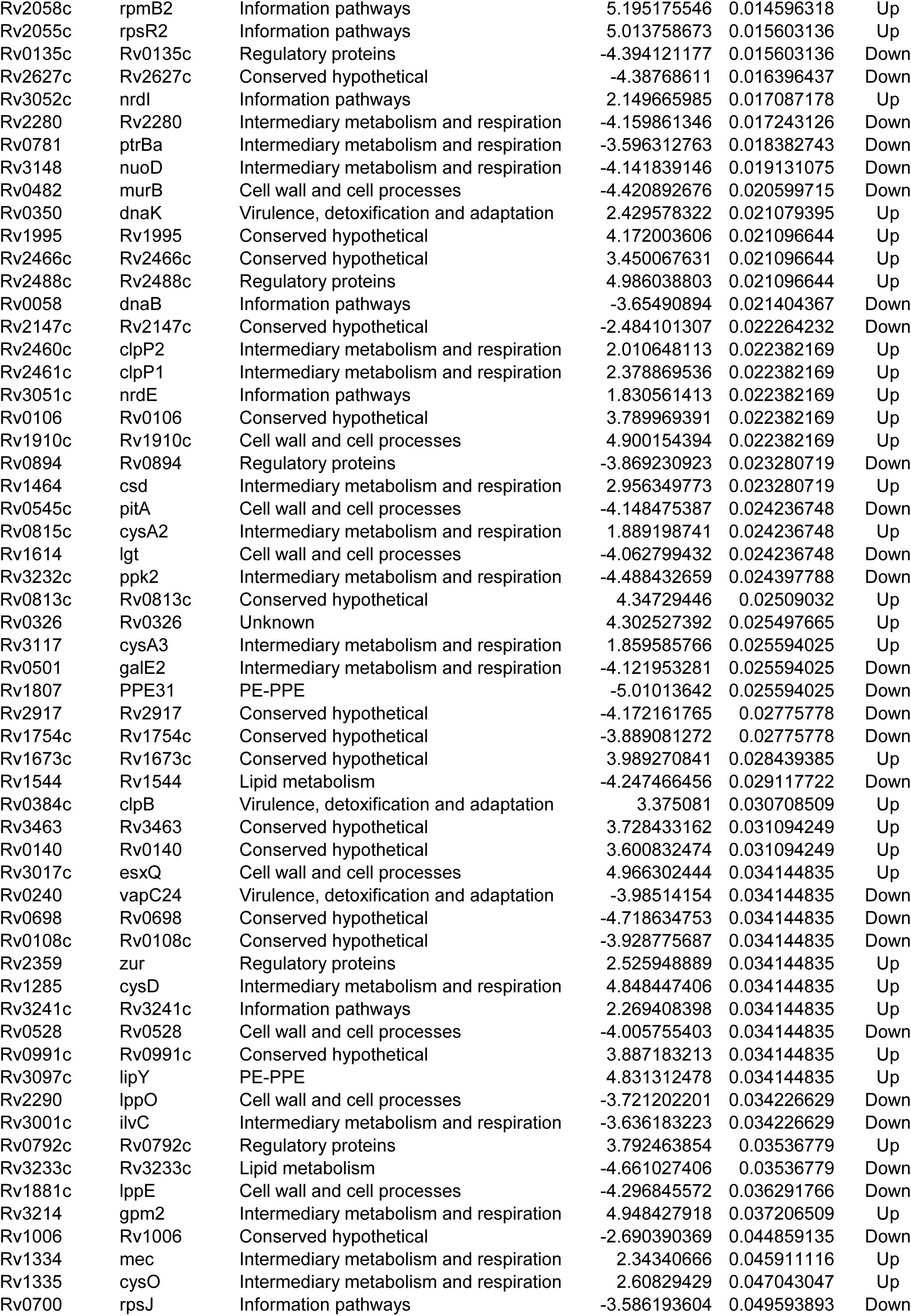
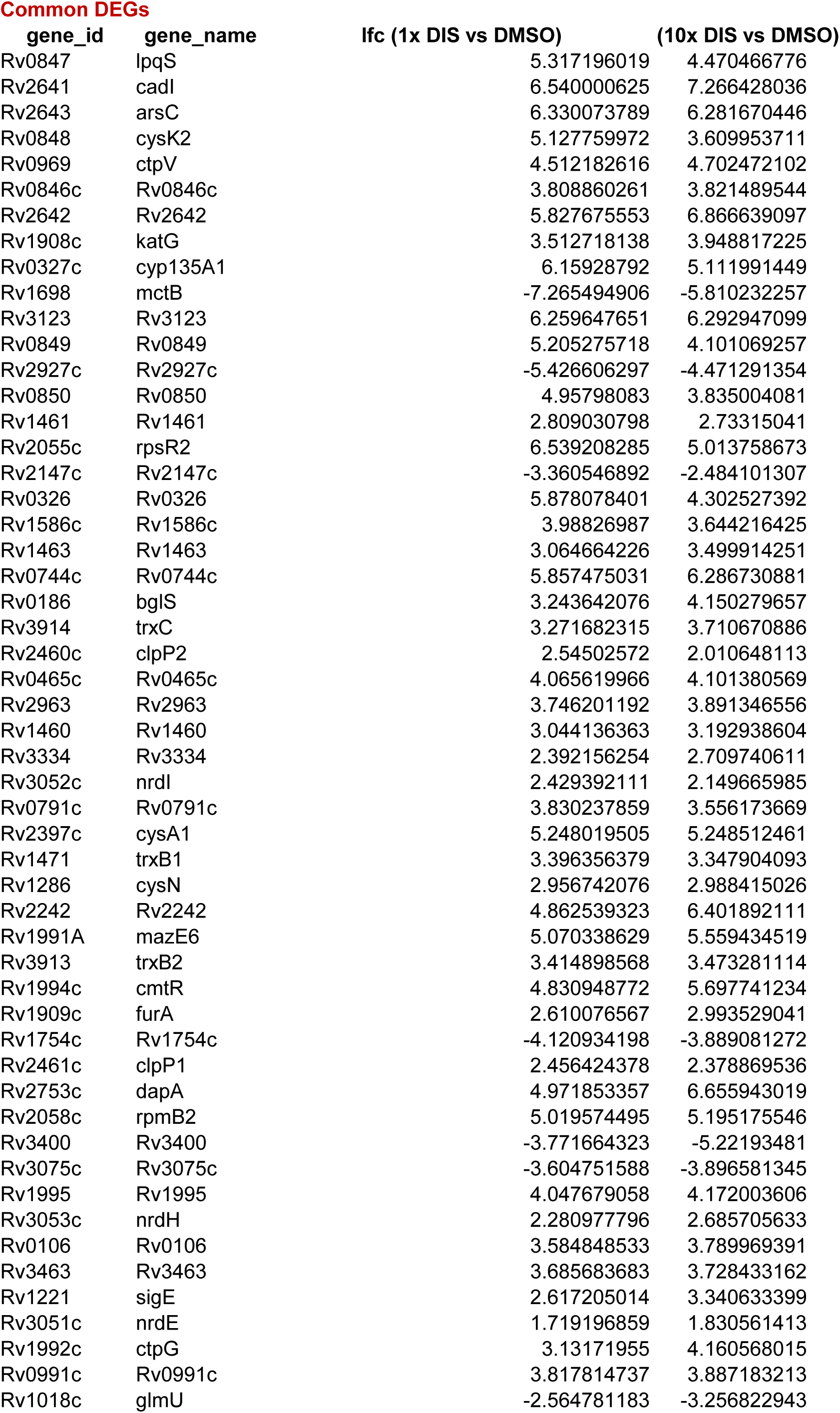
Differentially expressed genes between 1x MIC_99_ DIS, 10x MIC_99_ DIS and DMSO treated Mtb. Sheets contacting corresponding log fold changes of 1x MIC_99_ DIS vs DMSO and 10x MIC_99_ DIS vs DMSO DEGs is shown. 53 common DEGs are also indicated.

**Table S2:**
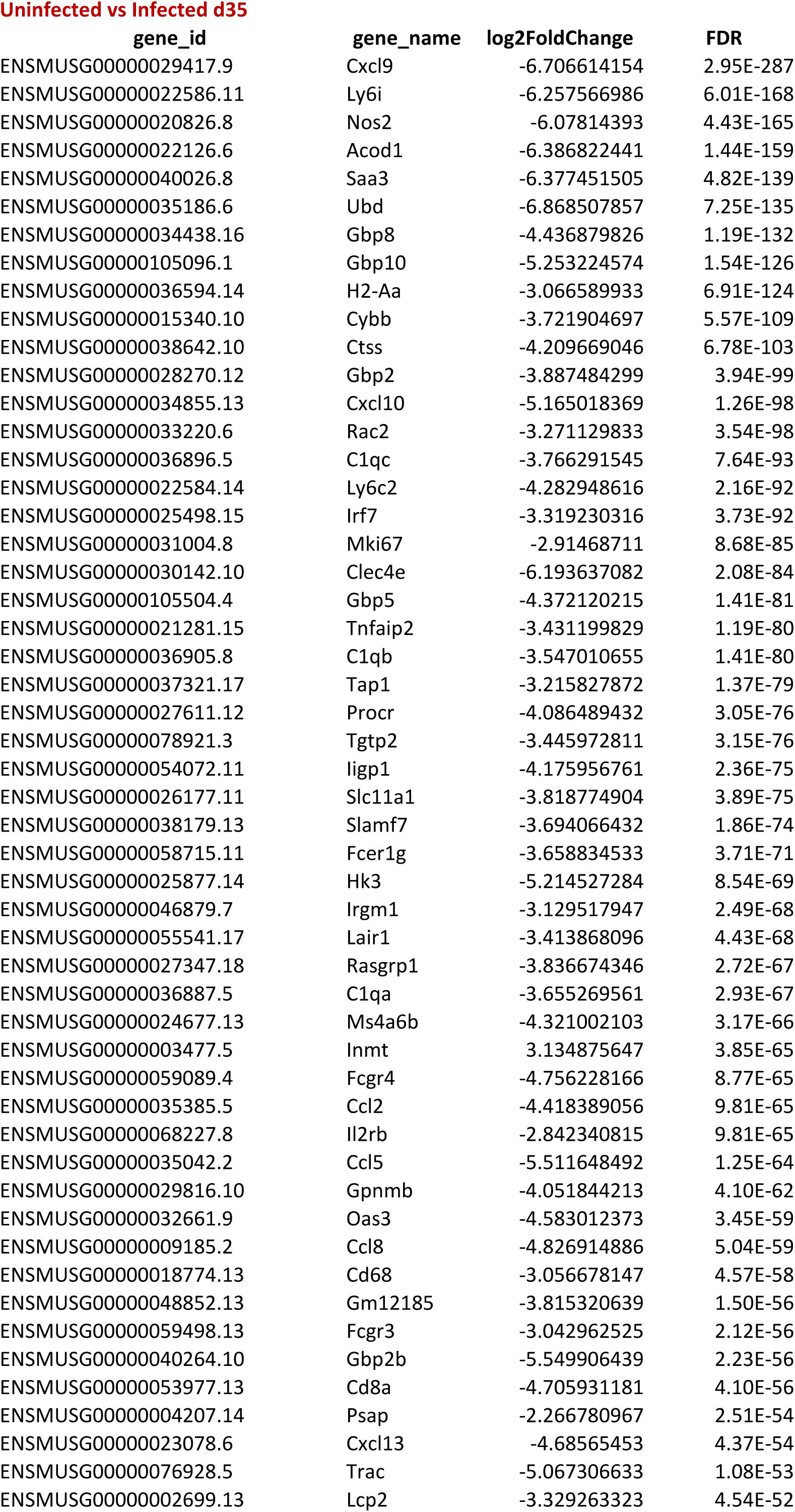
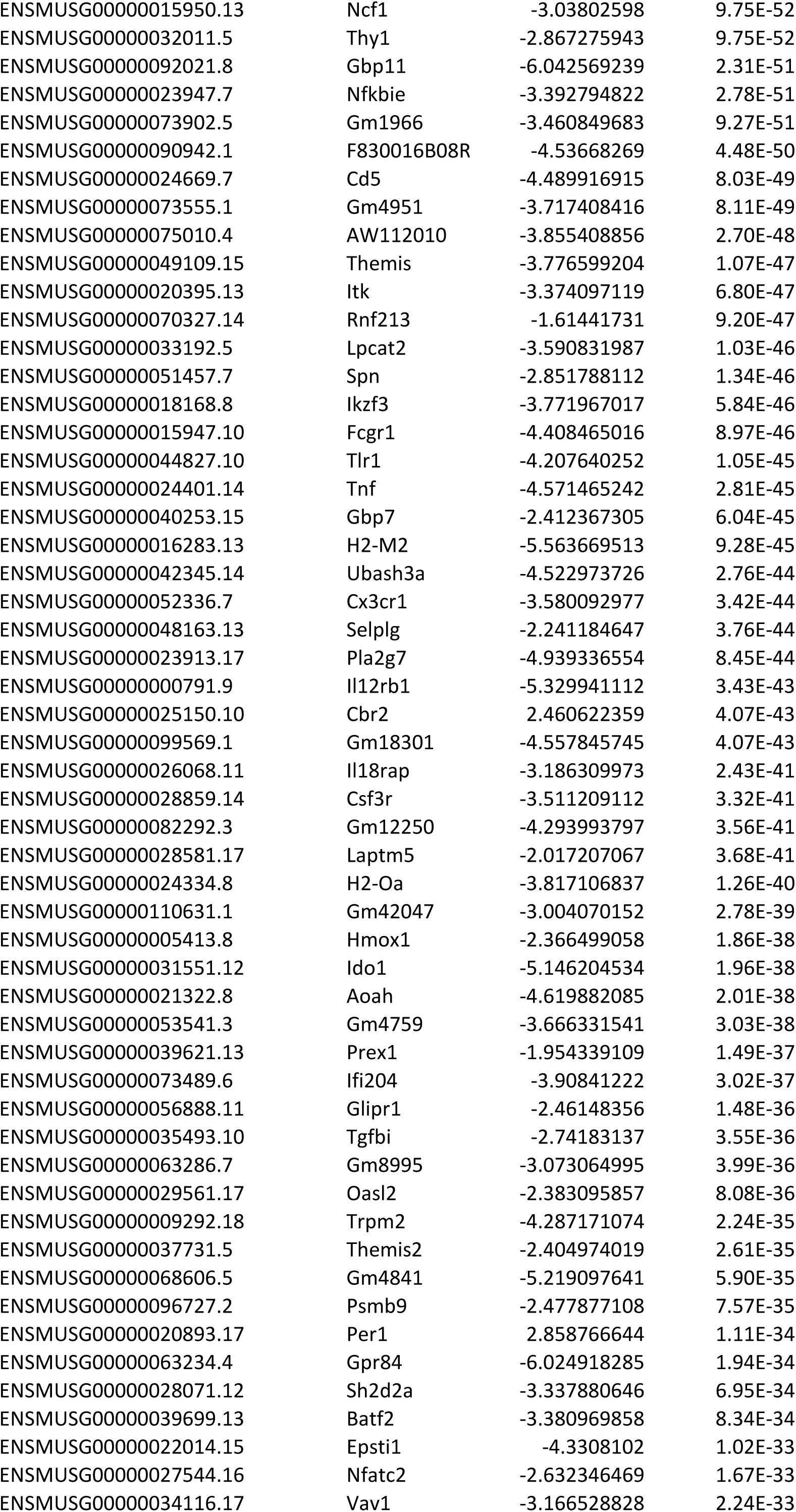
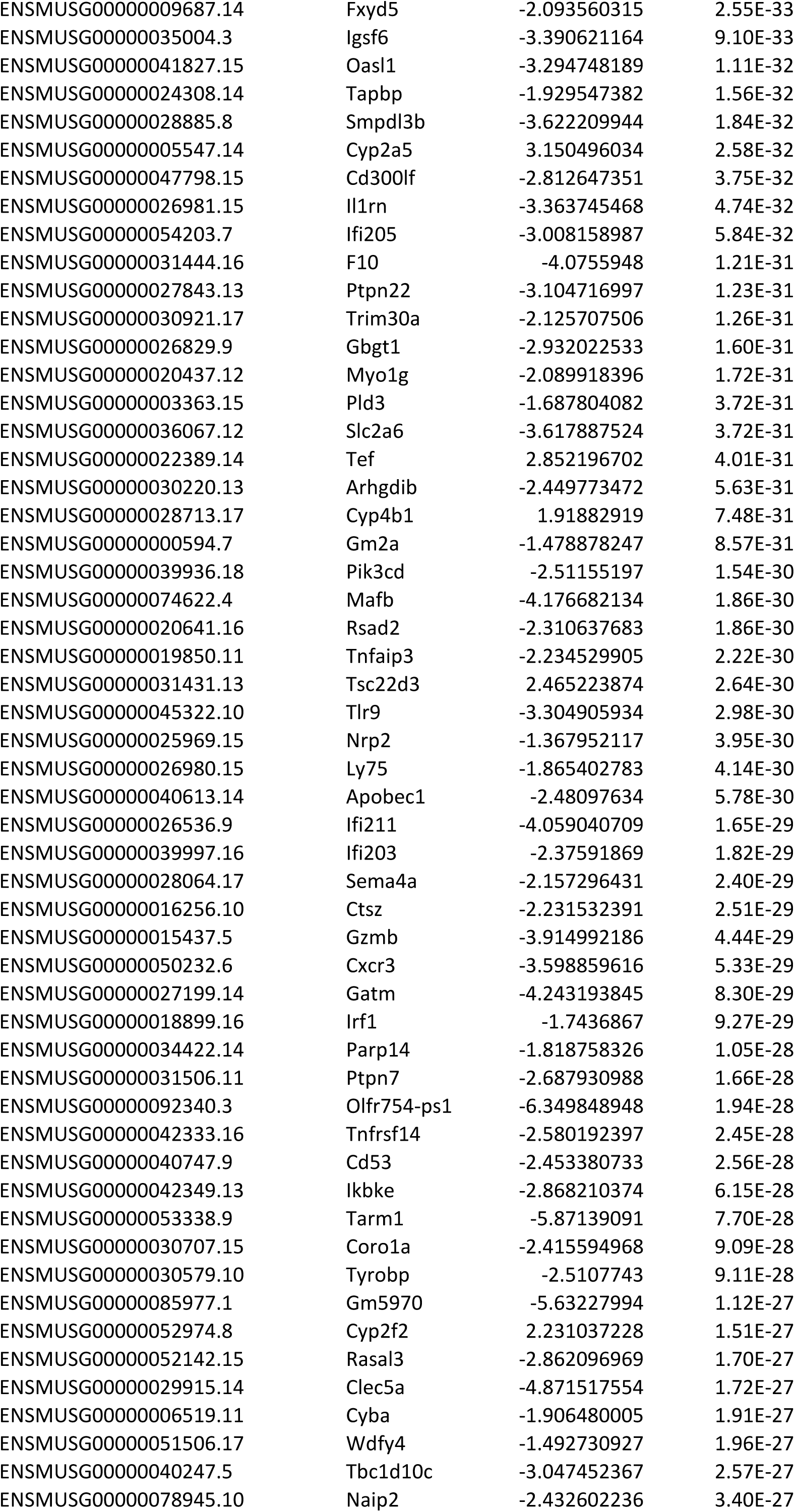
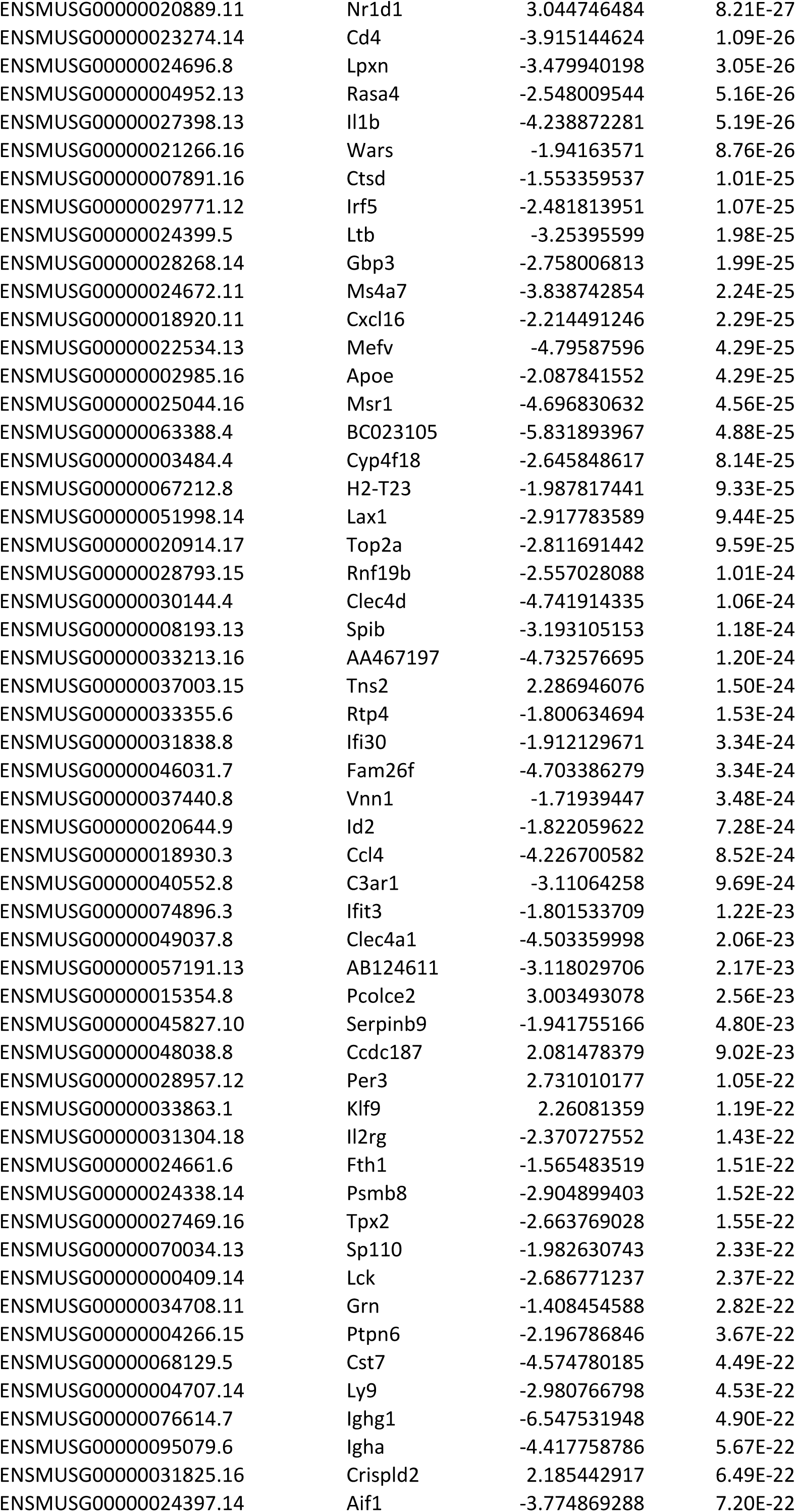
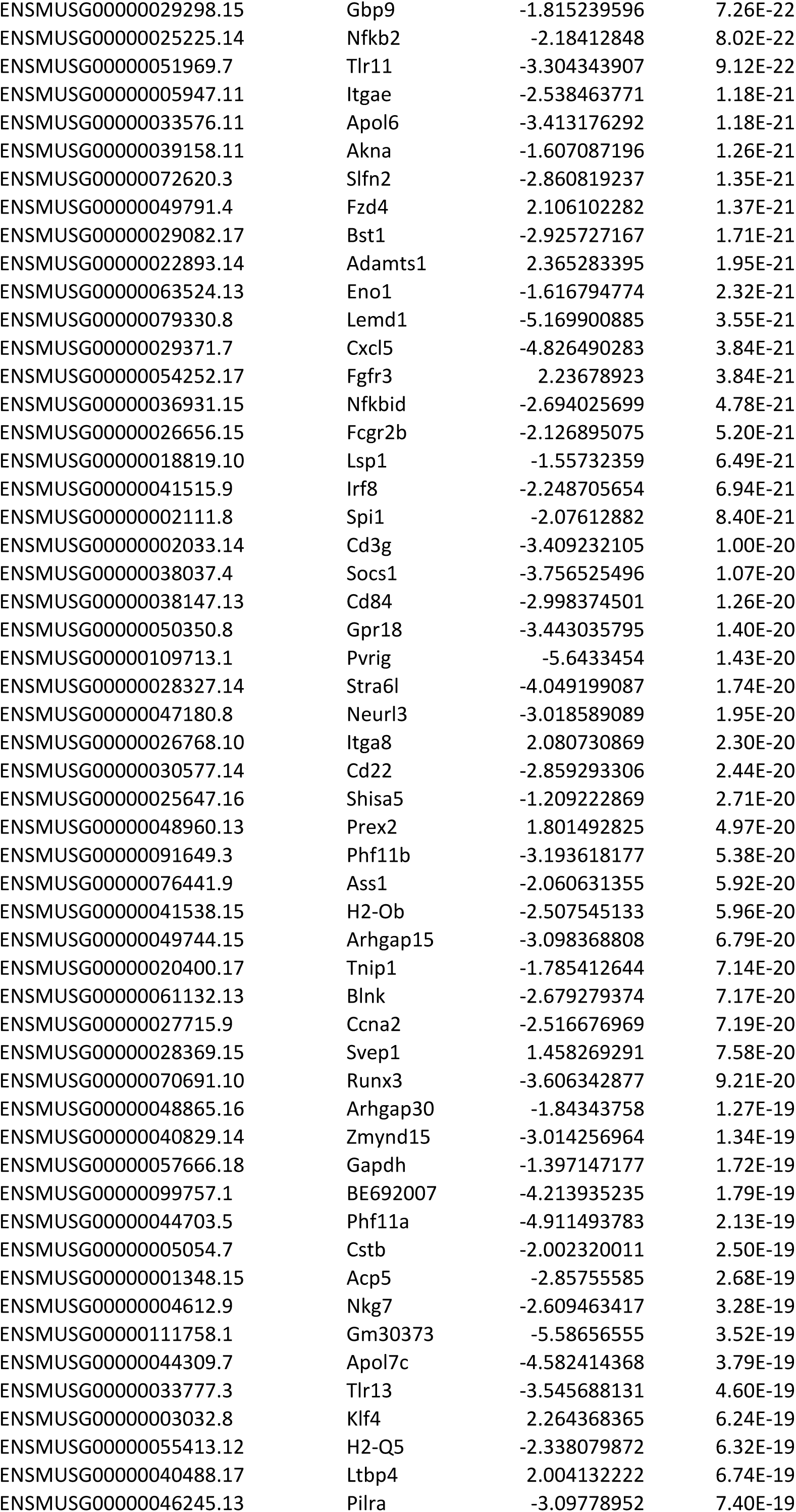
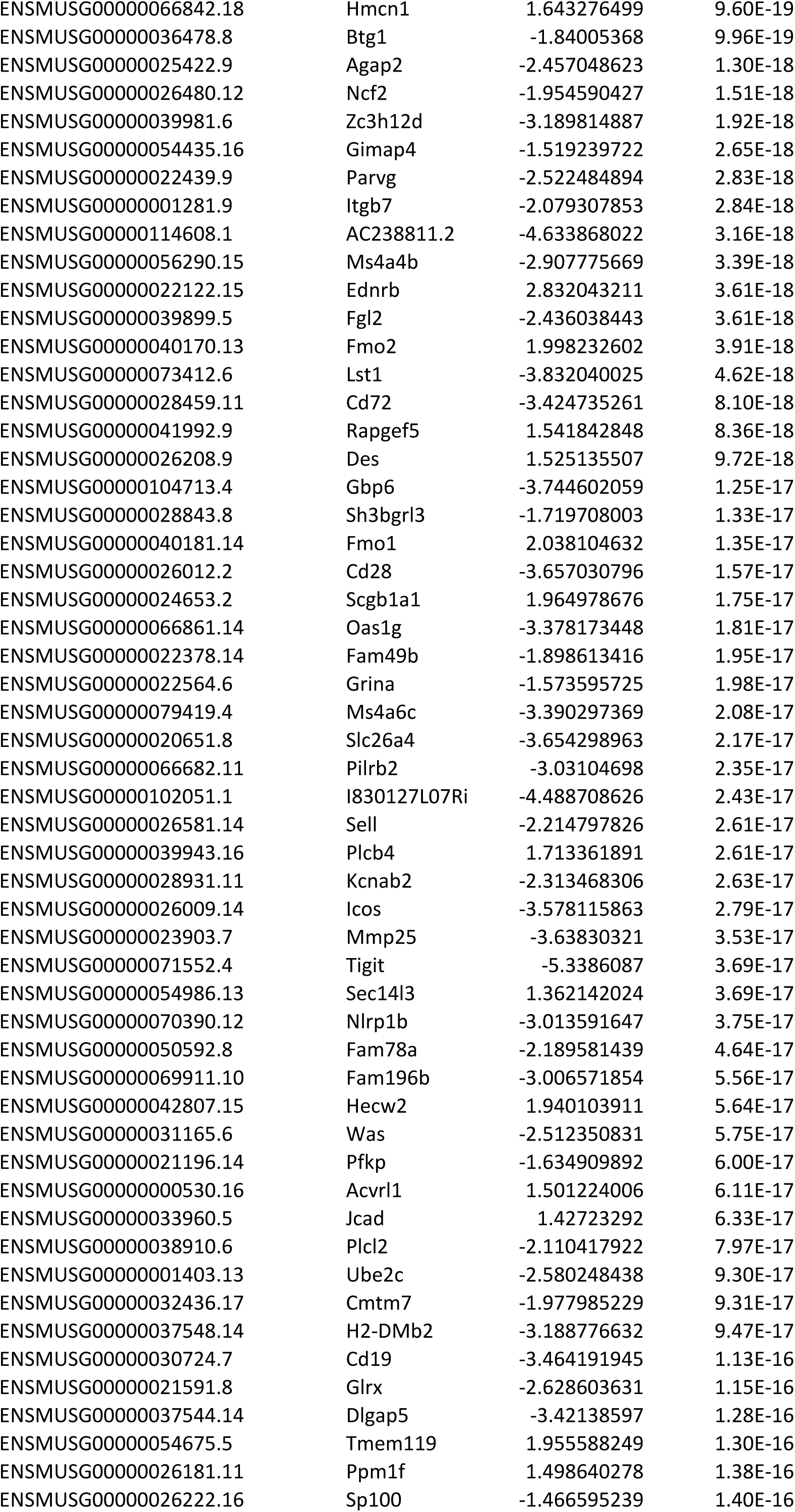
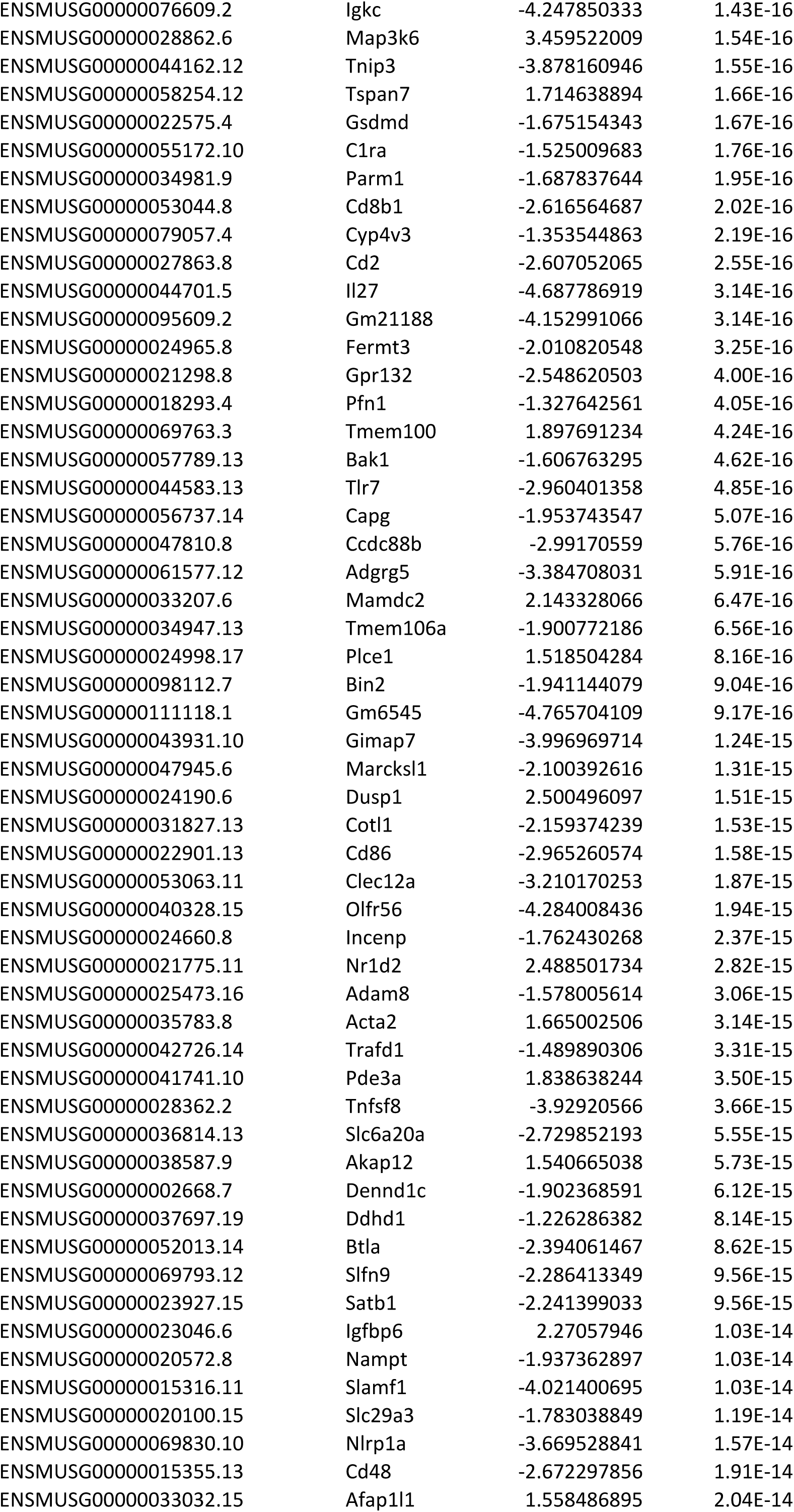
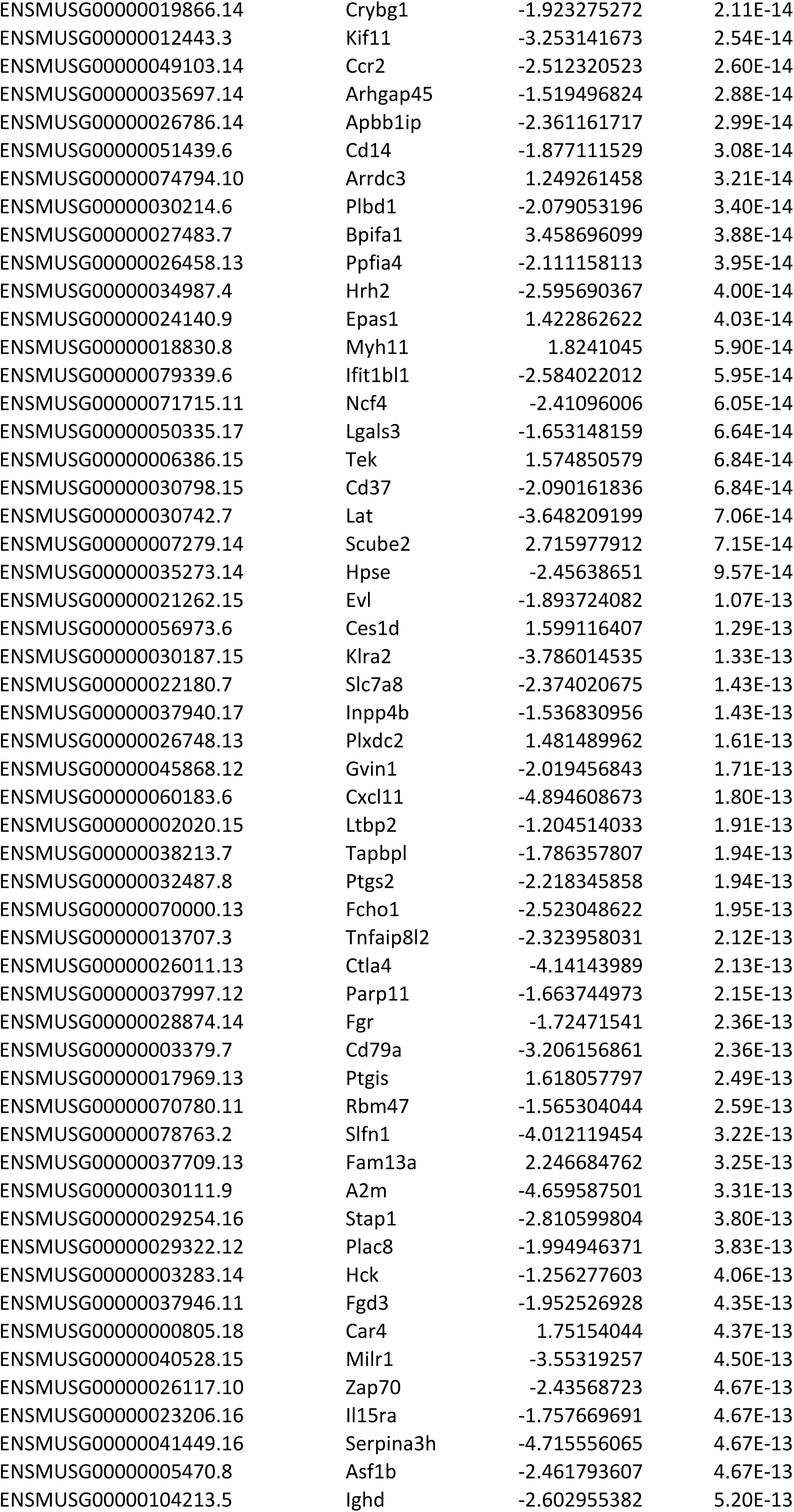
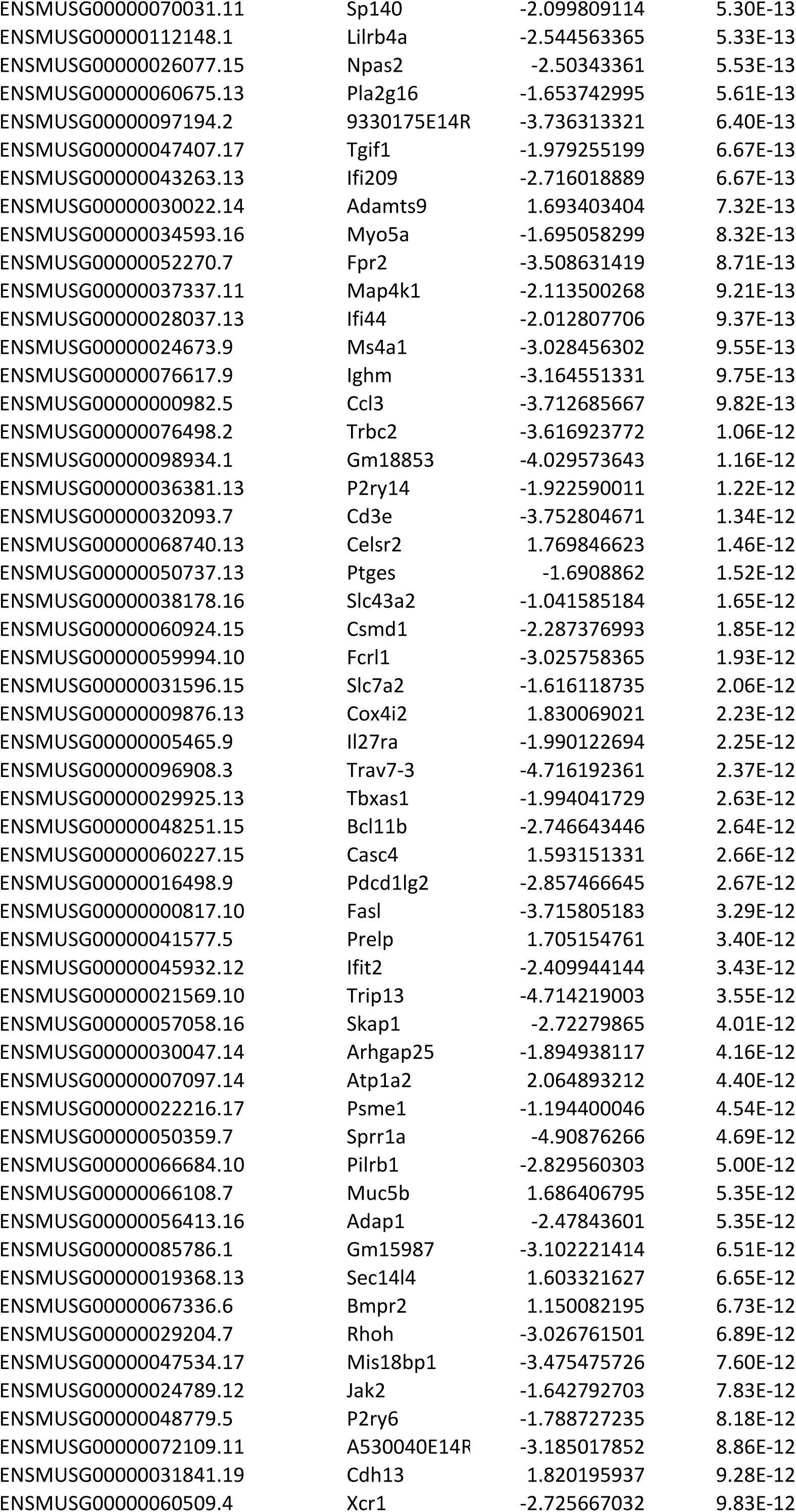
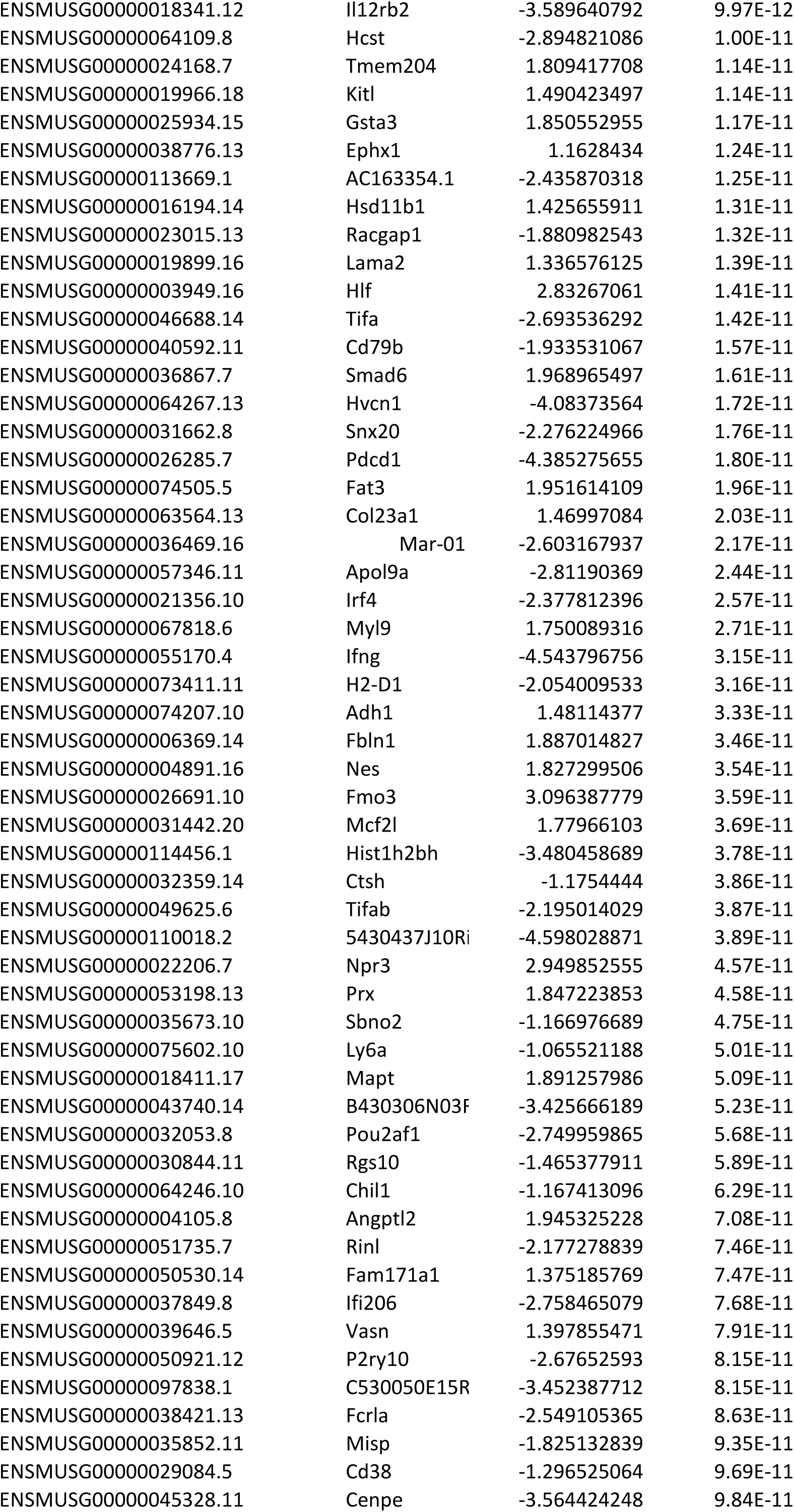
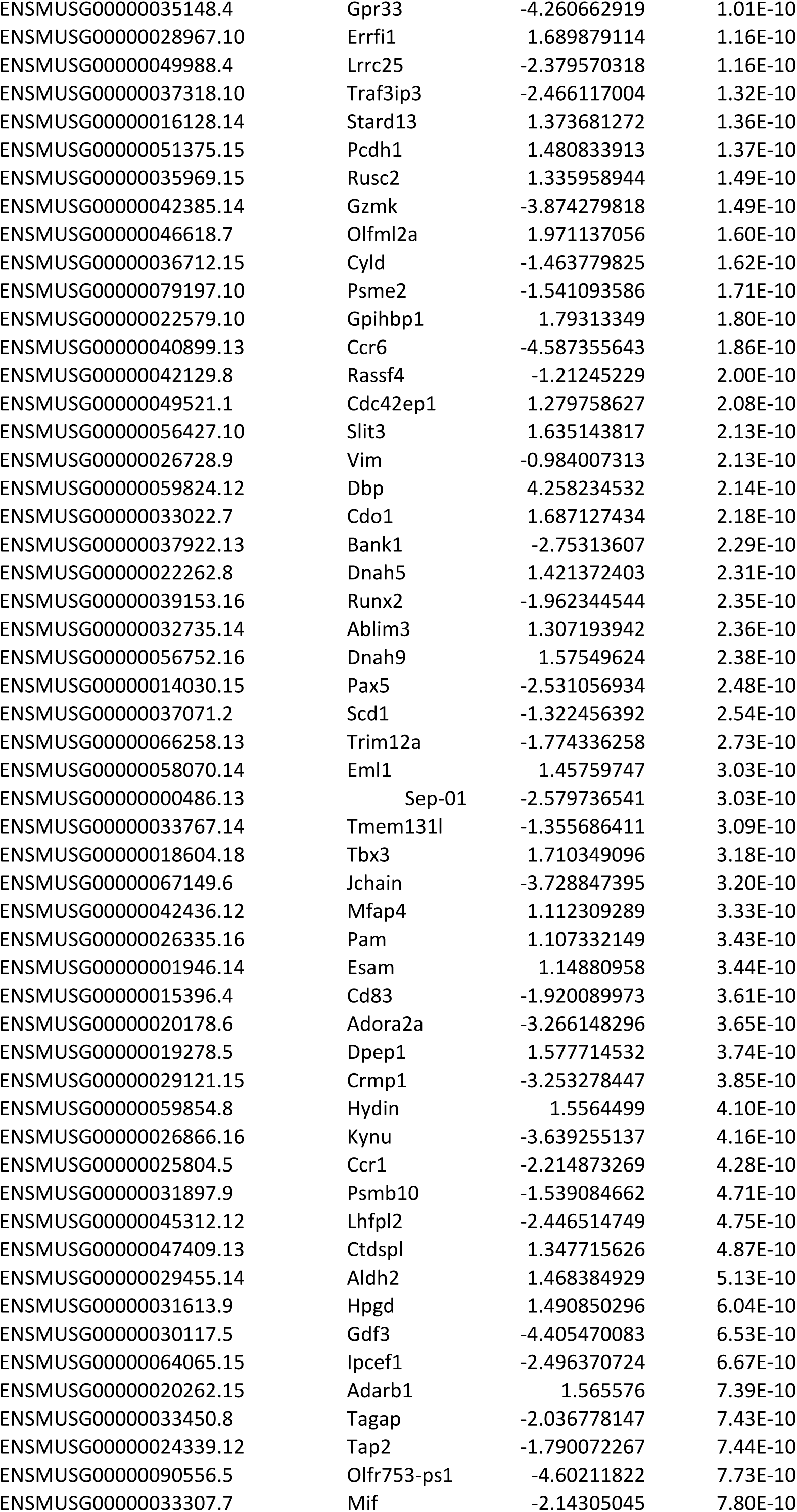
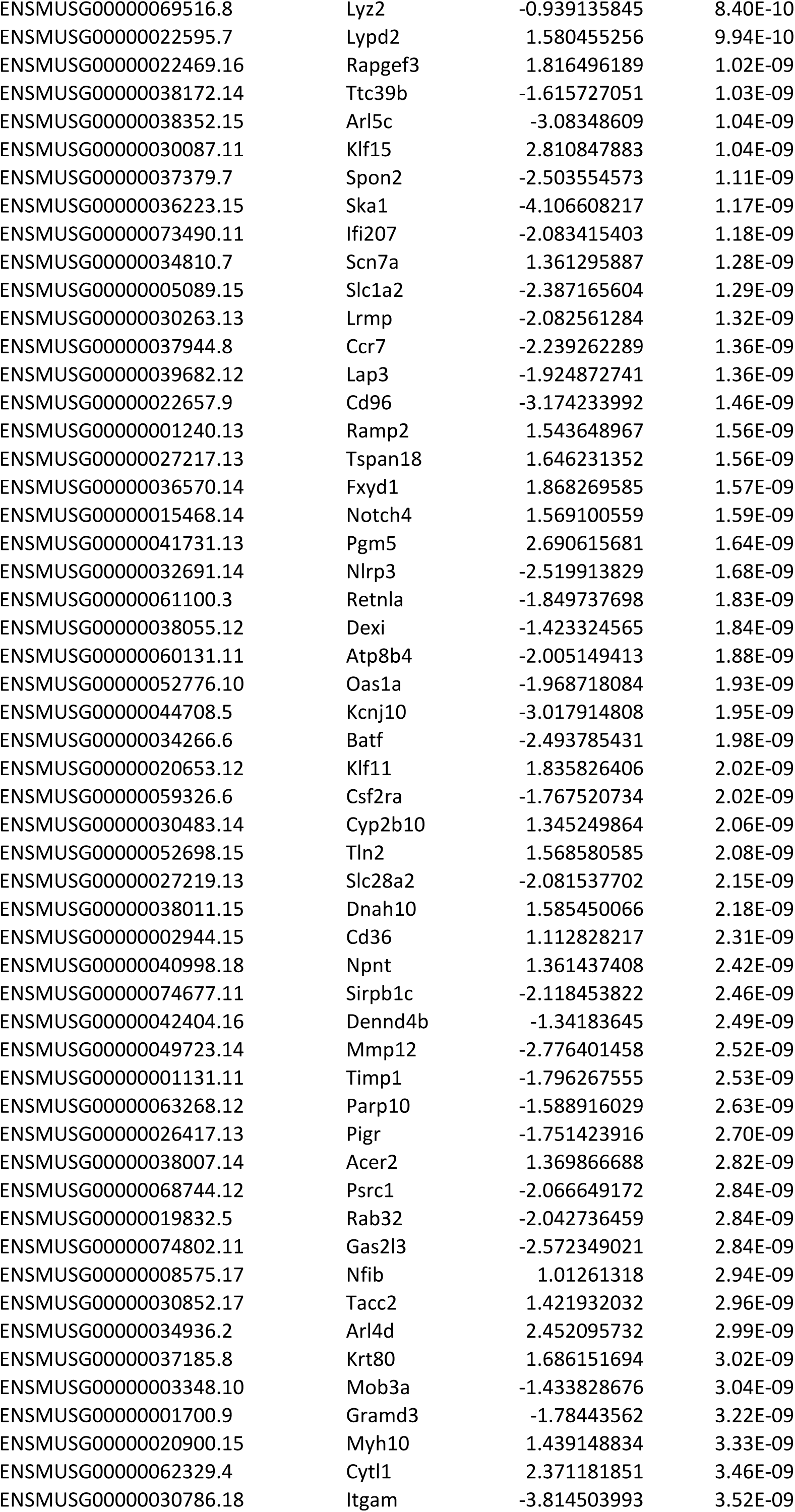
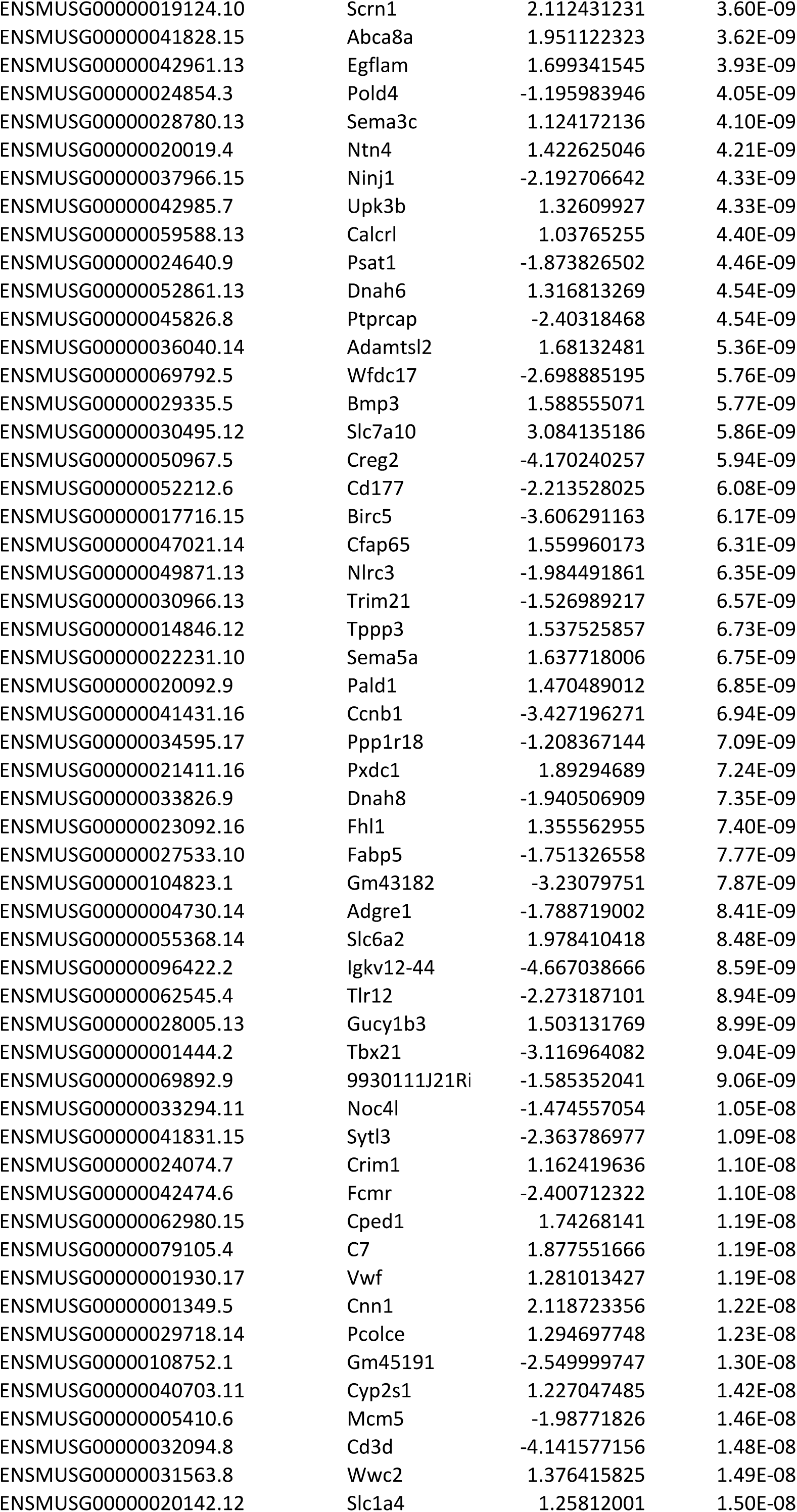
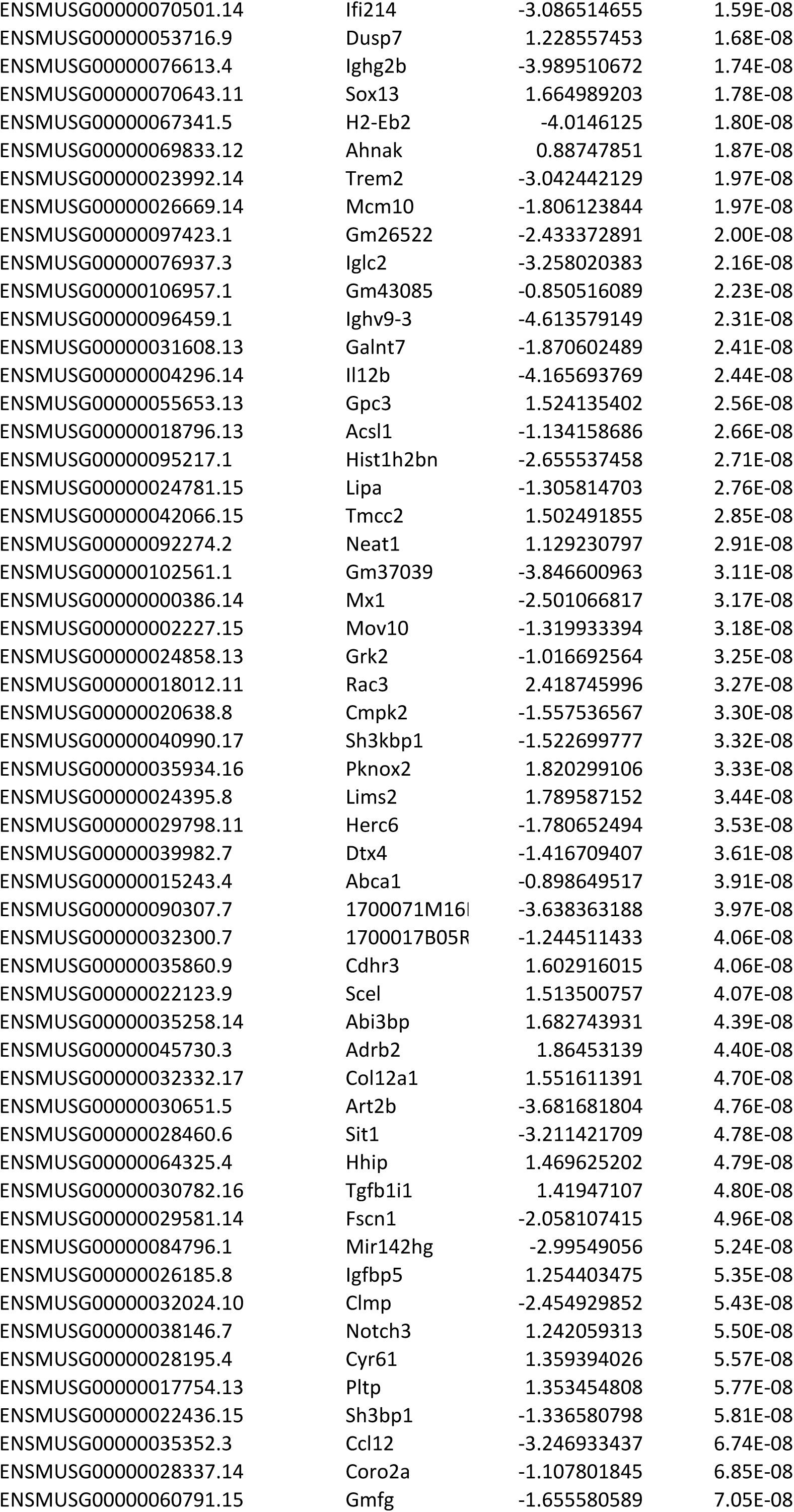
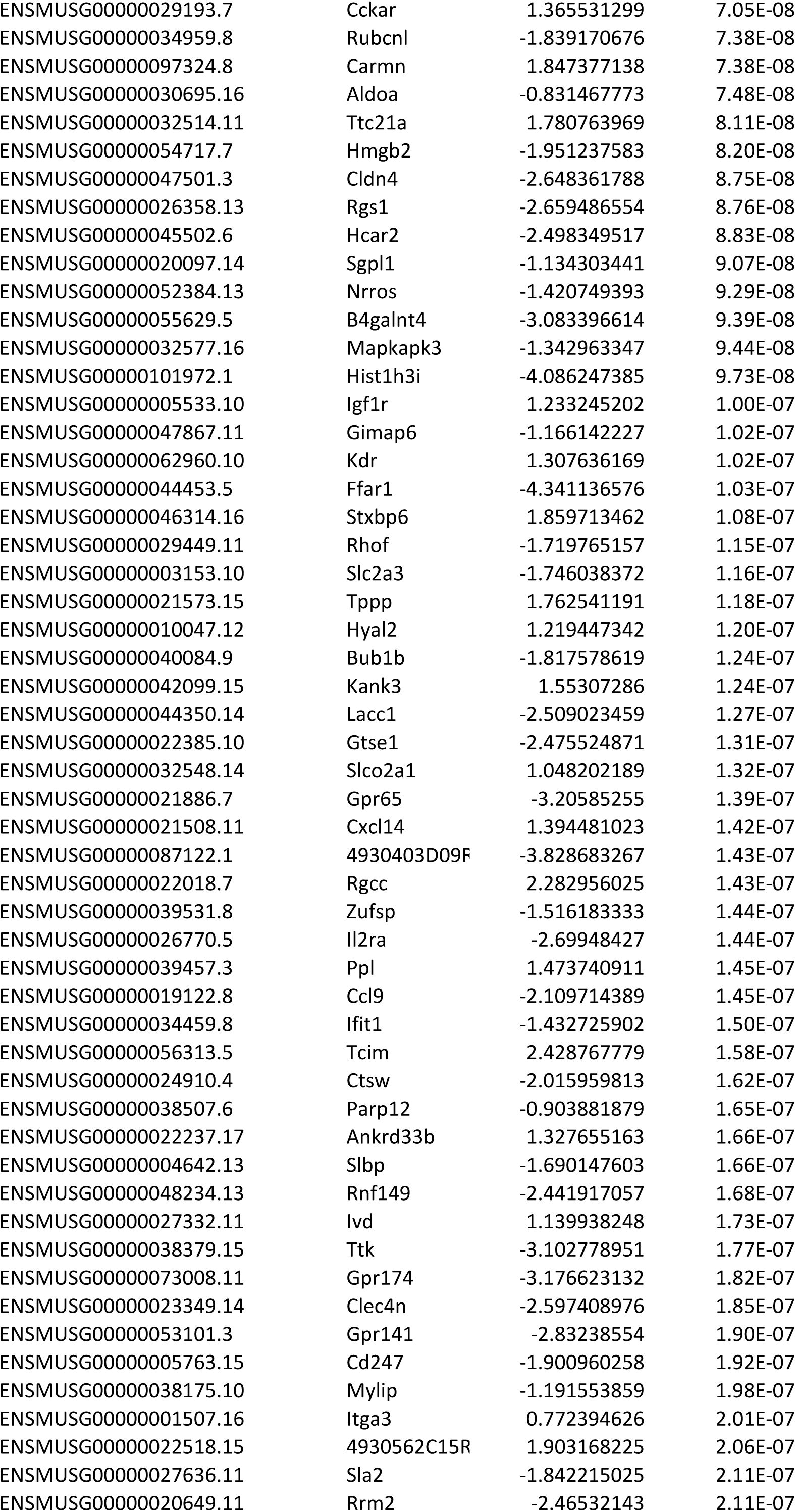
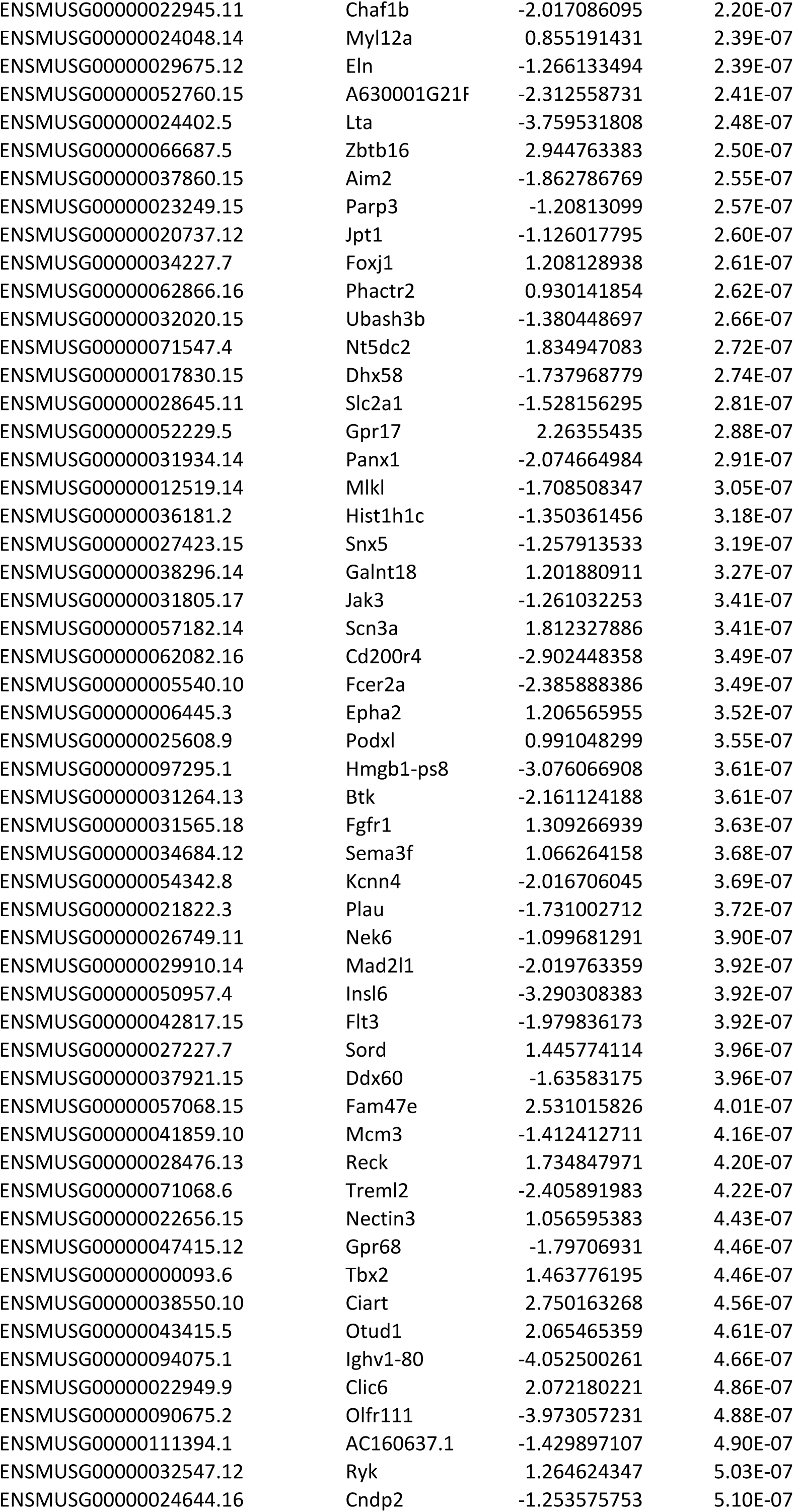
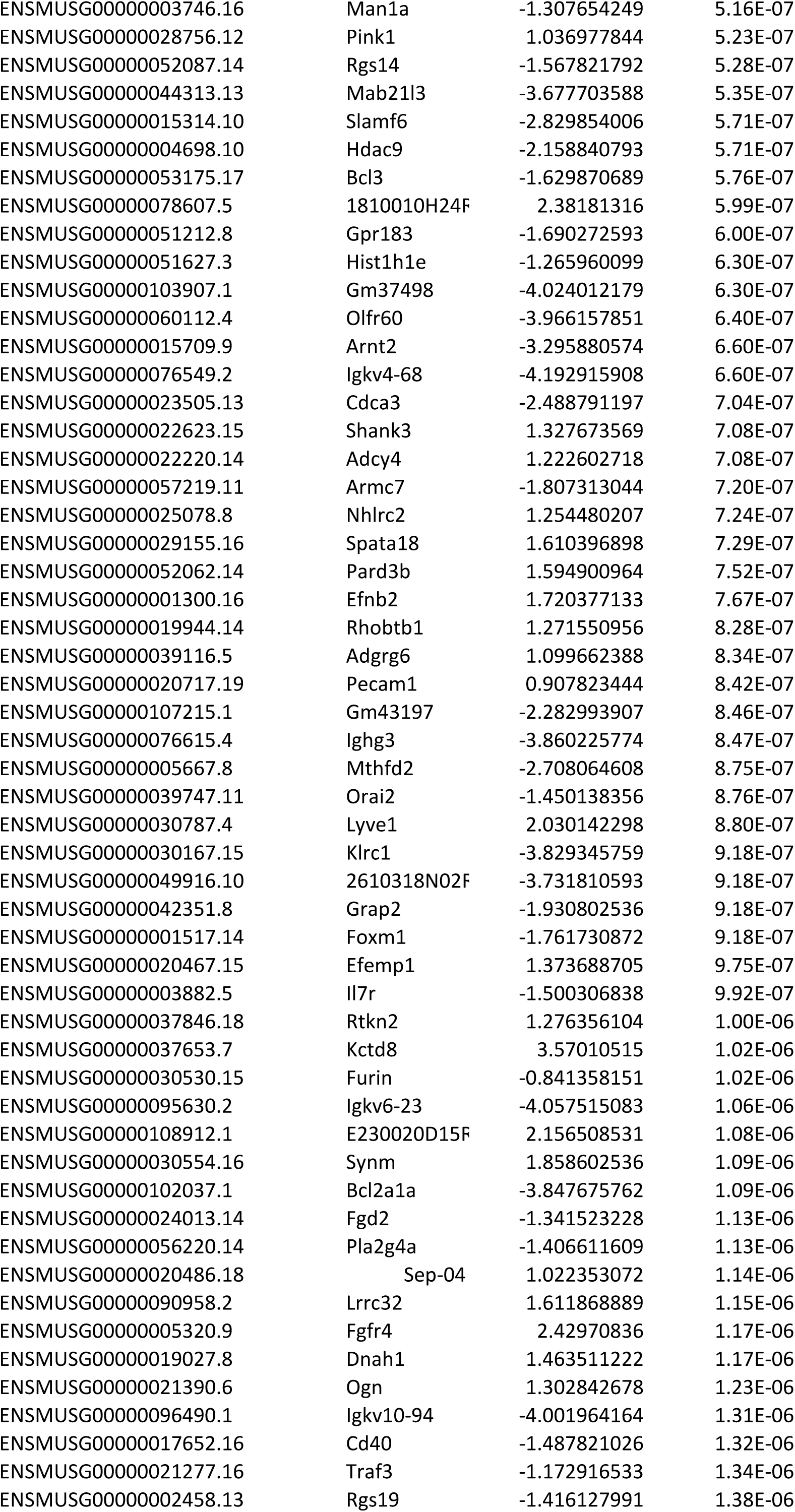
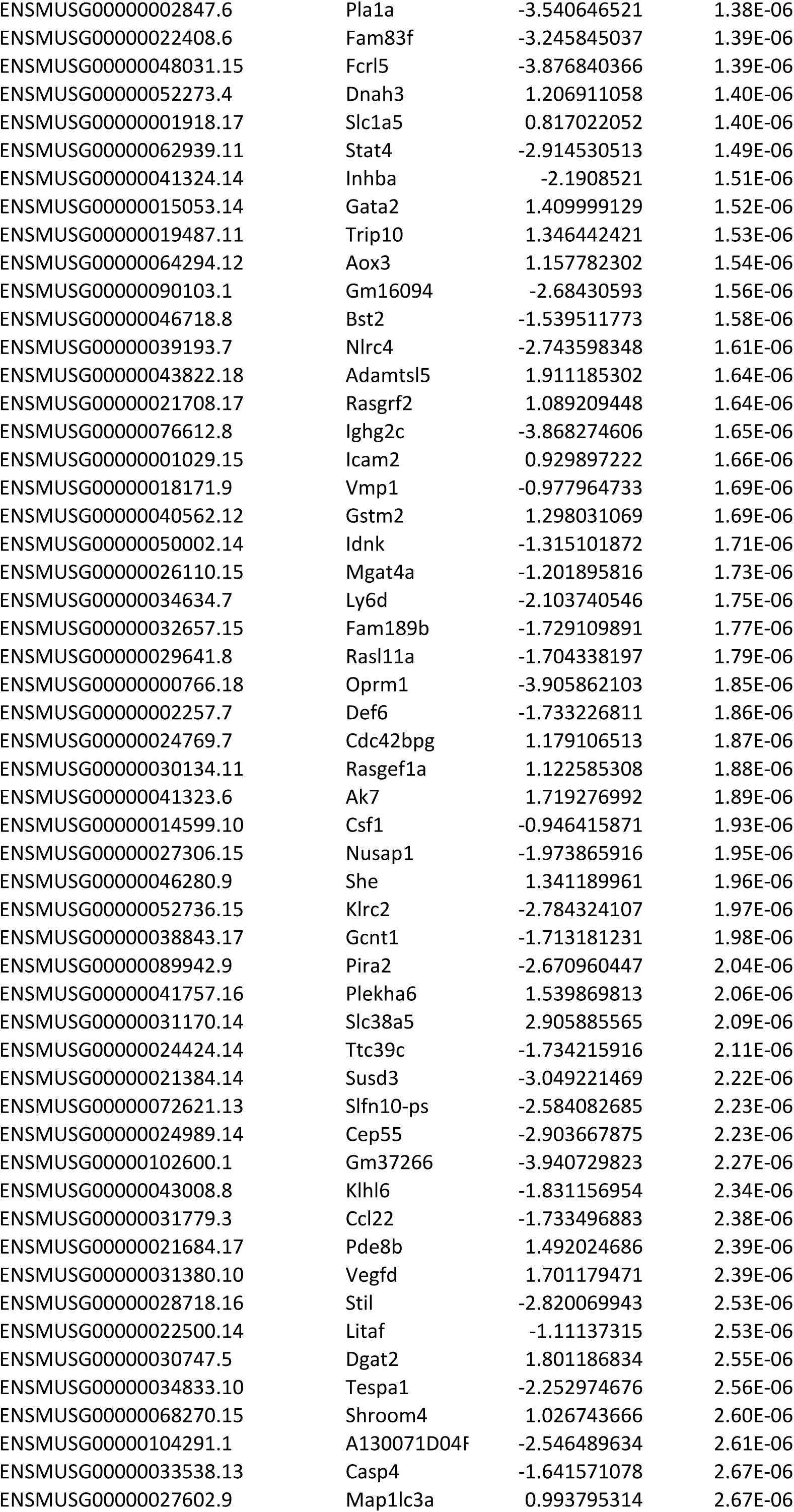
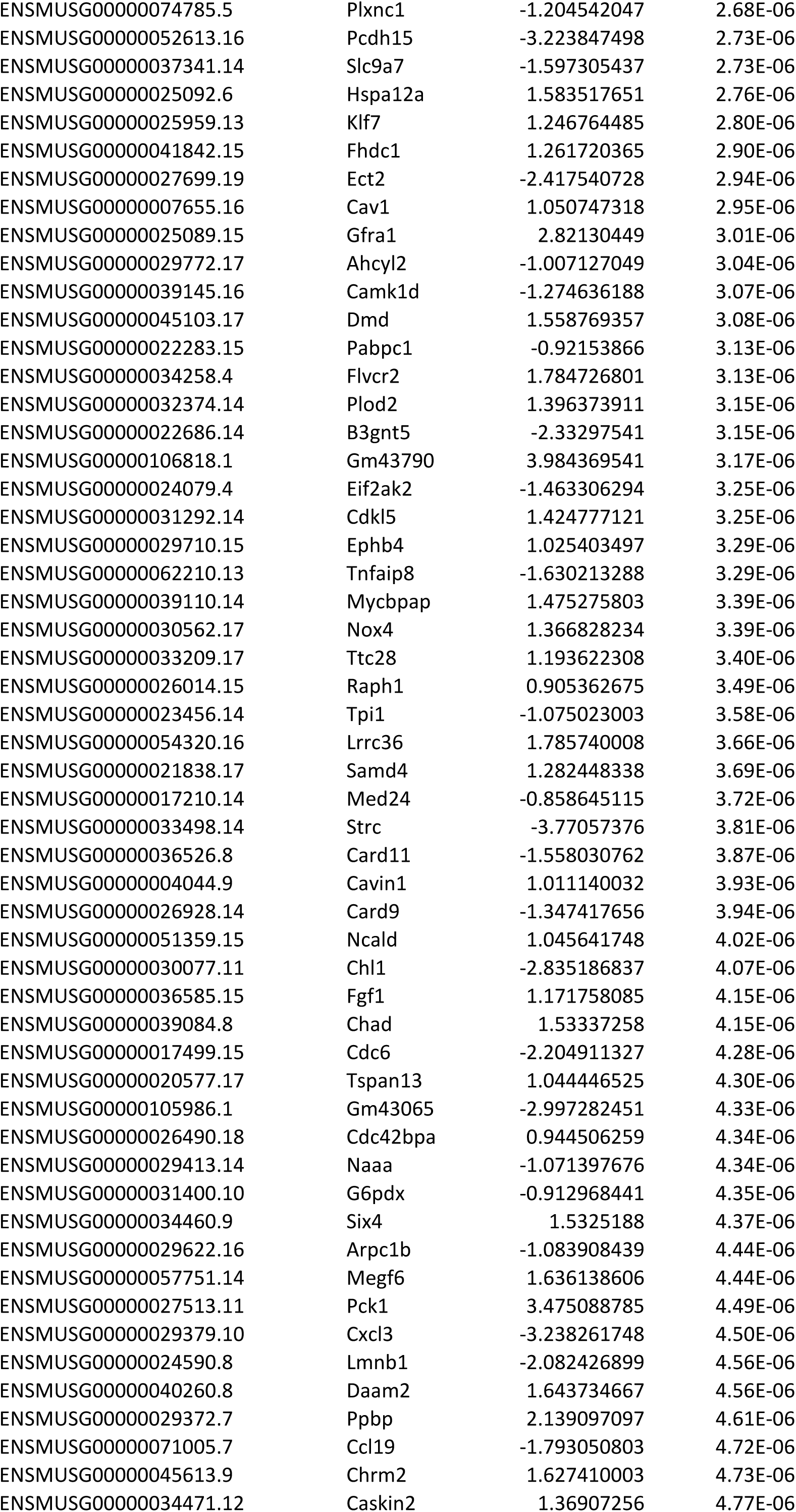
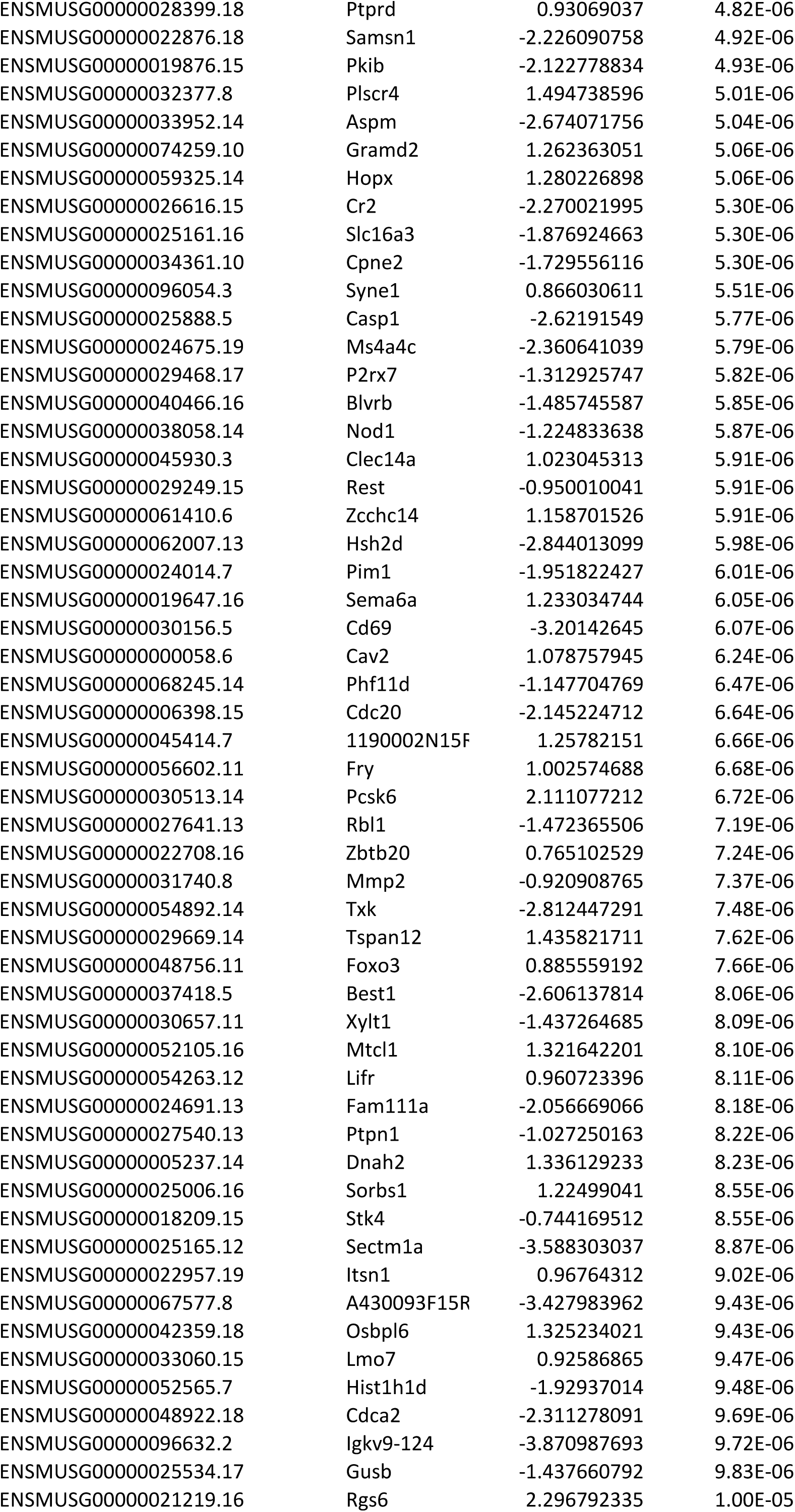
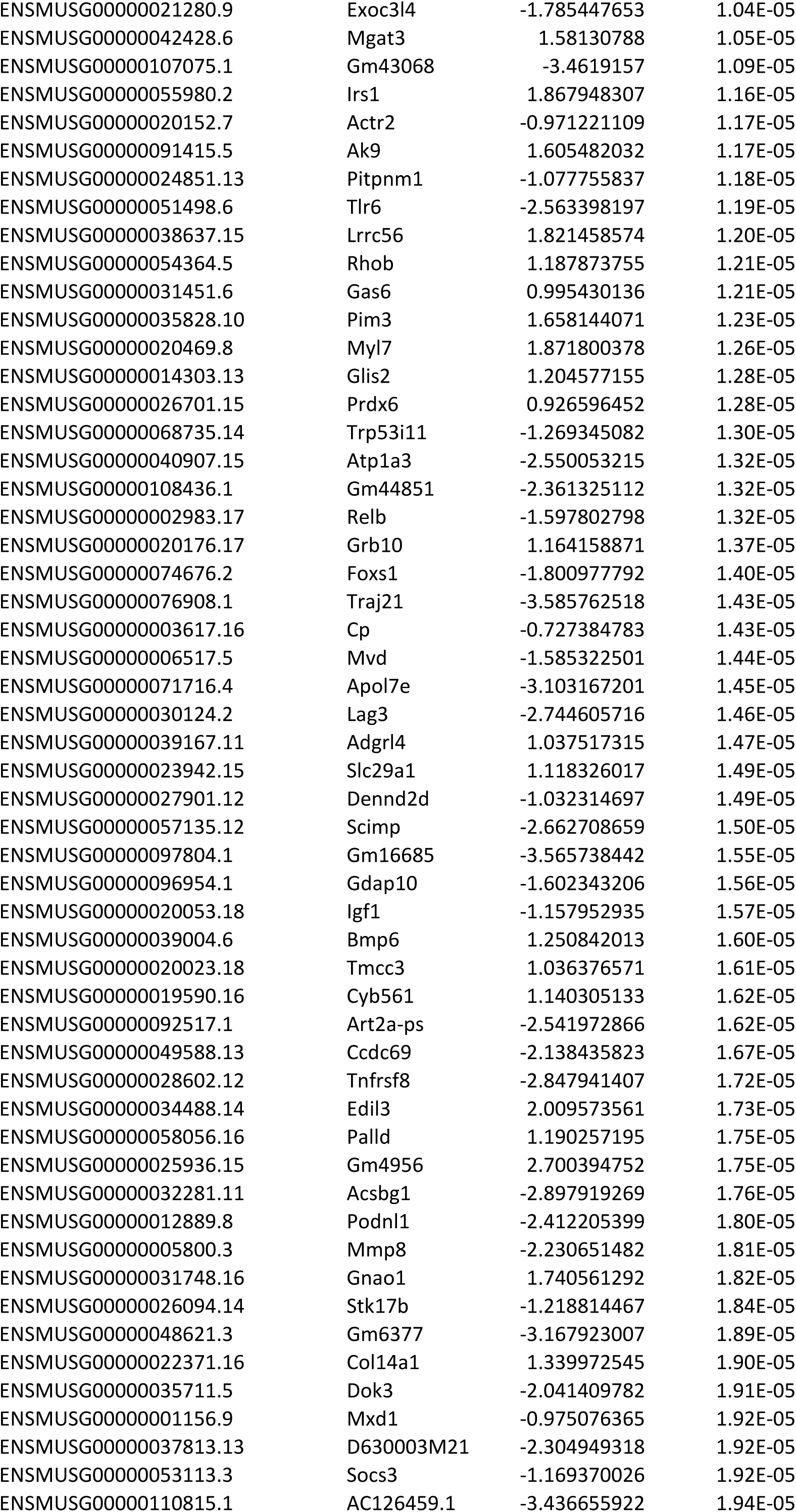
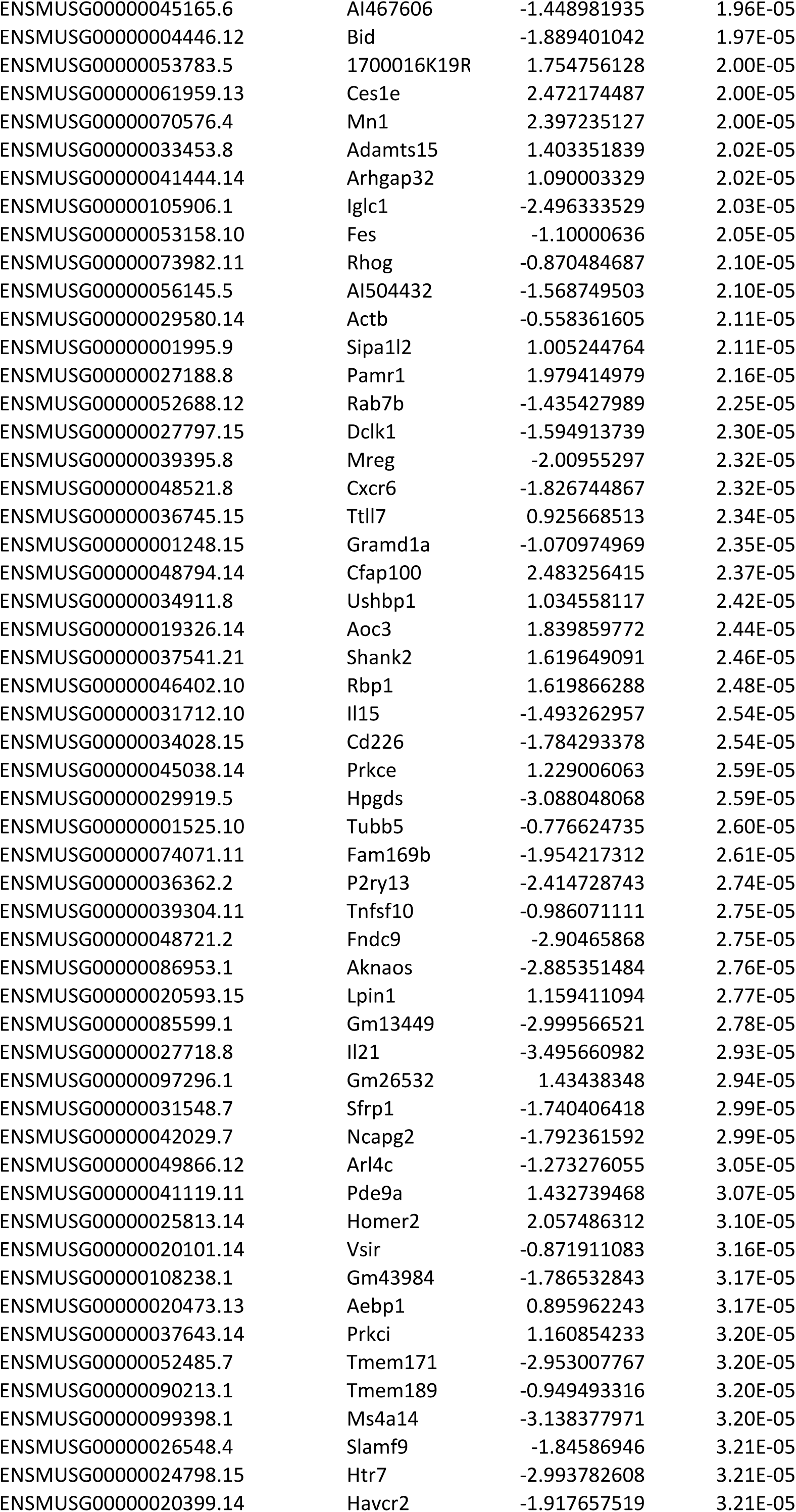
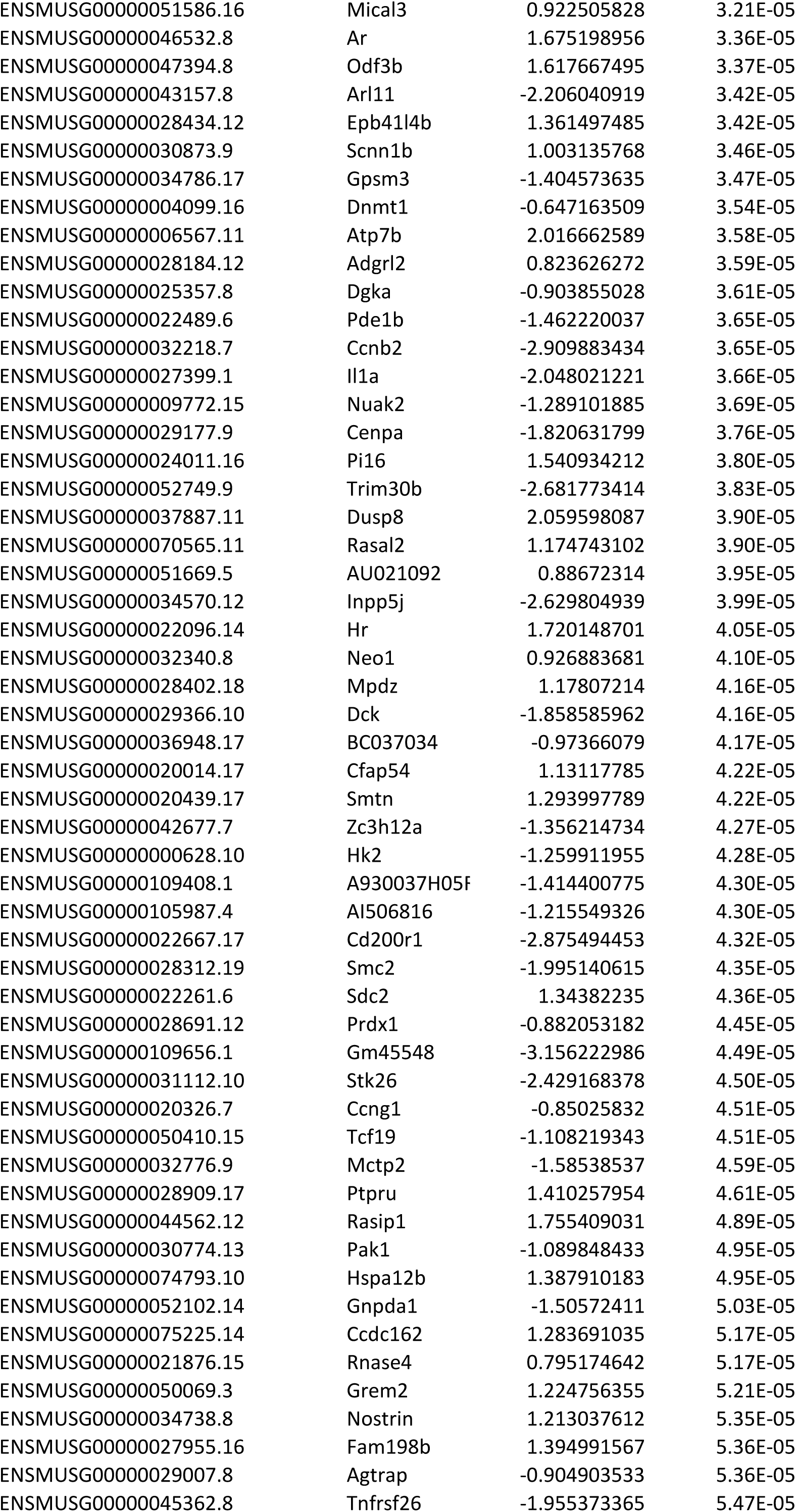
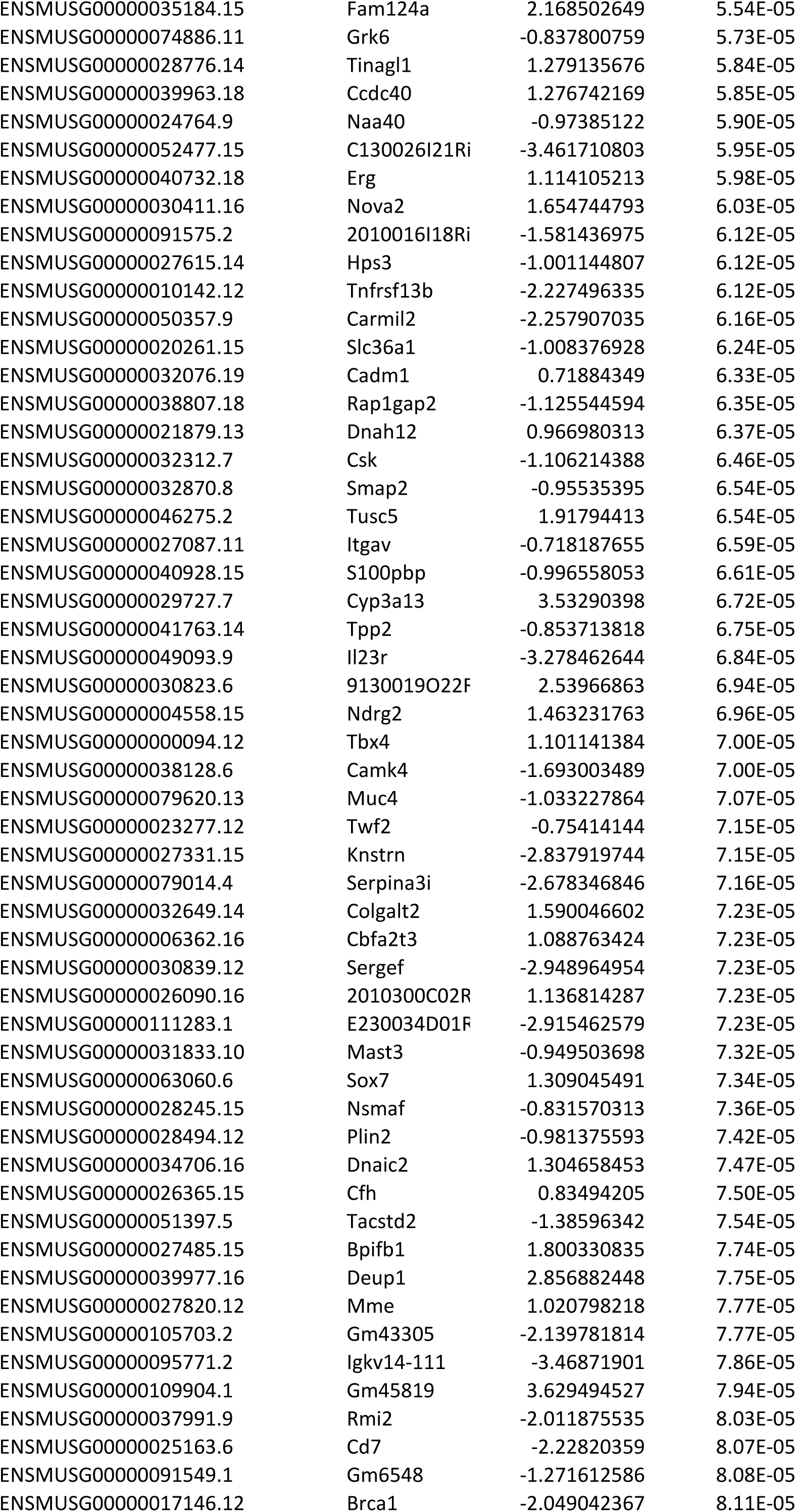
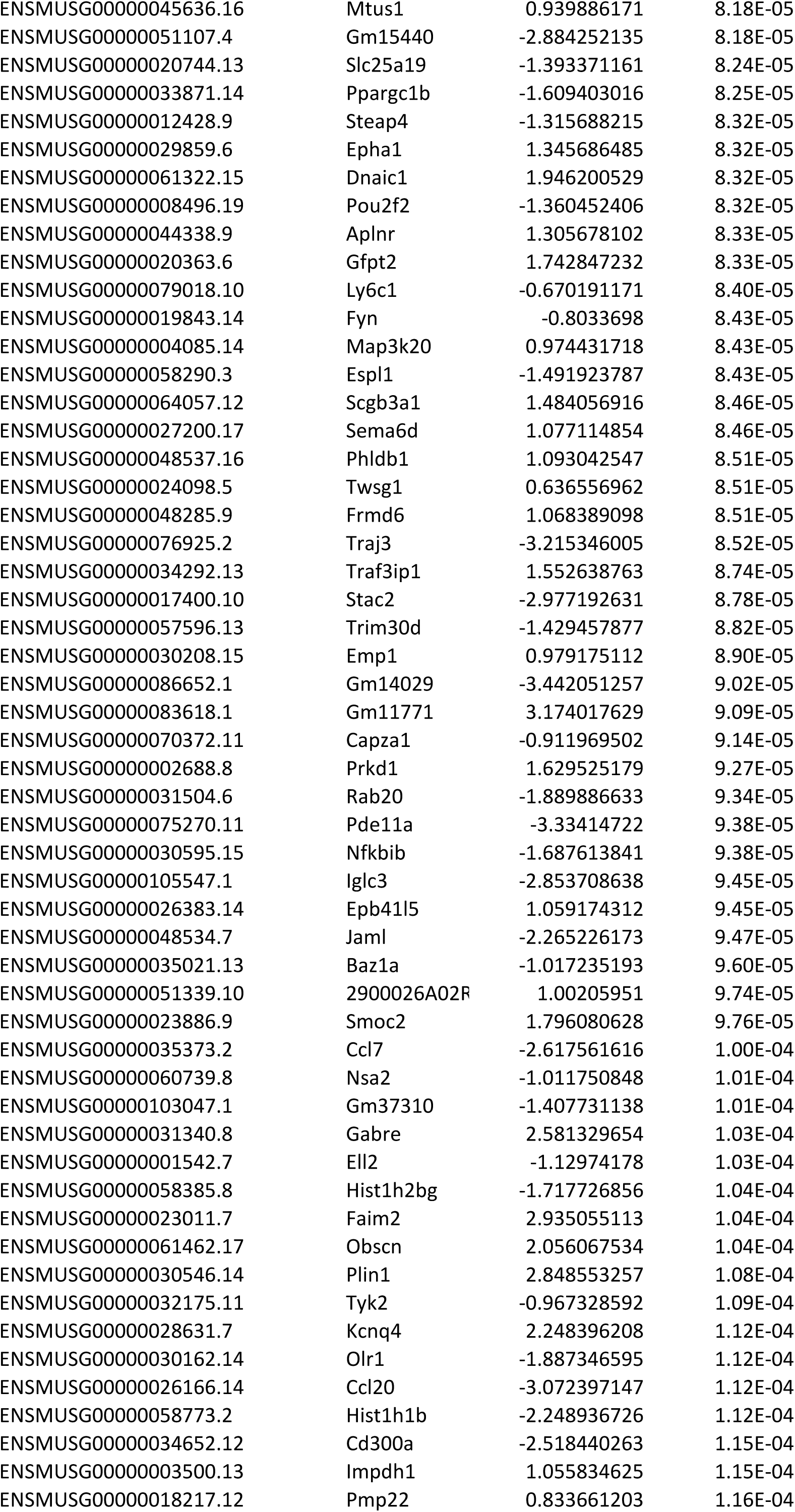
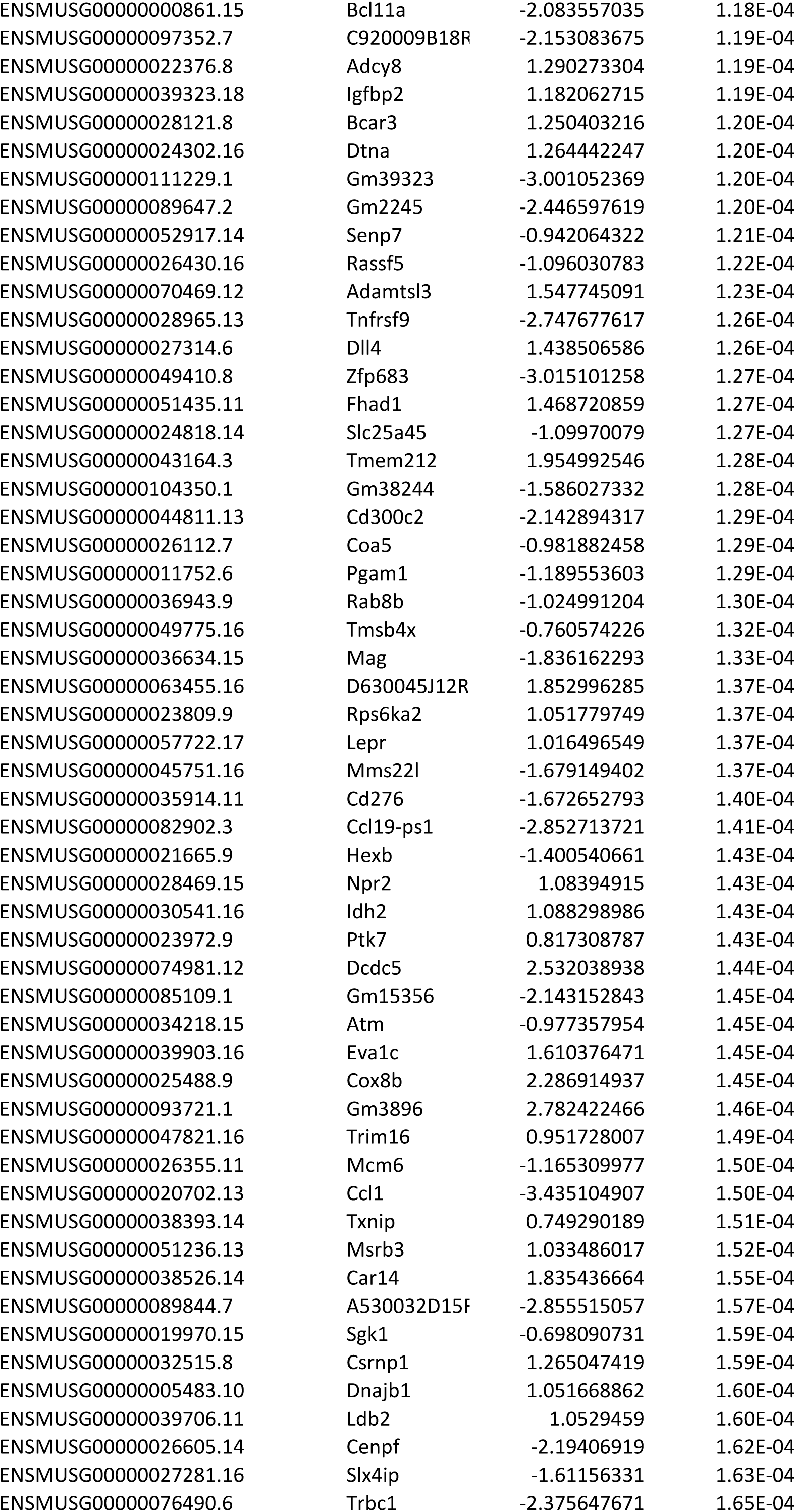
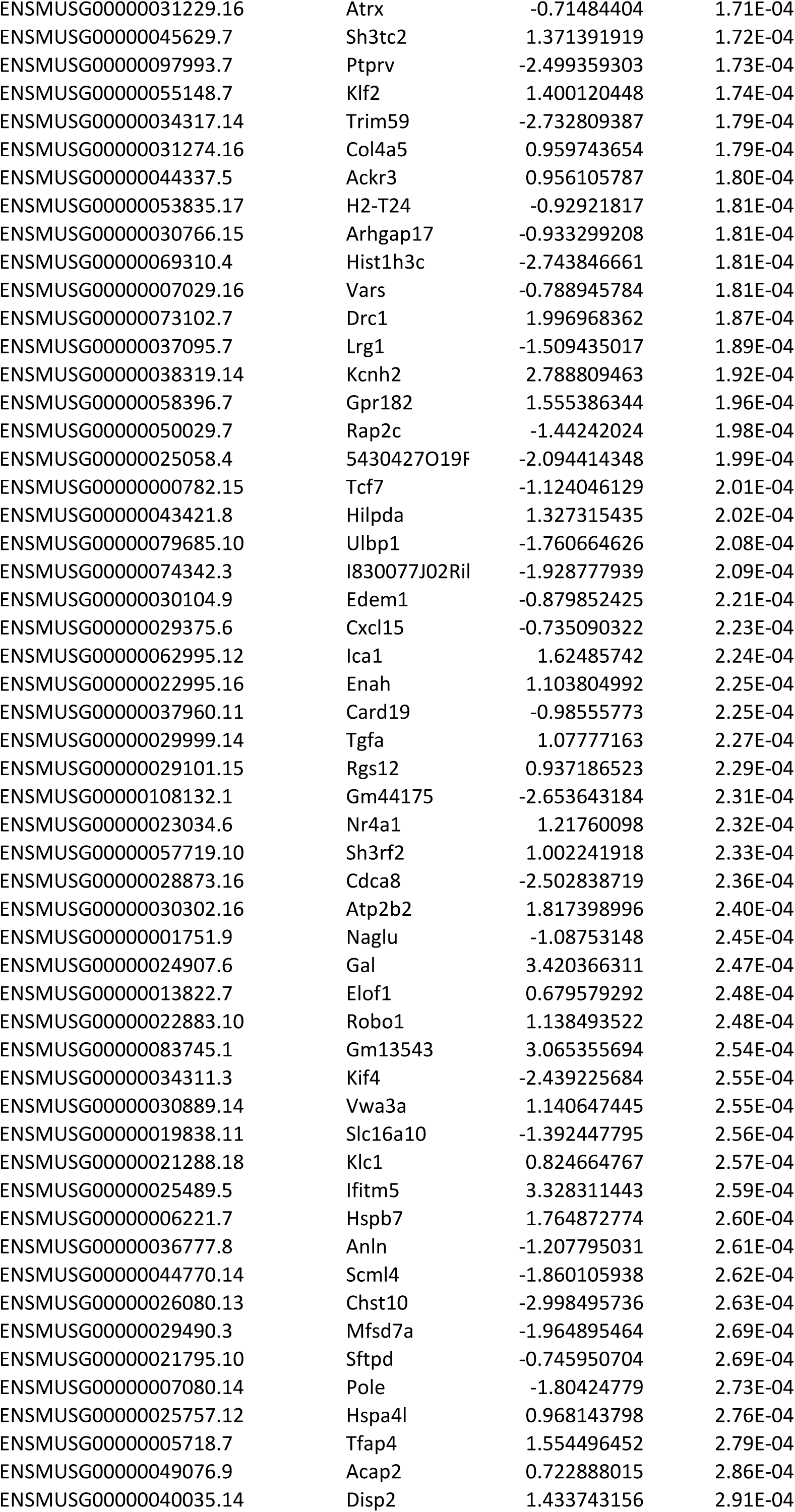
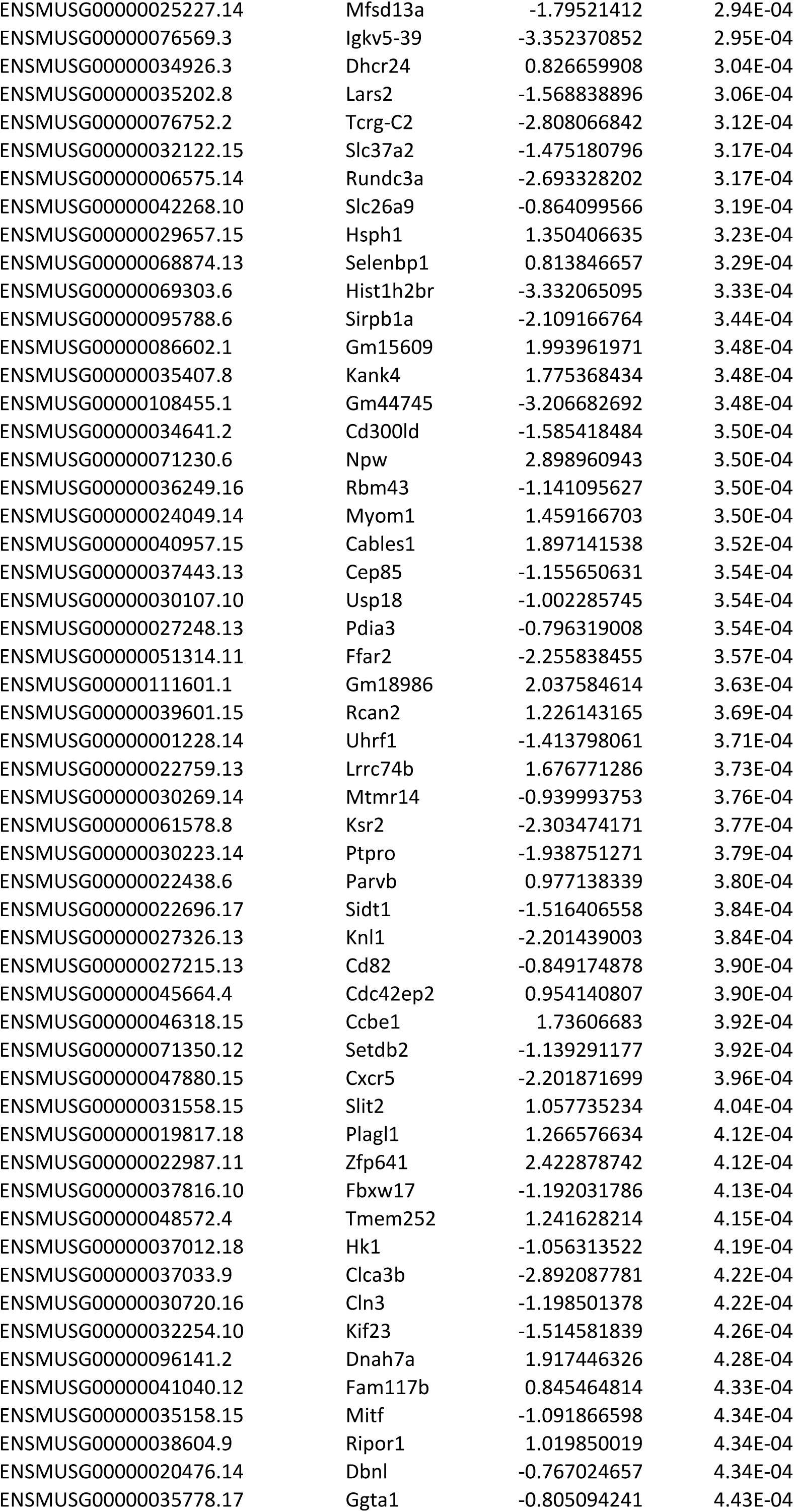
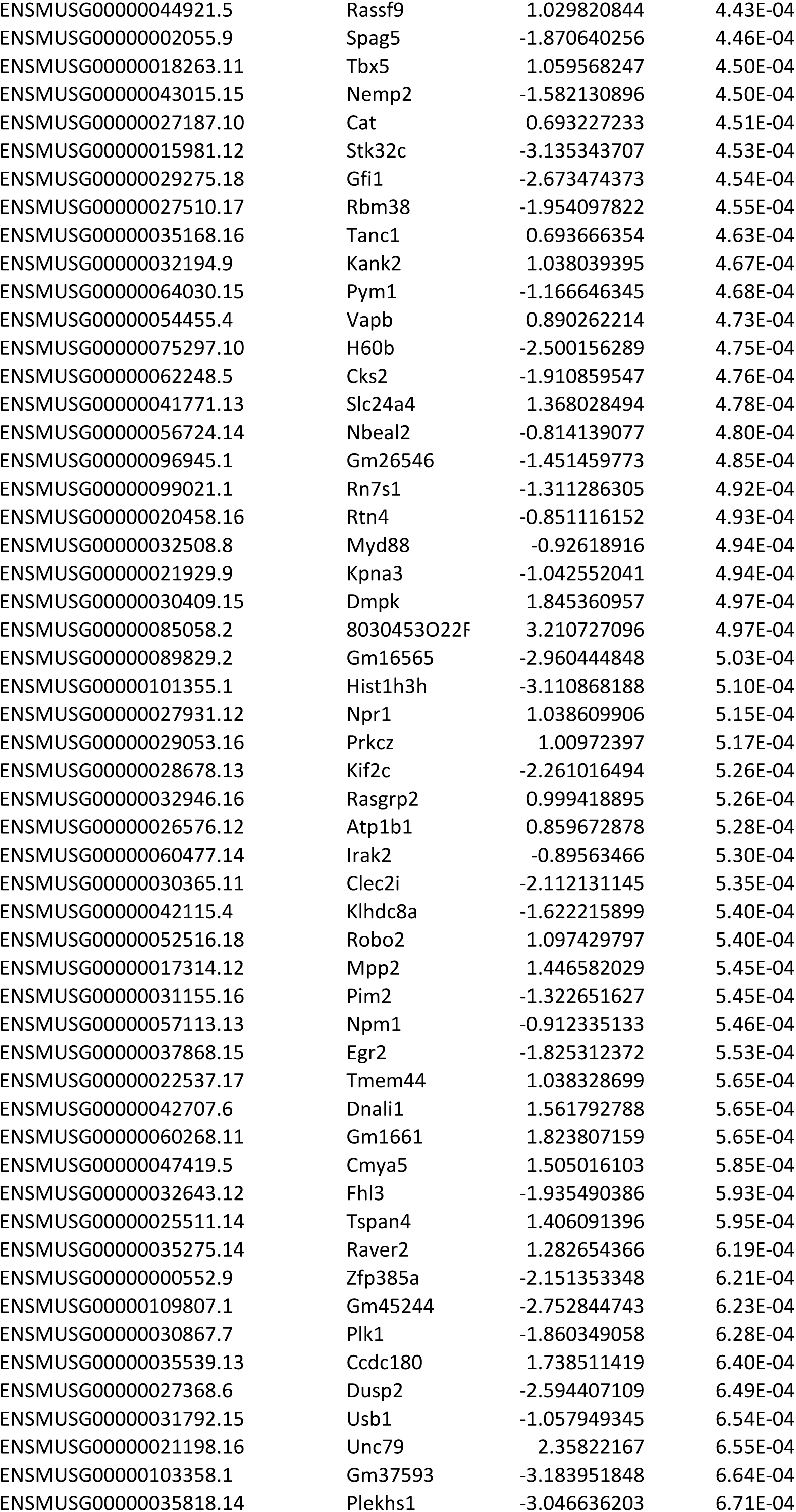
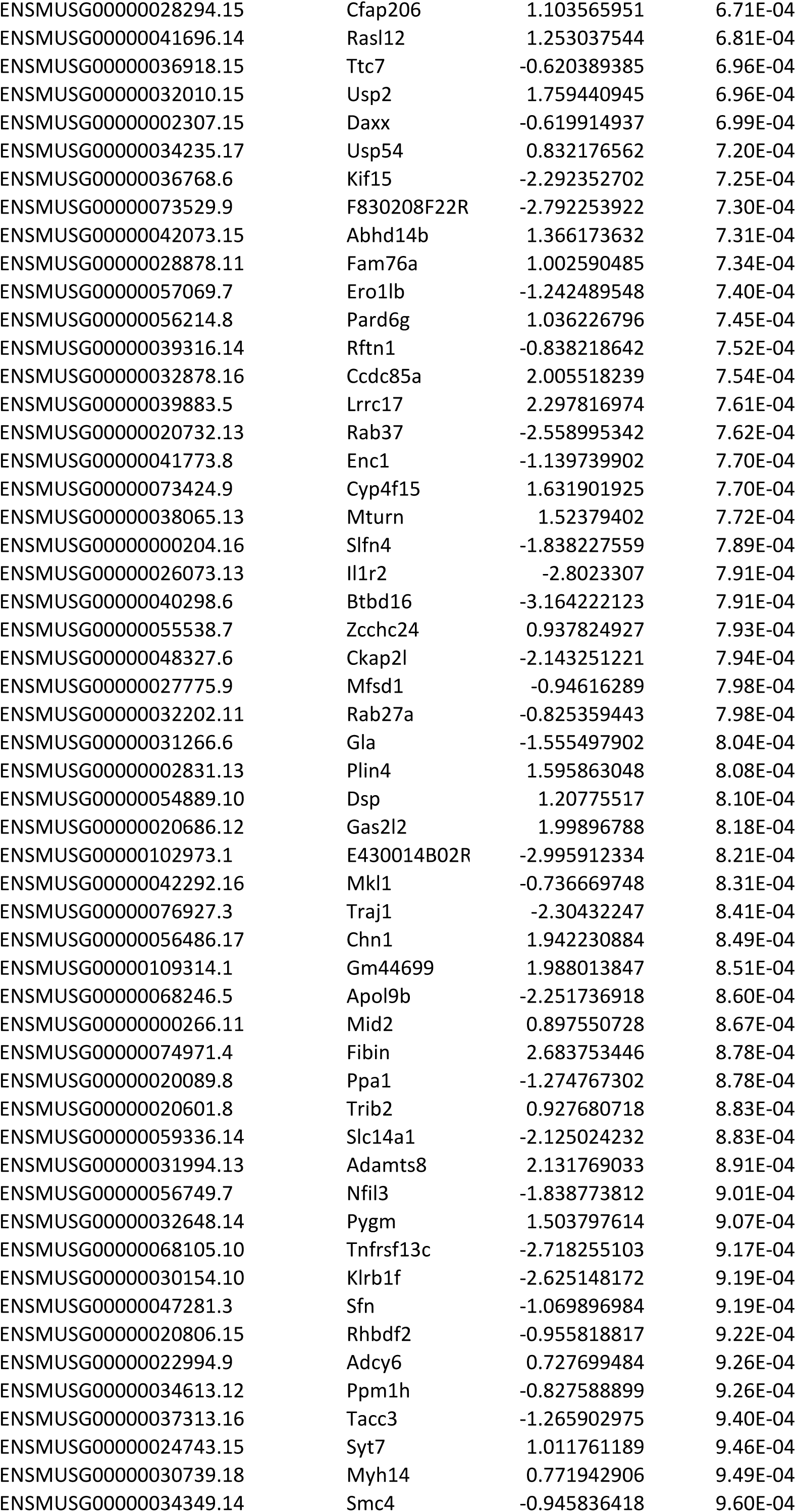
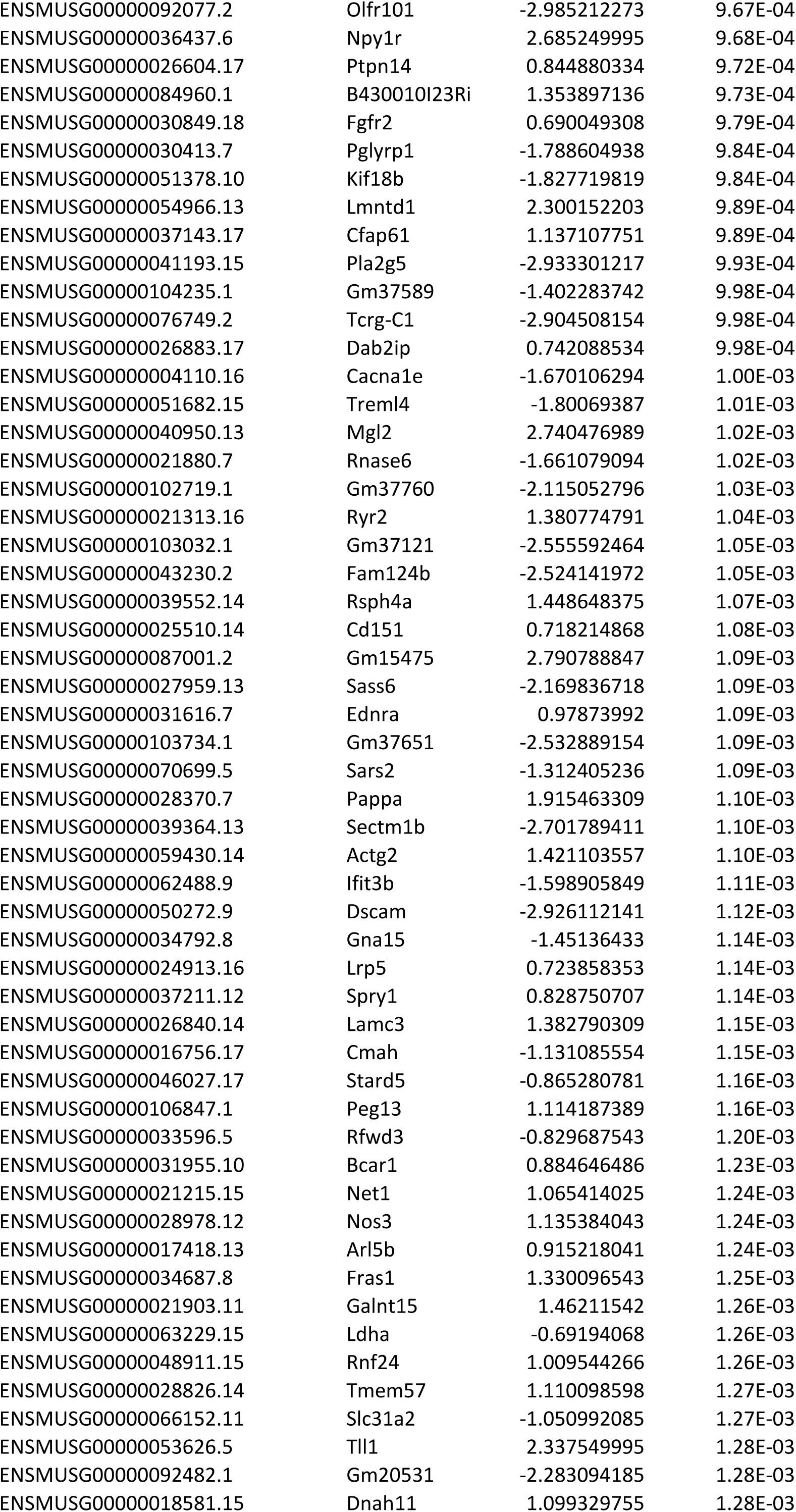
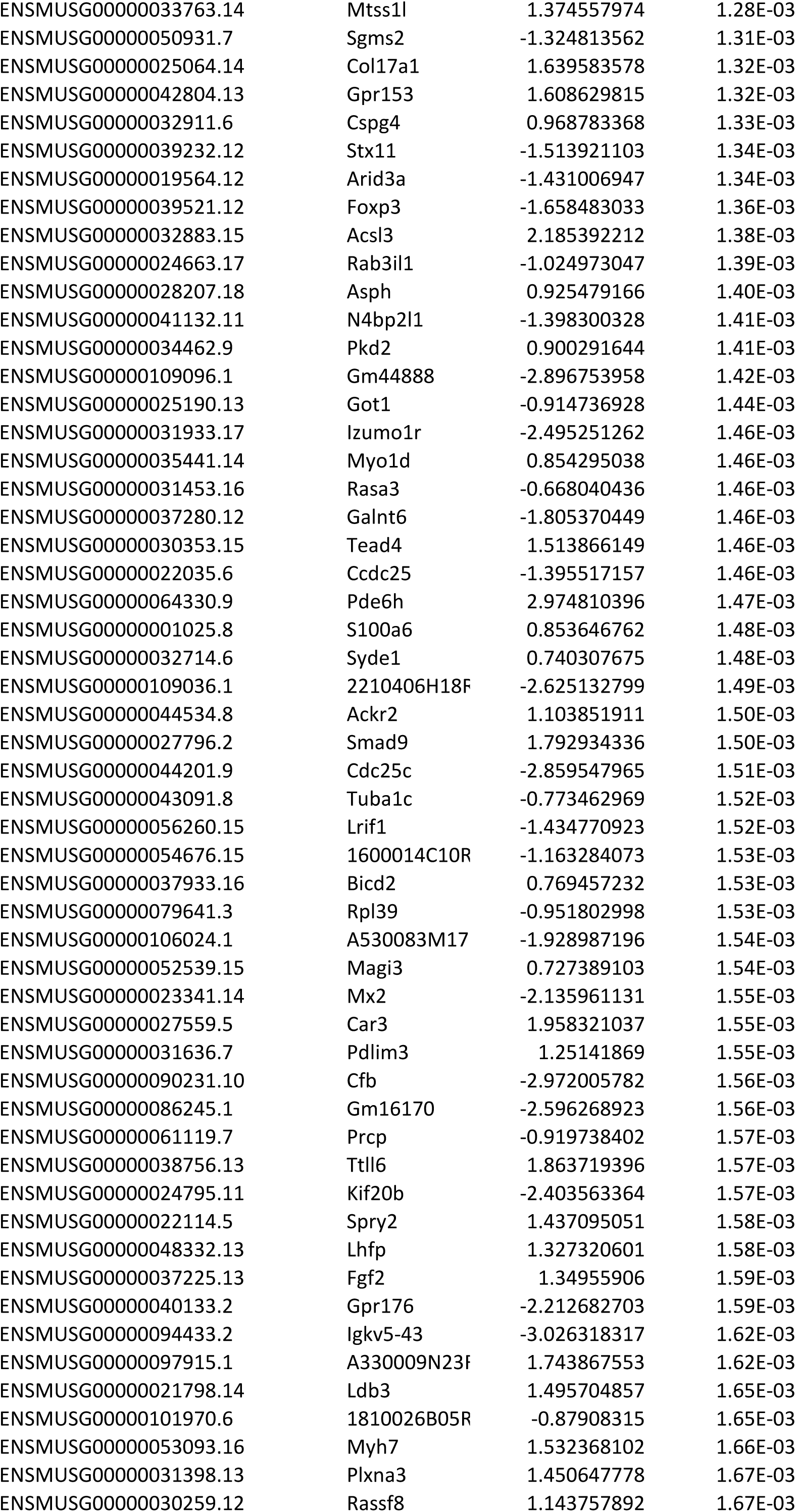
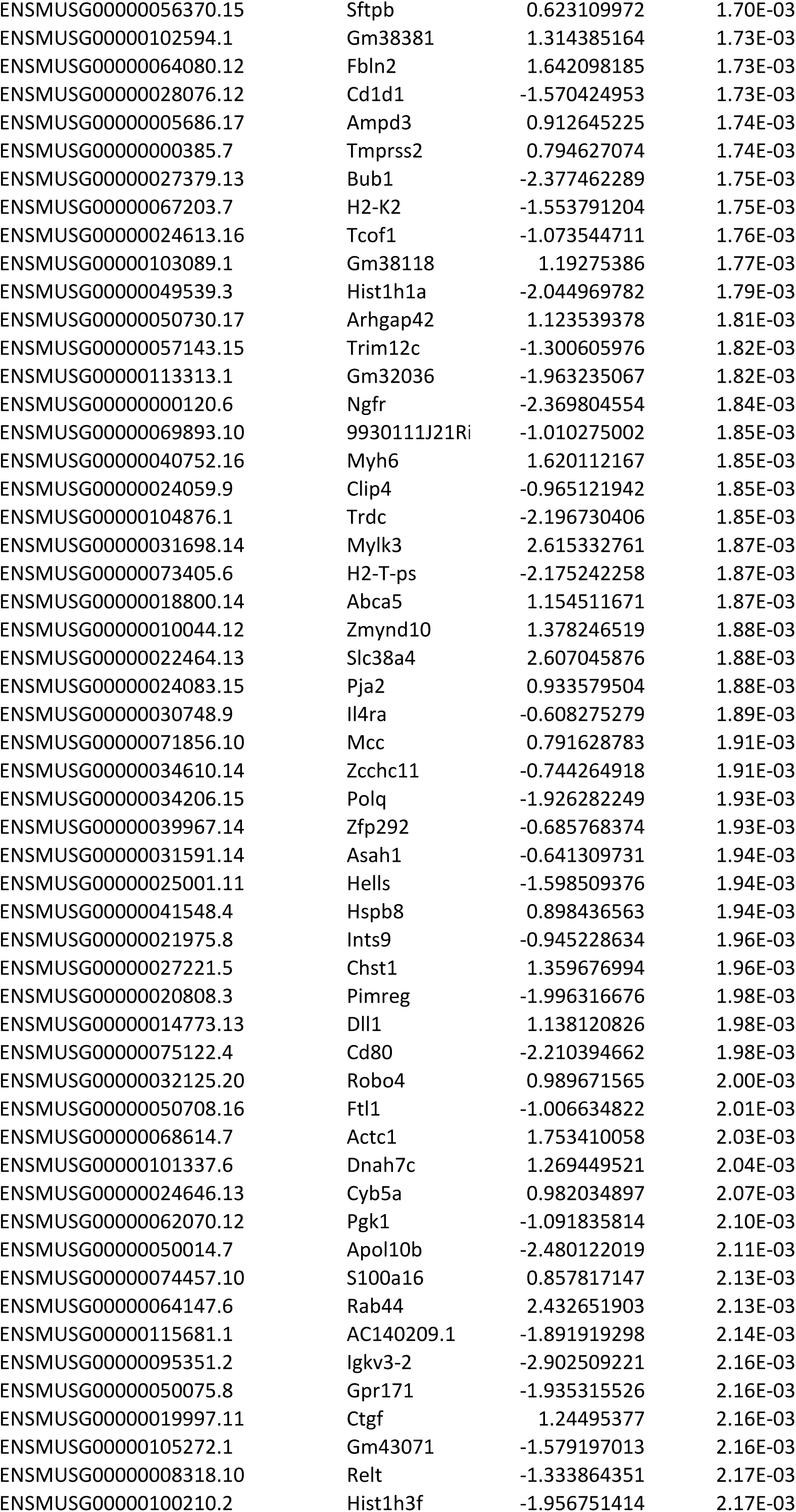
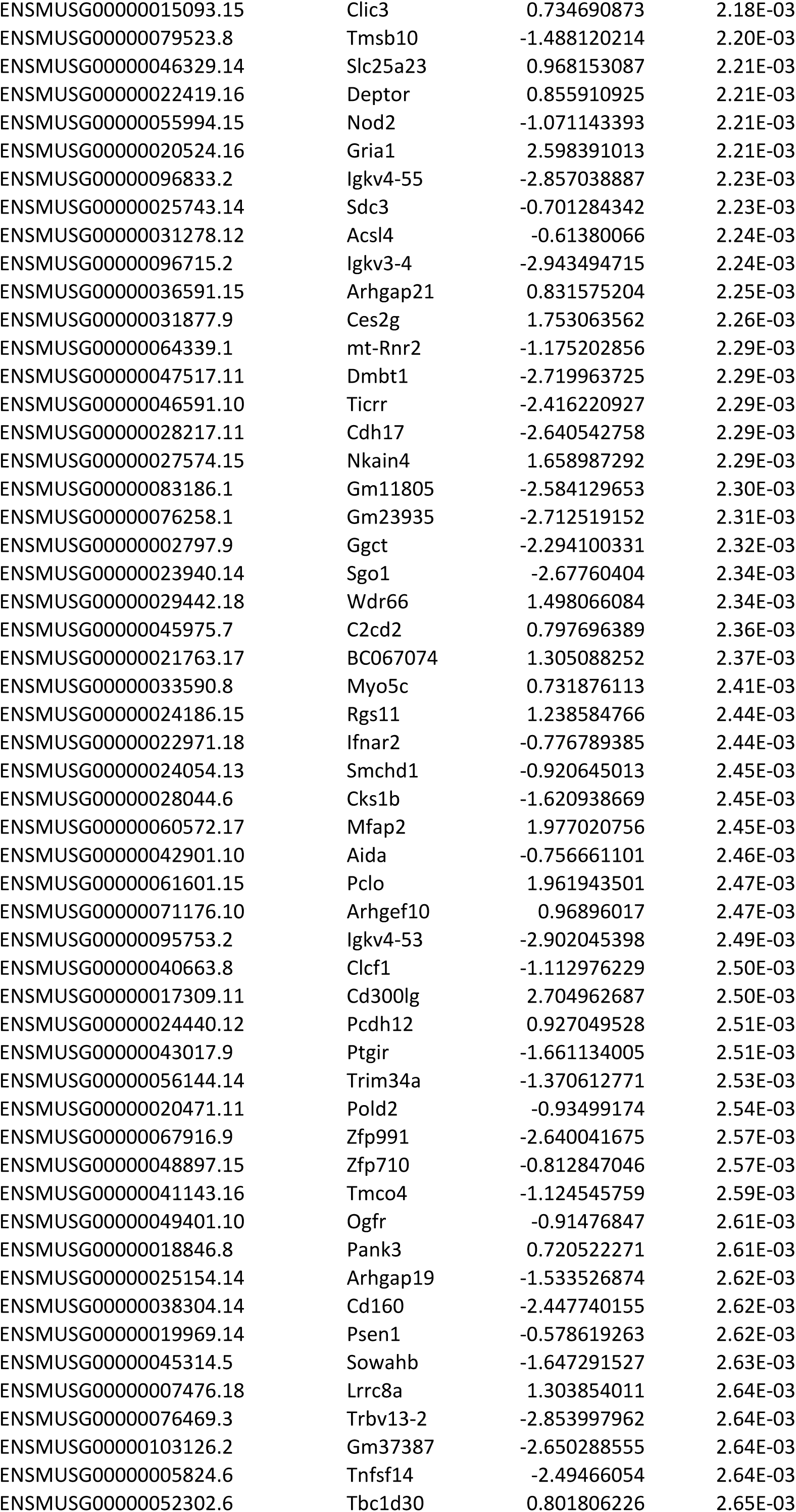
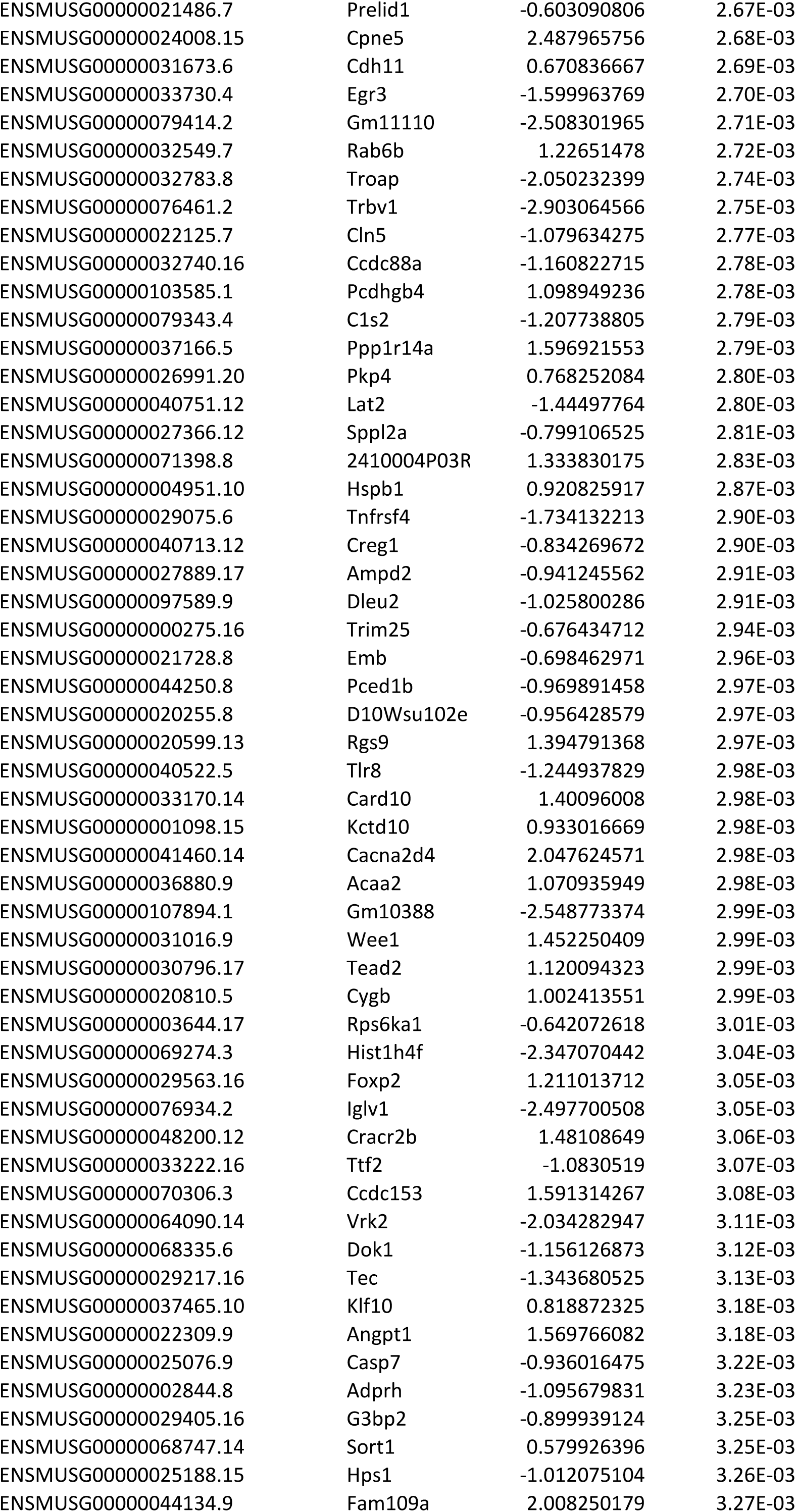
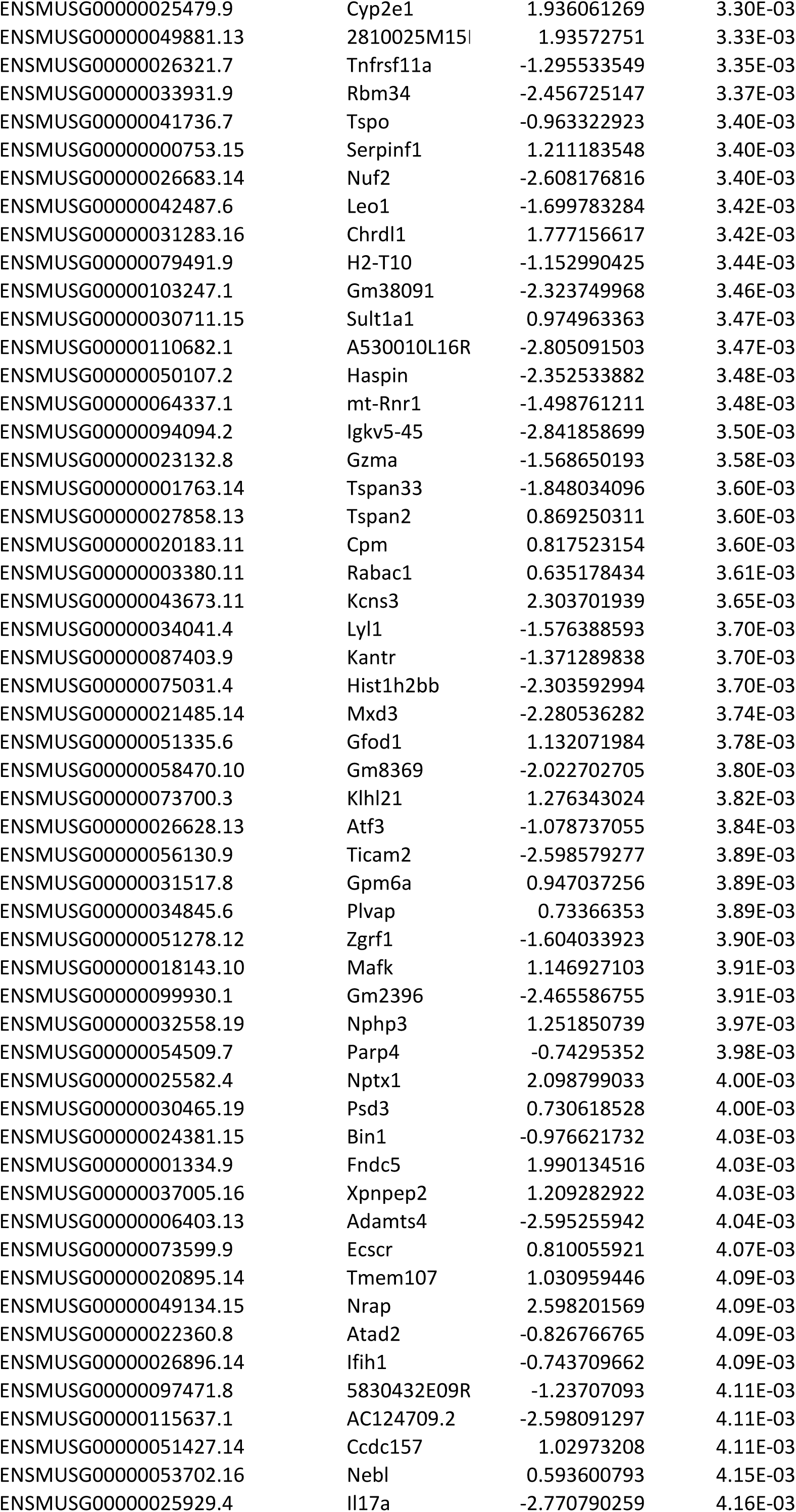
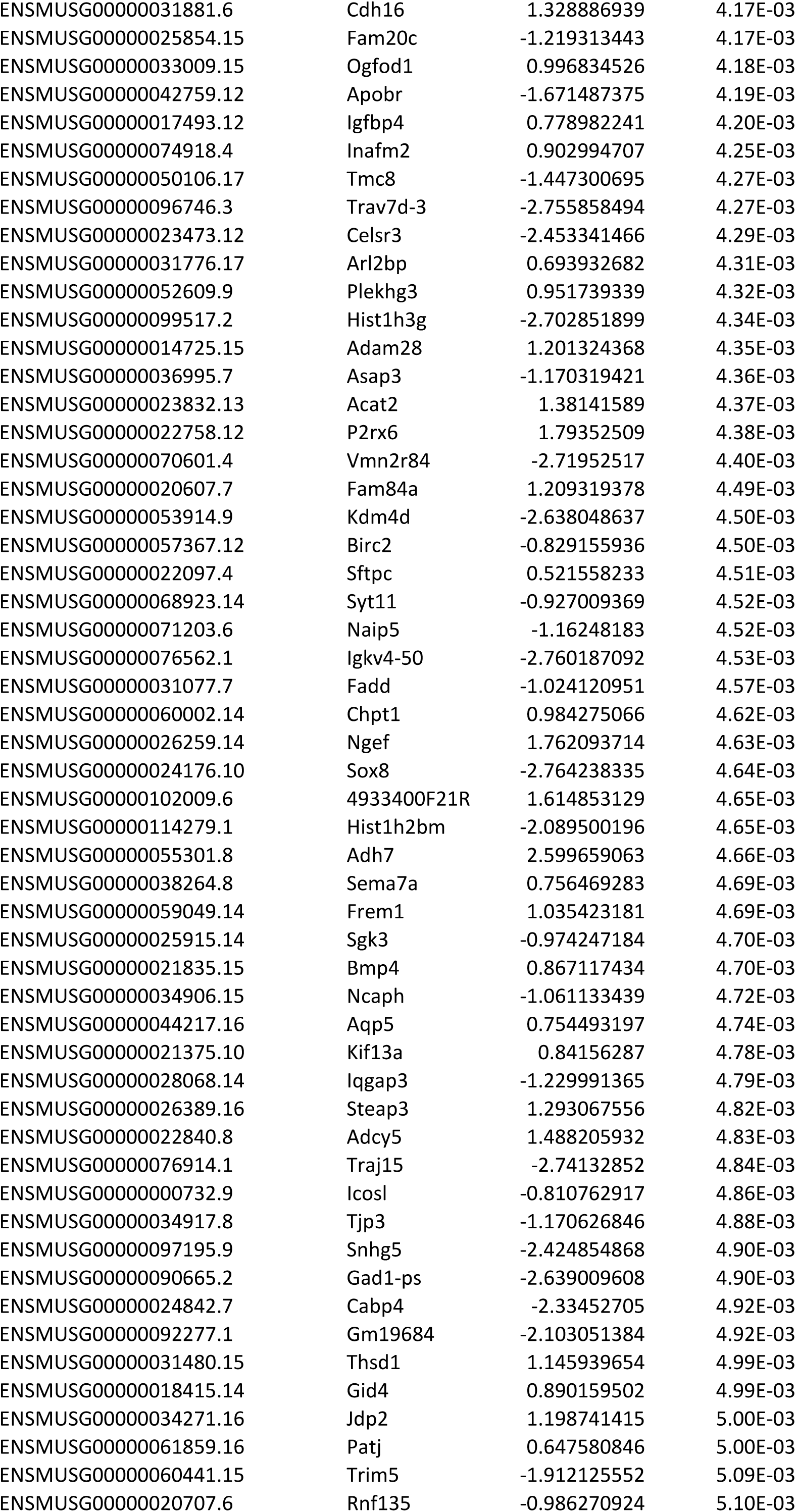
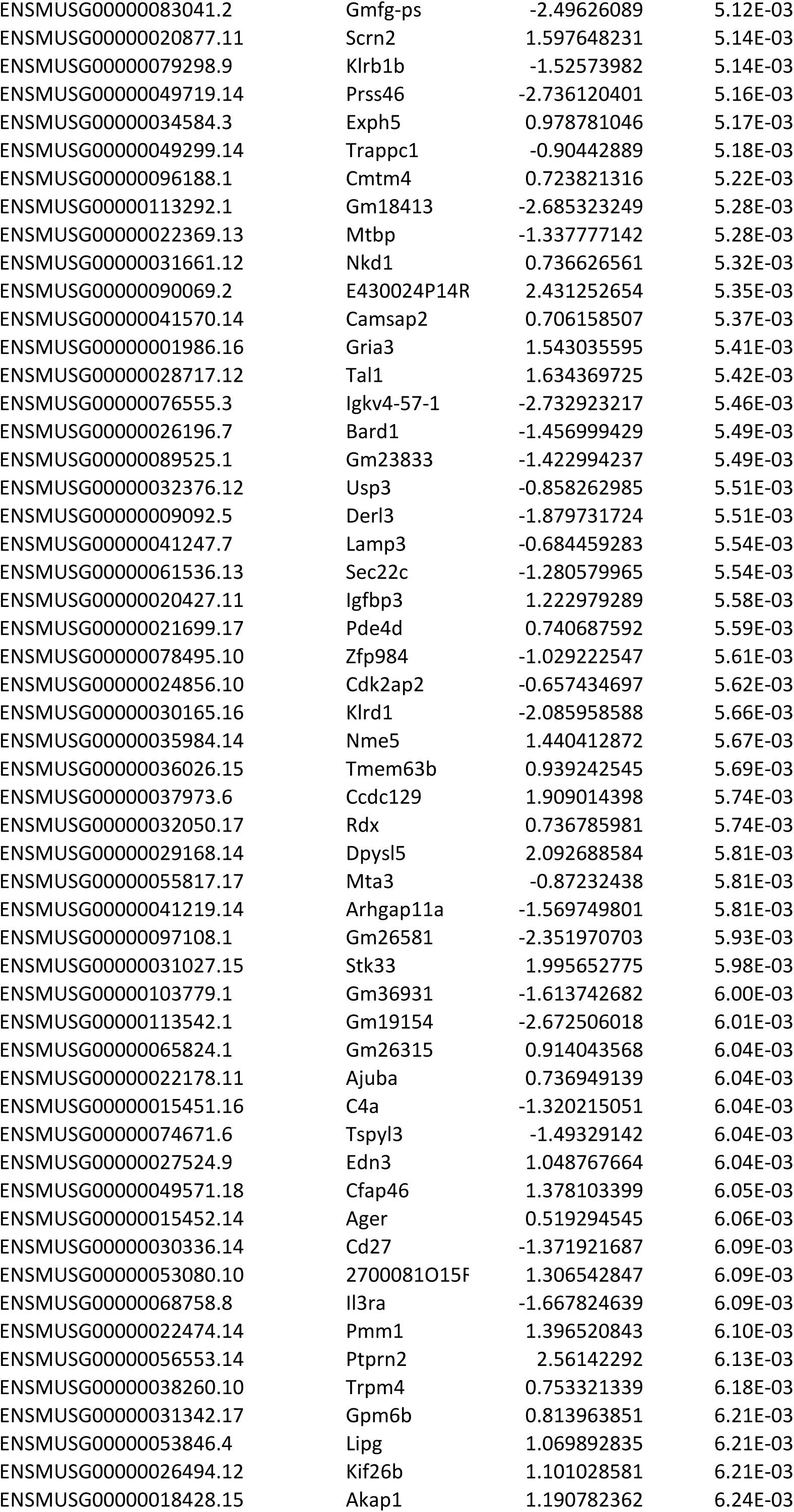
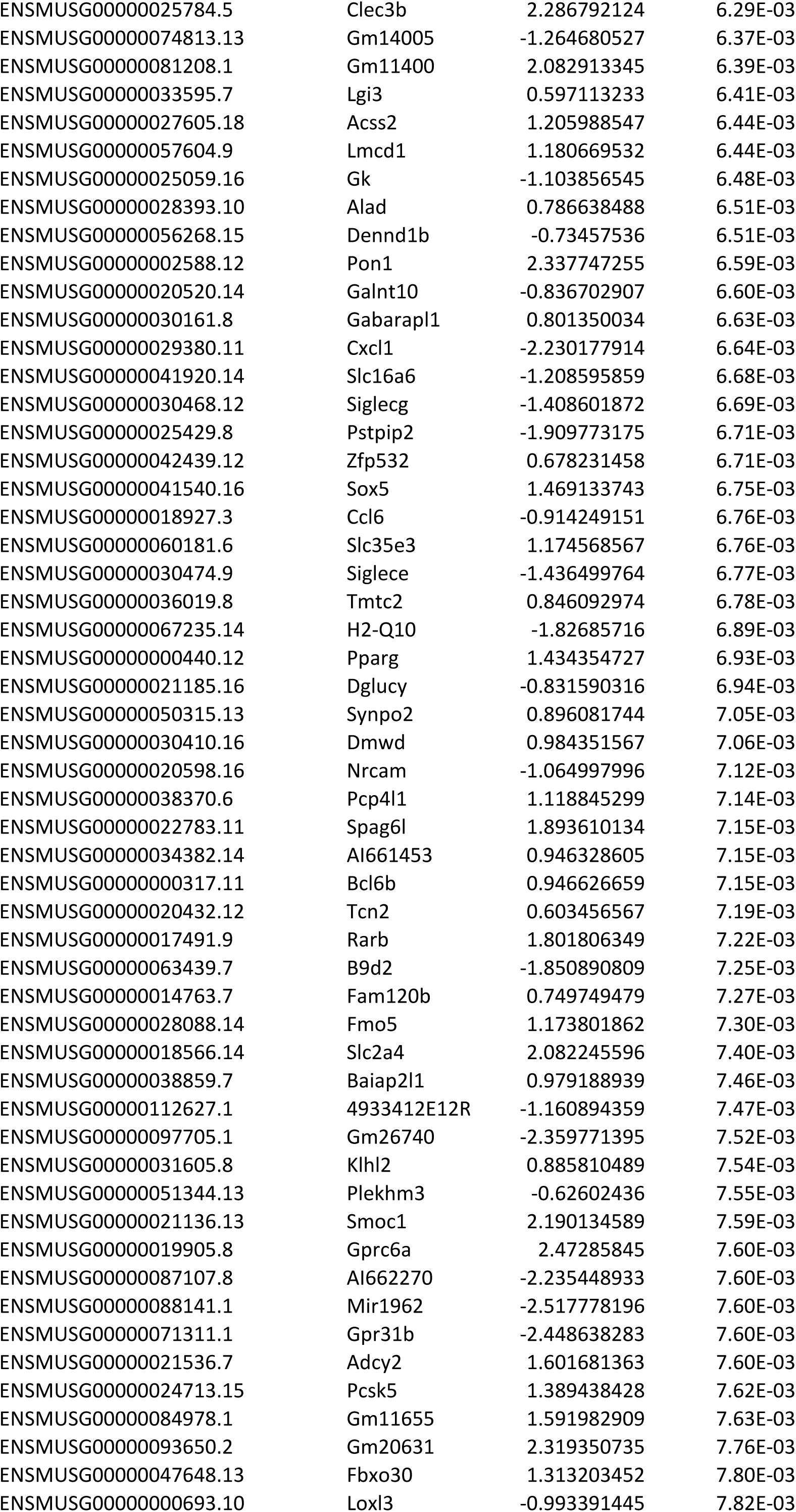
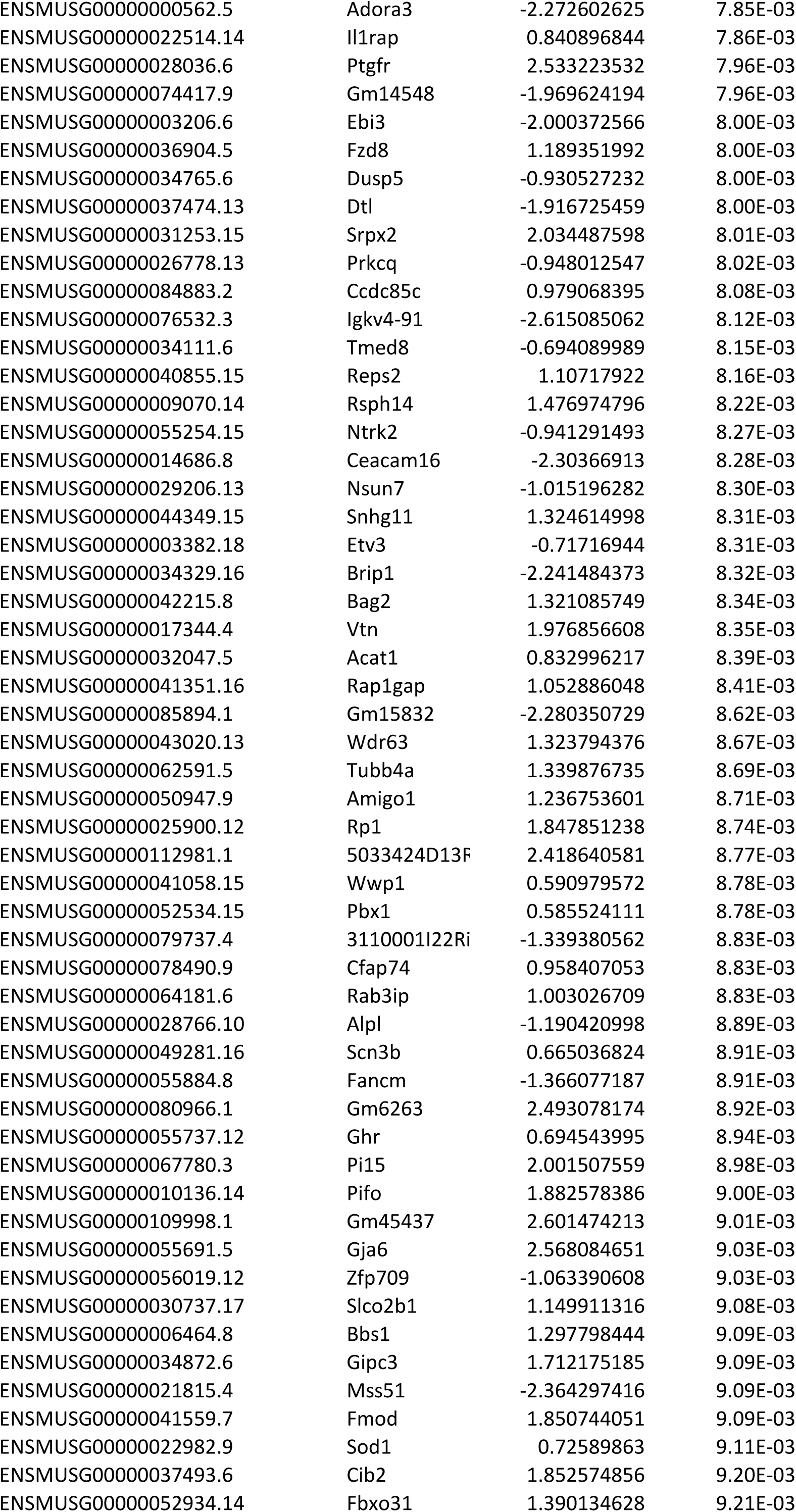
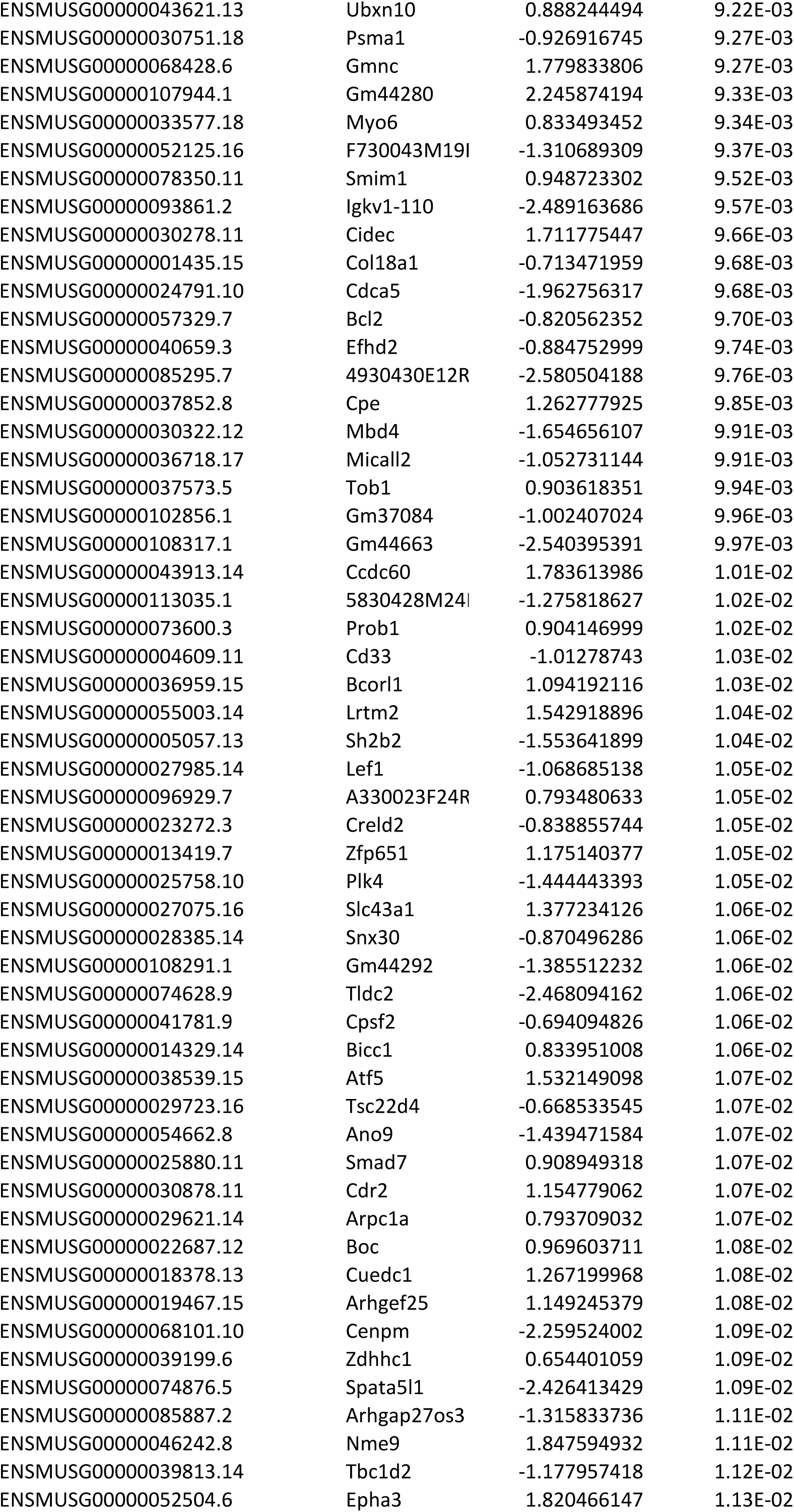
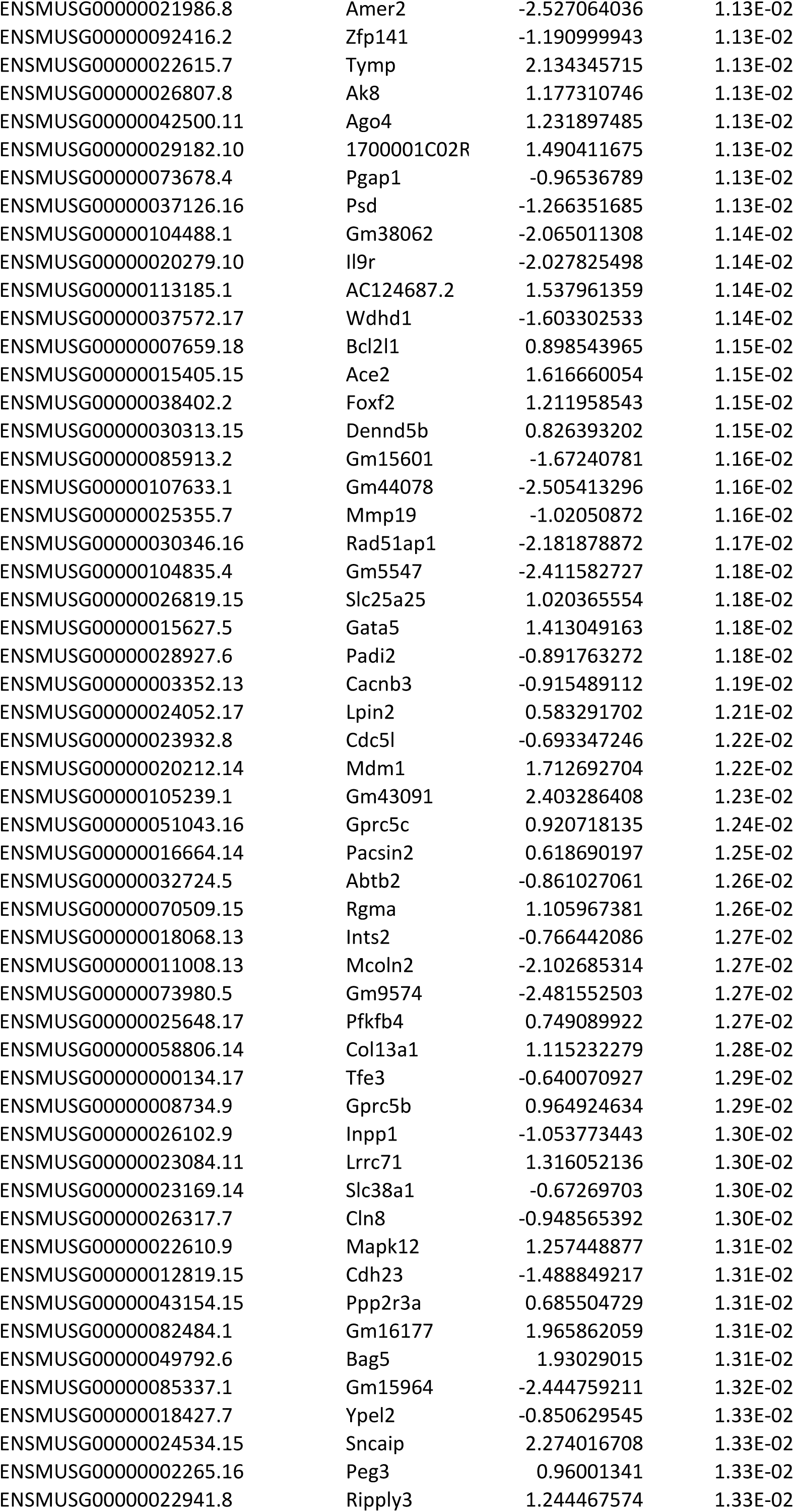
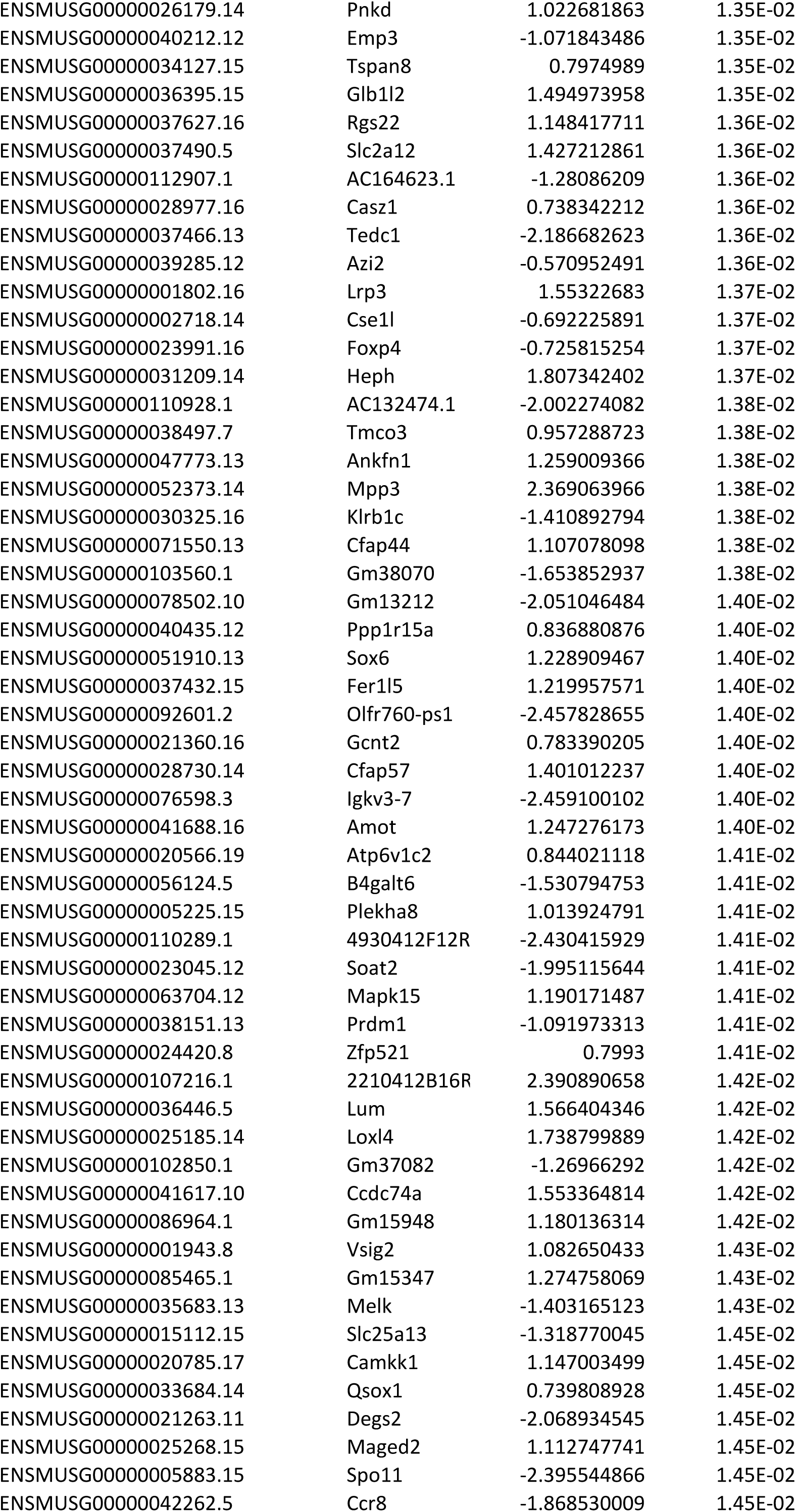
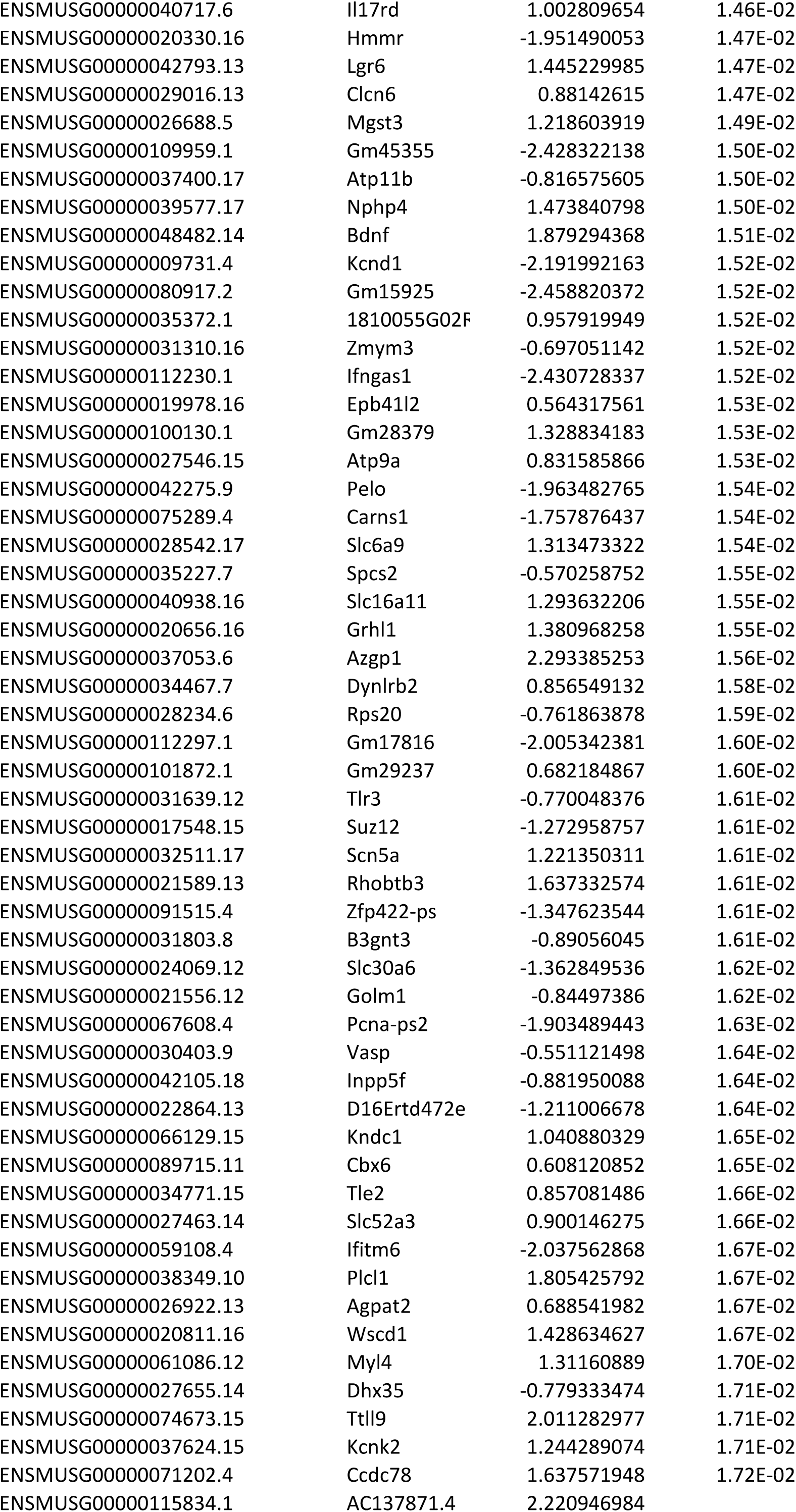
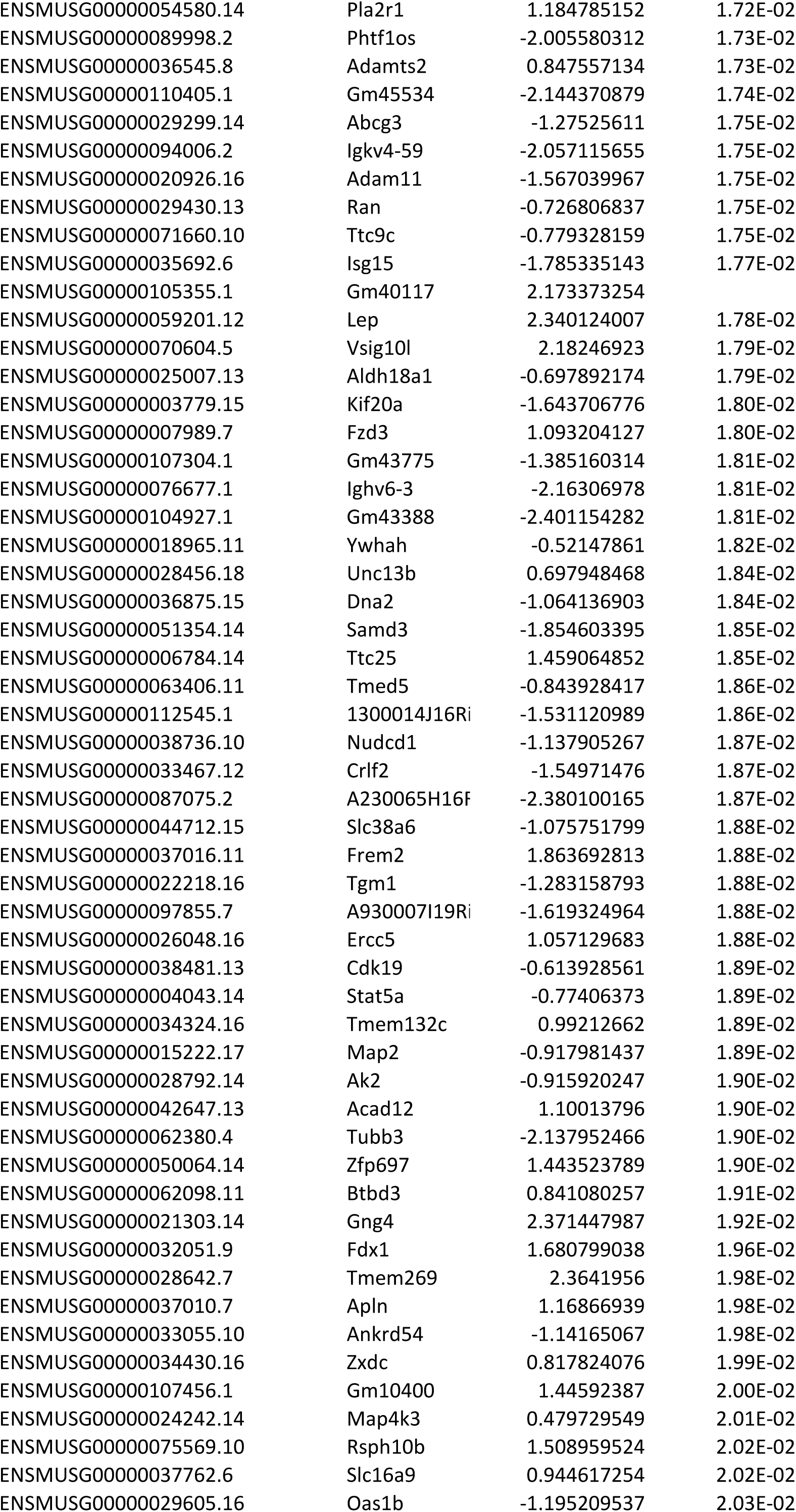
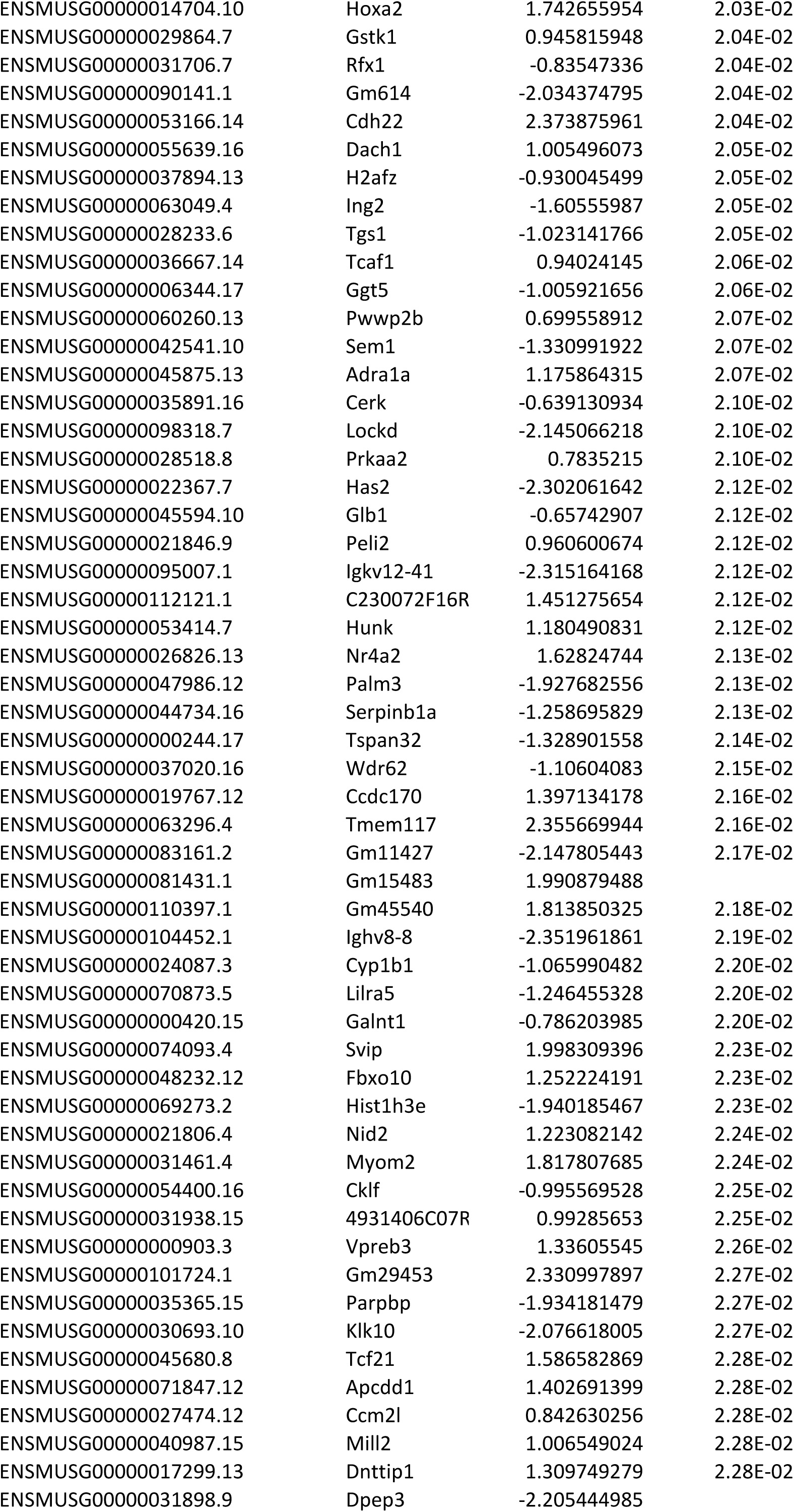
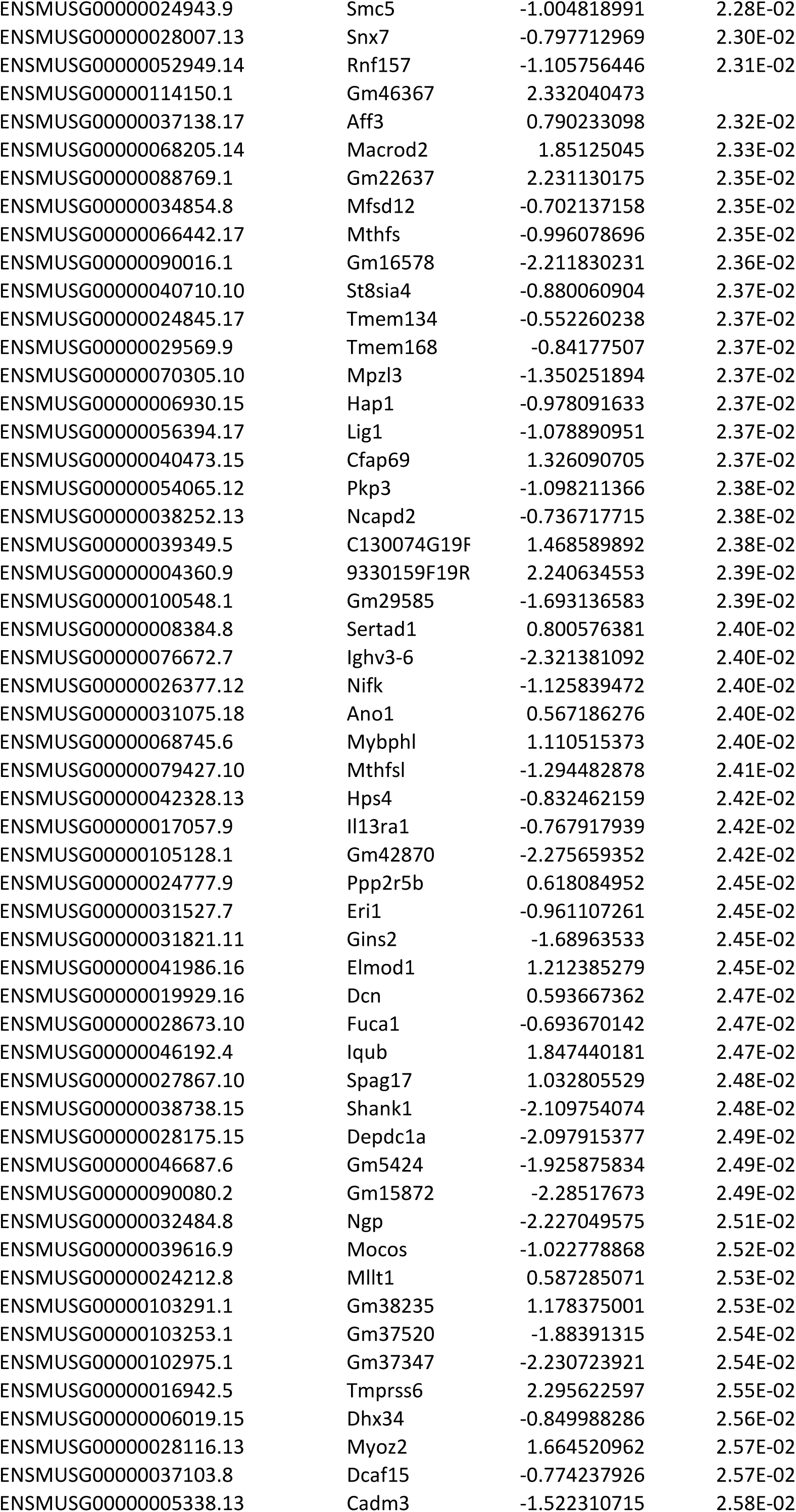
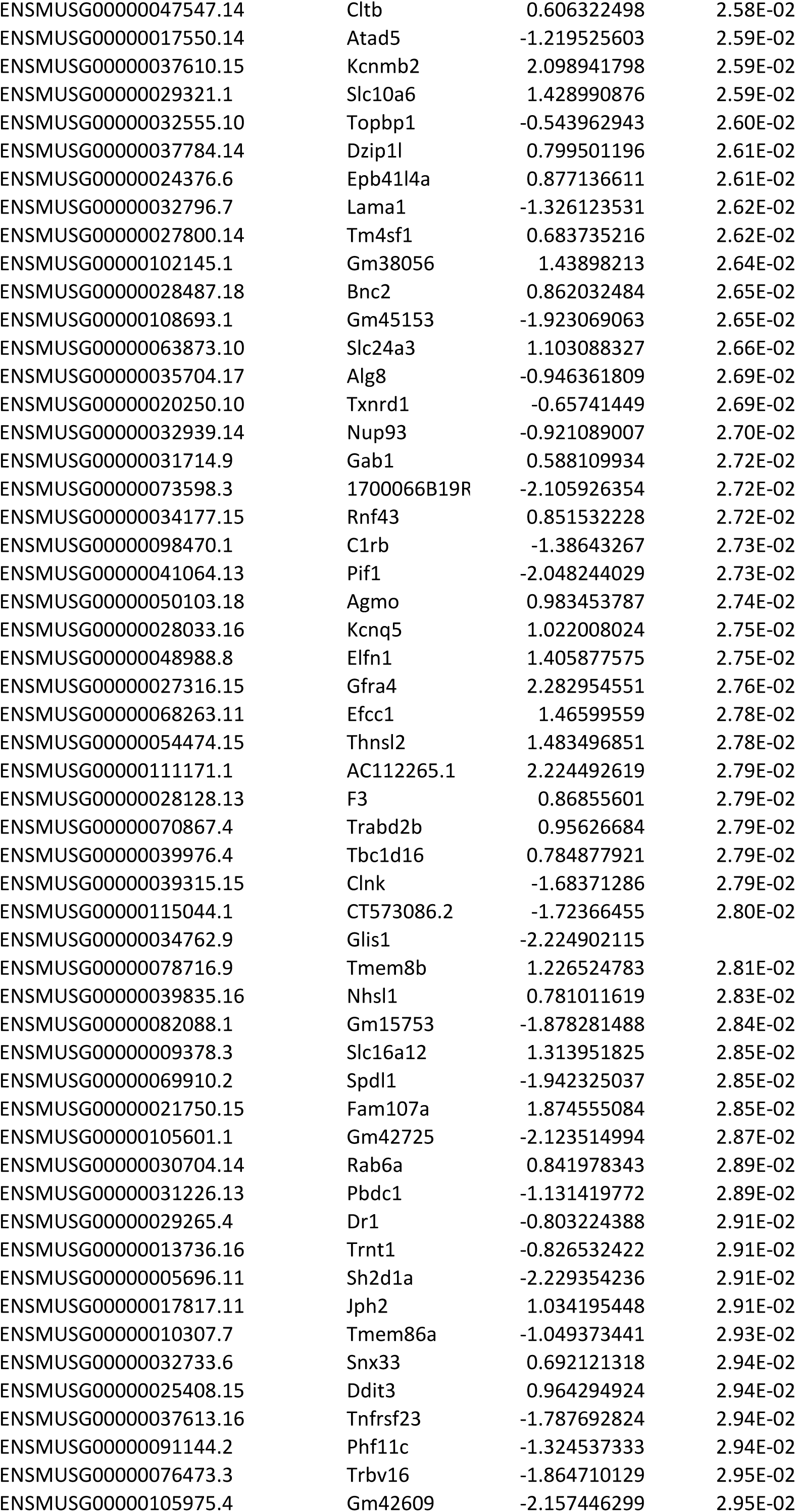
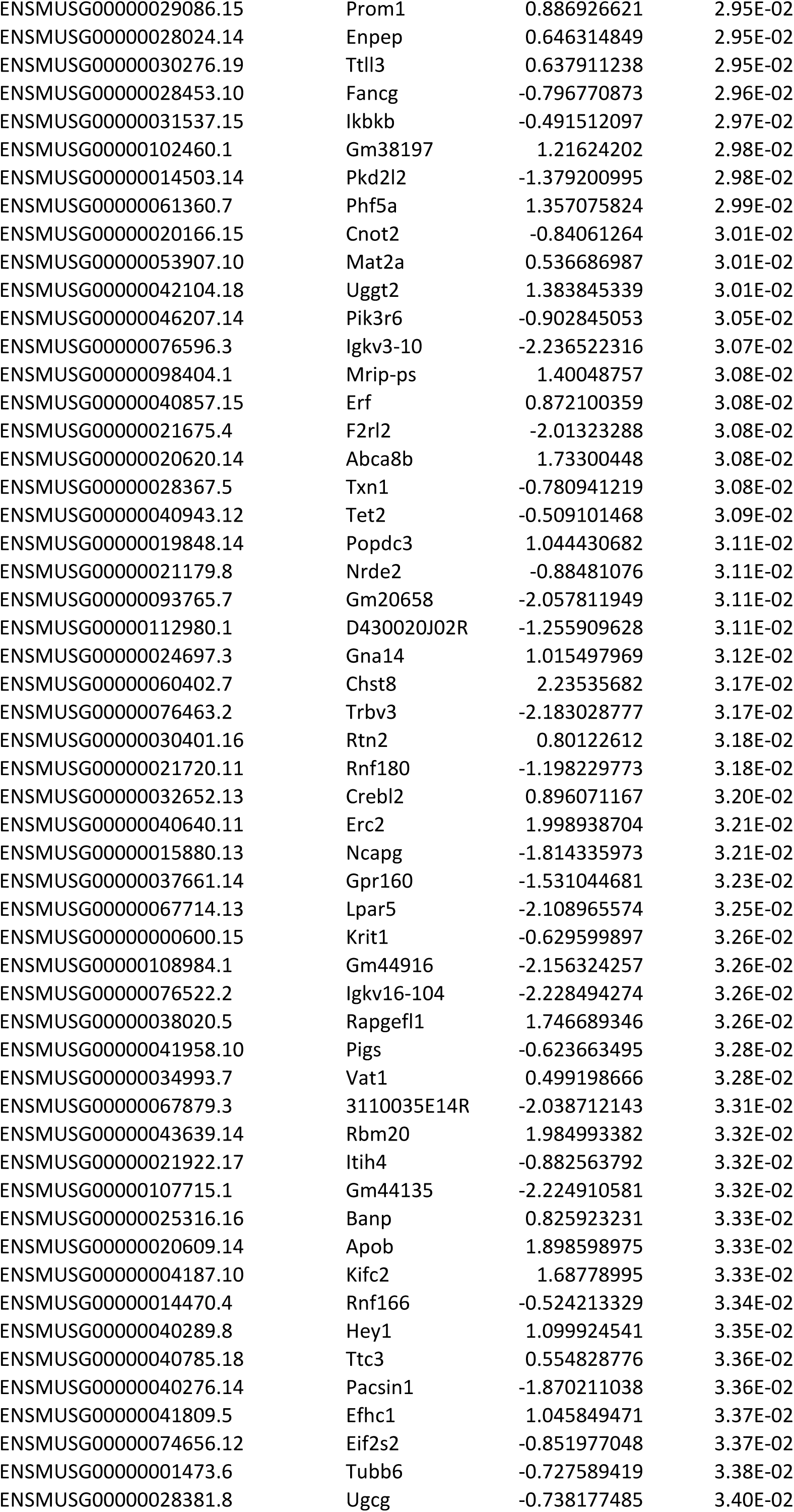
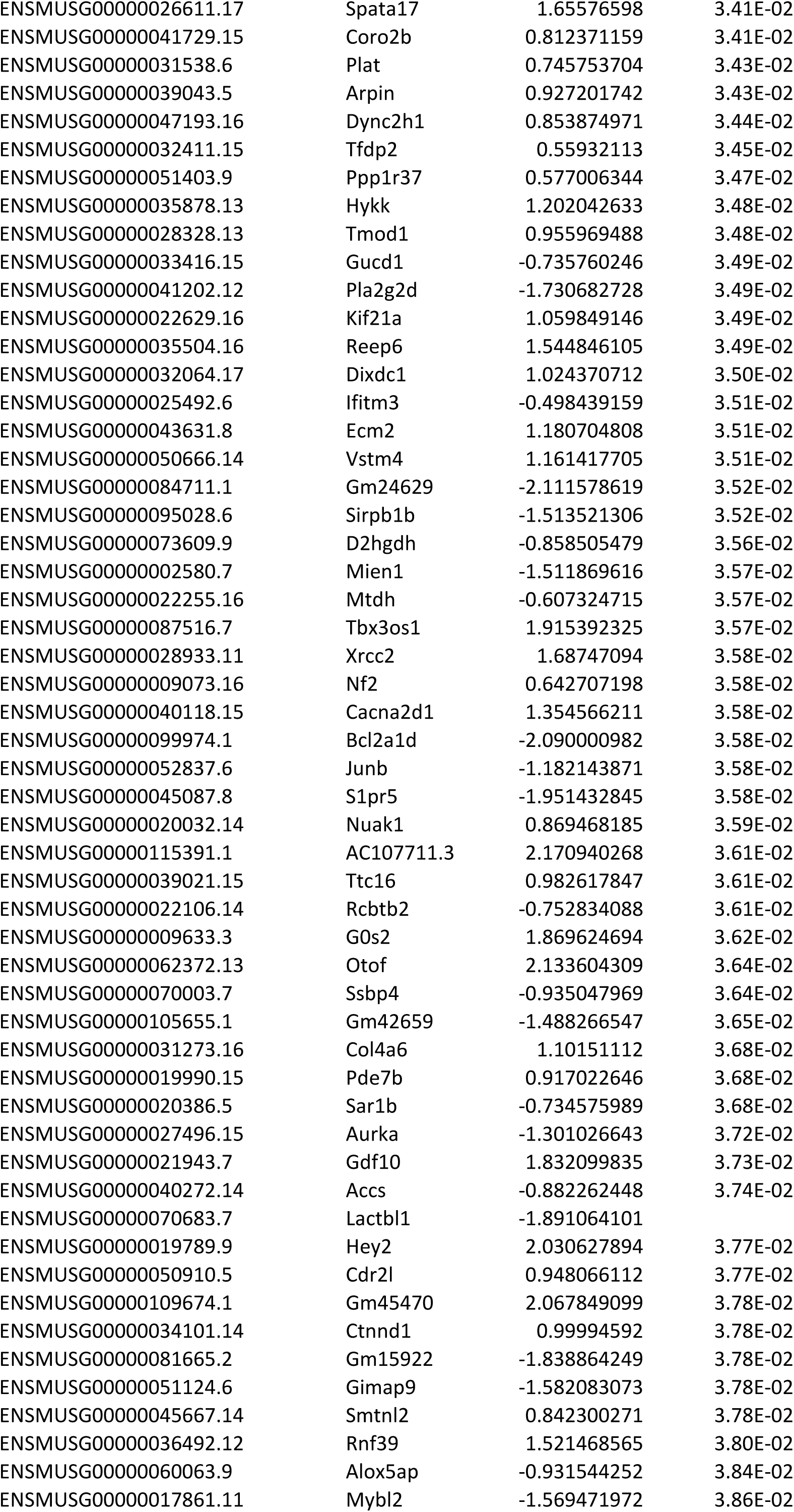
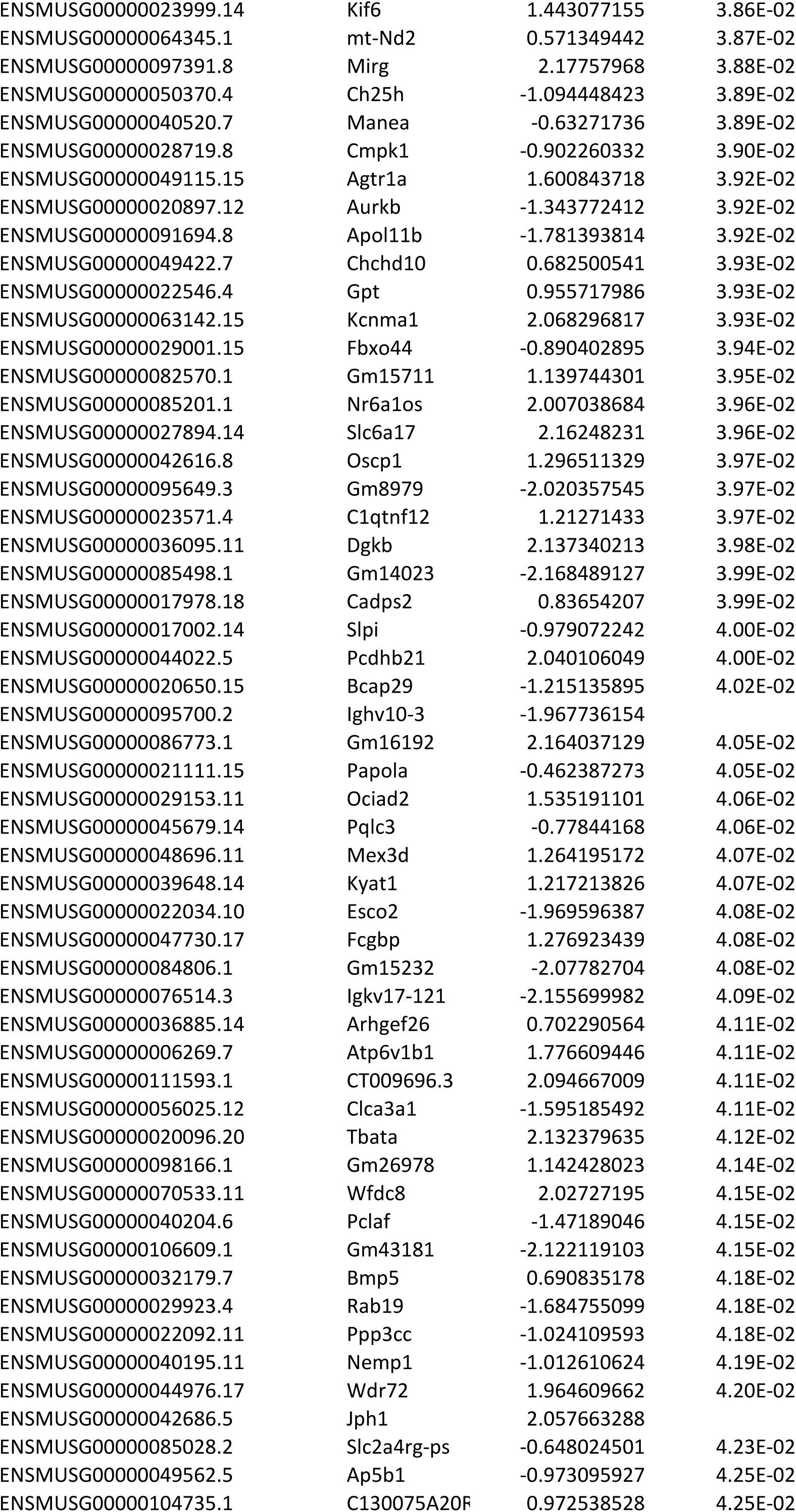
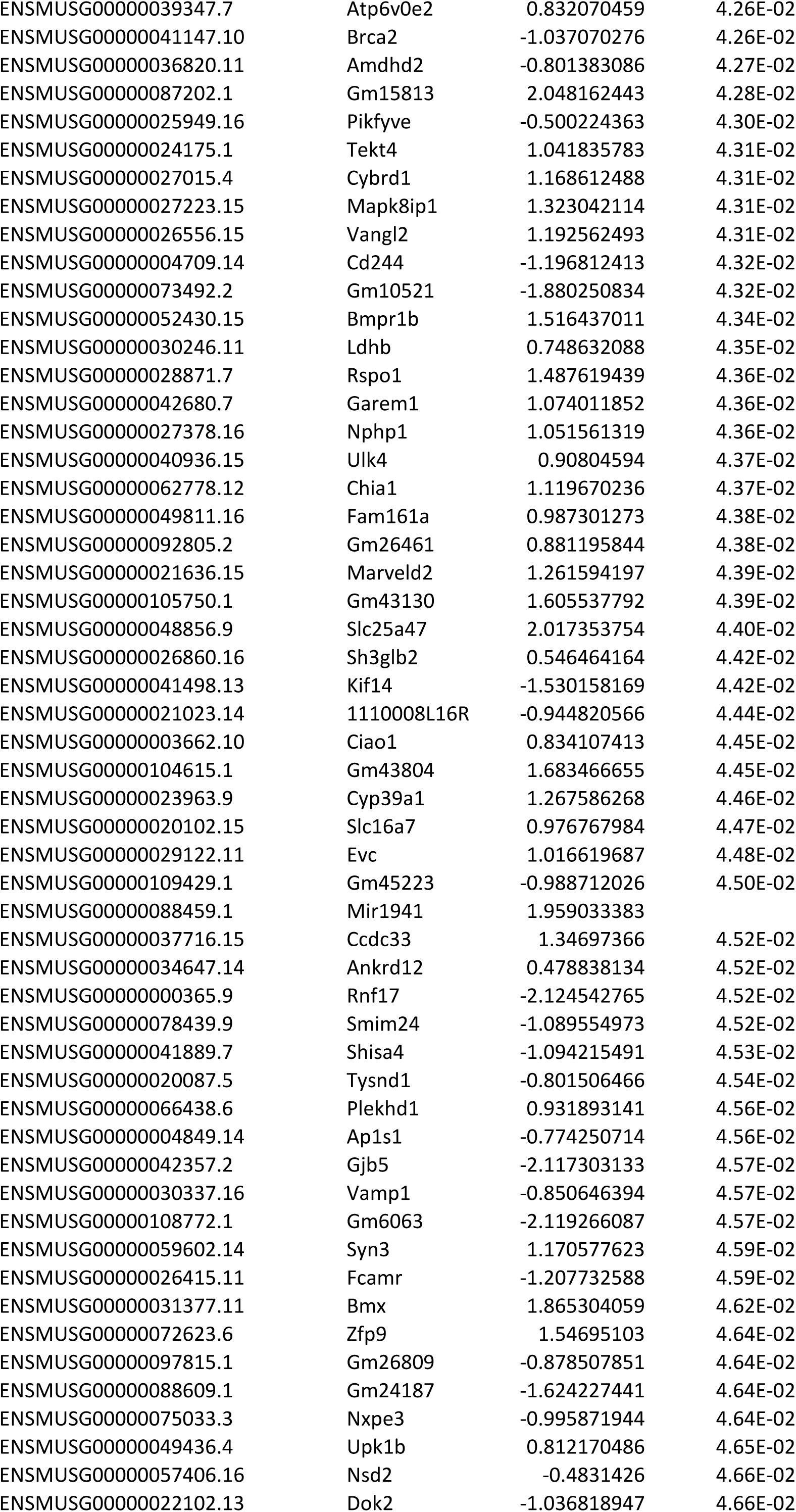
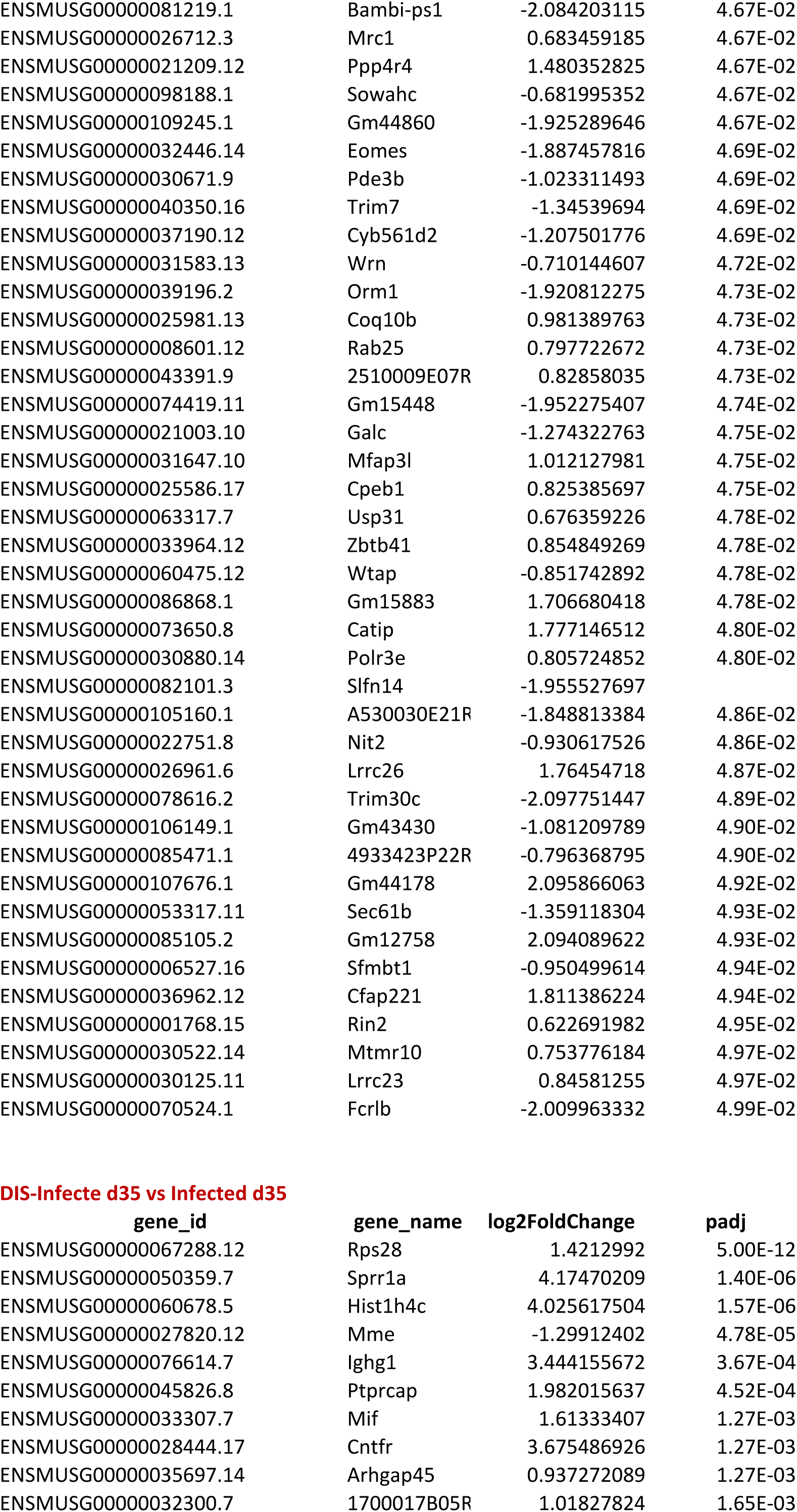
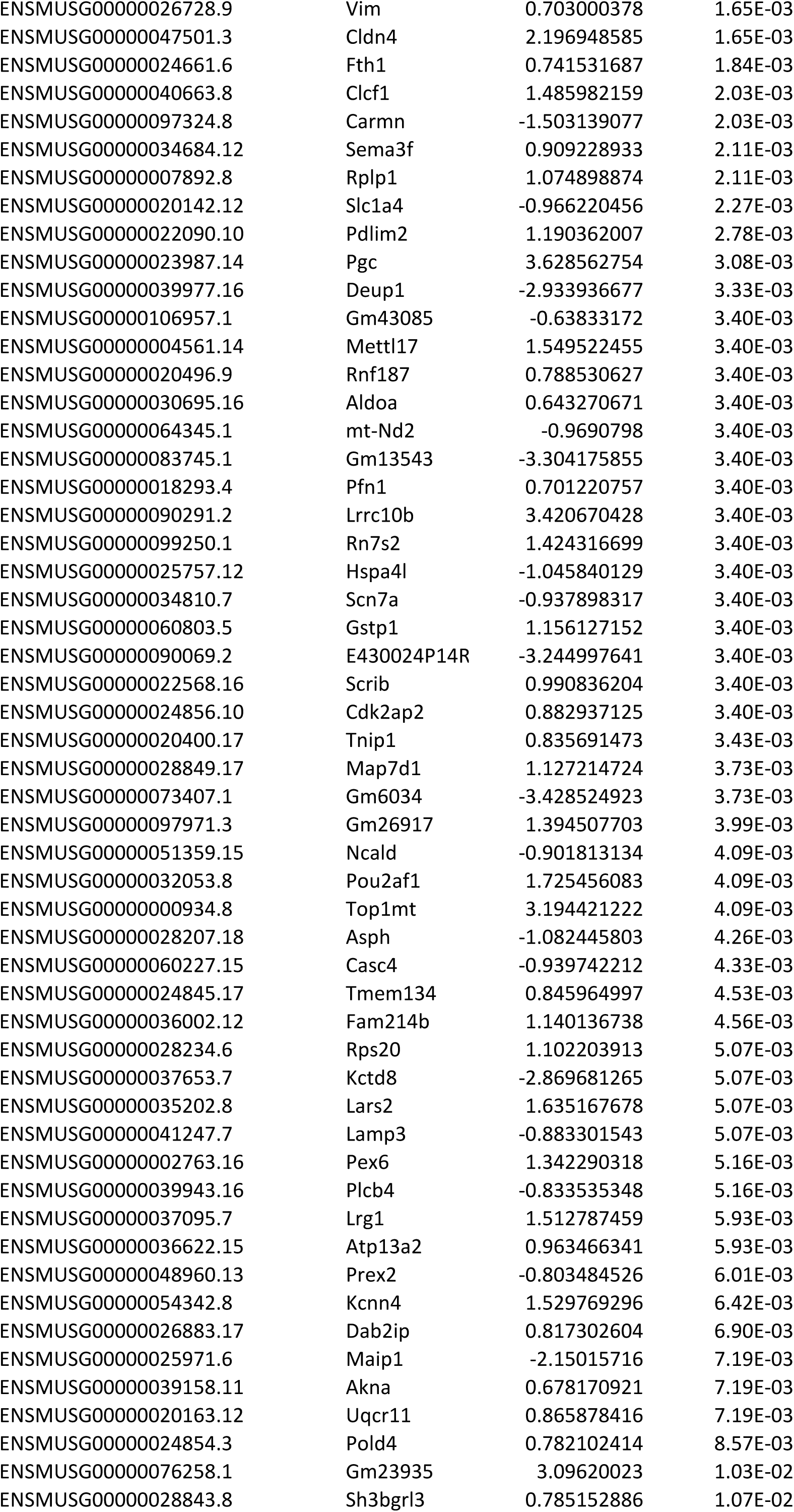
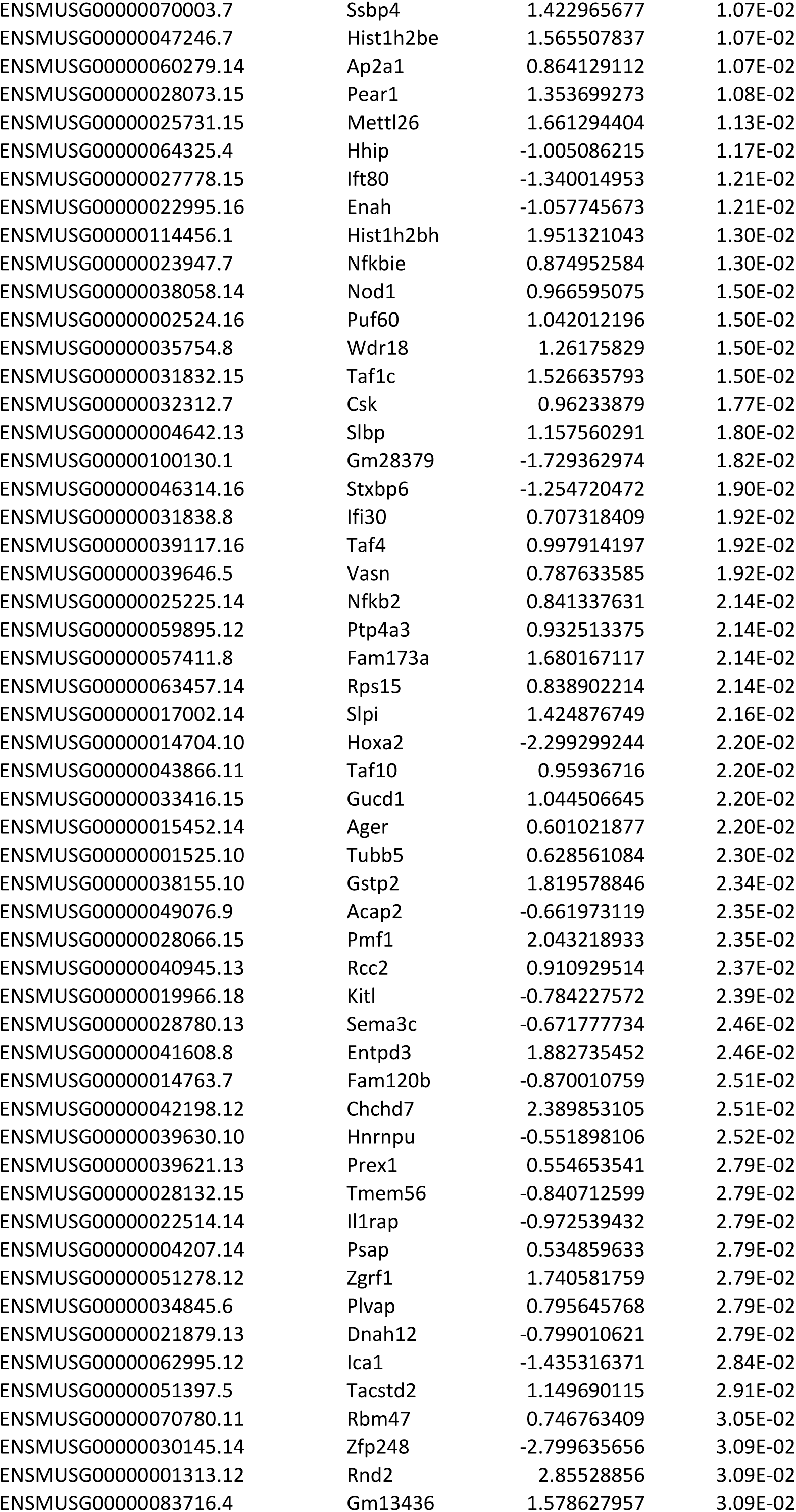
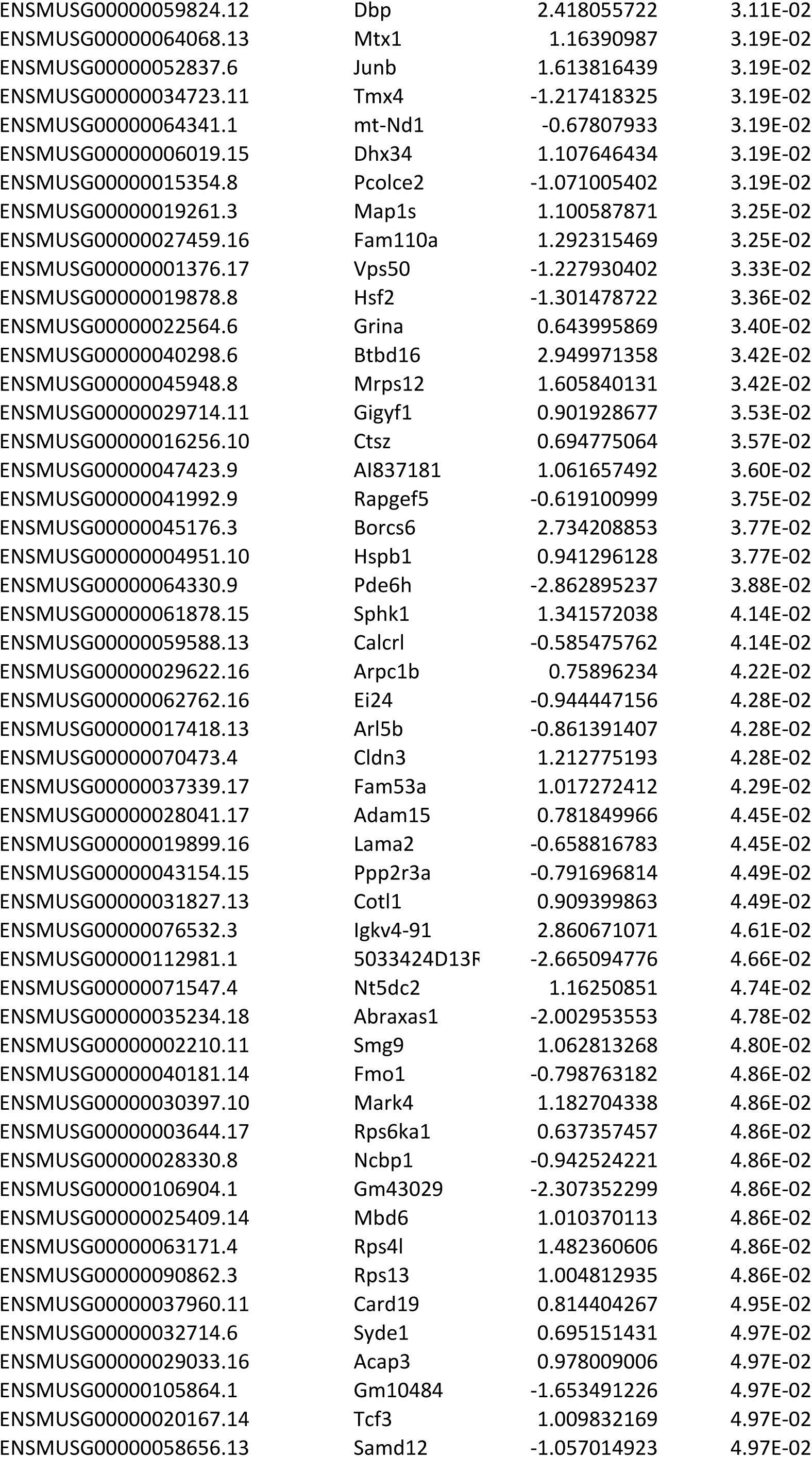
Differentially expressed genes between the RNA isolated from the lungs of uninfected, *Mtb-*infected and DIS-treated *Mtb*-infected mice at 4 wk p.t. Two sheets depict differentially expressed genes of uninfected vs *Mtb*-infected and DIS-treated vs untreated *Mt*b-infected animals.

**Table S3:**
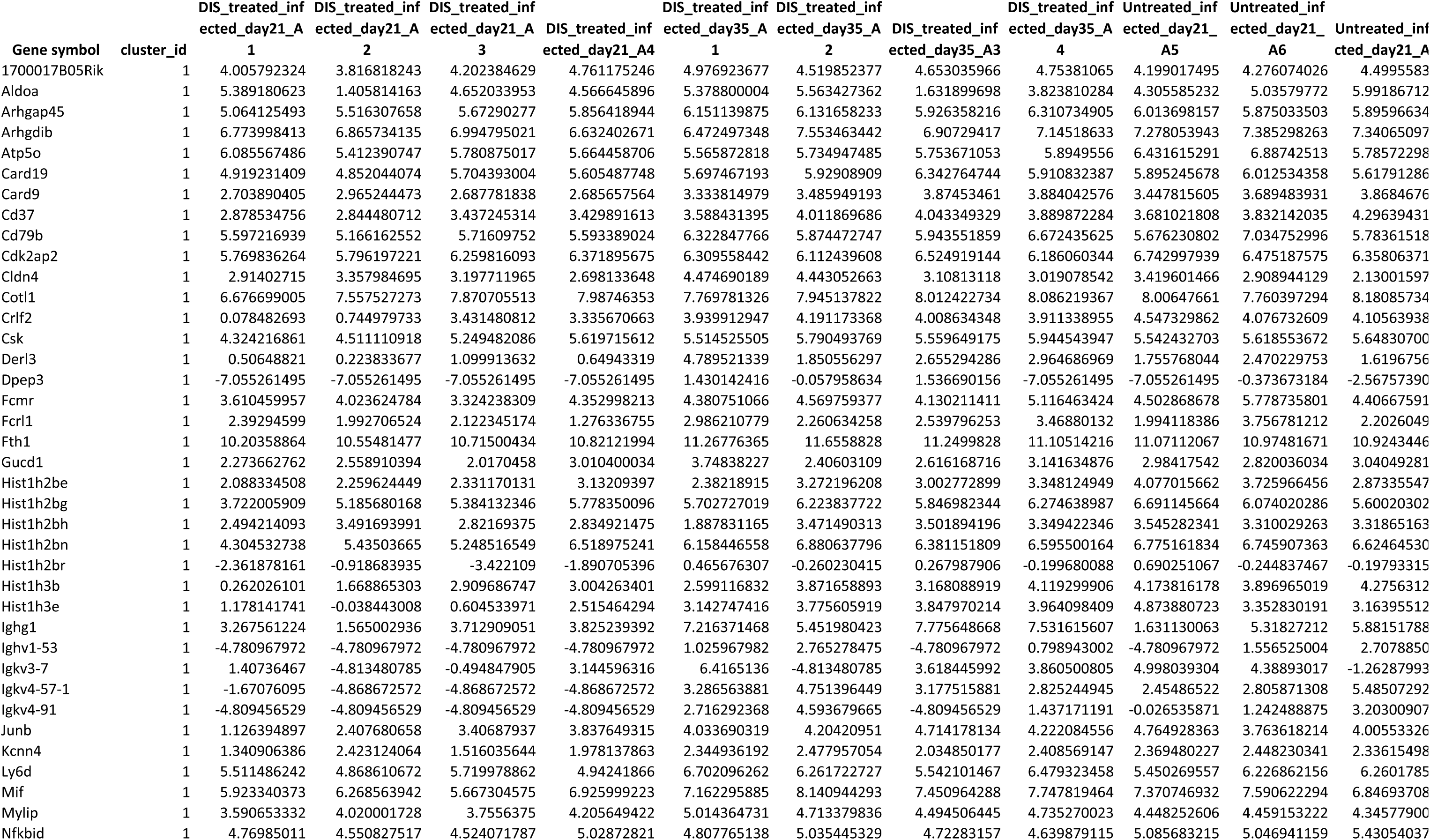
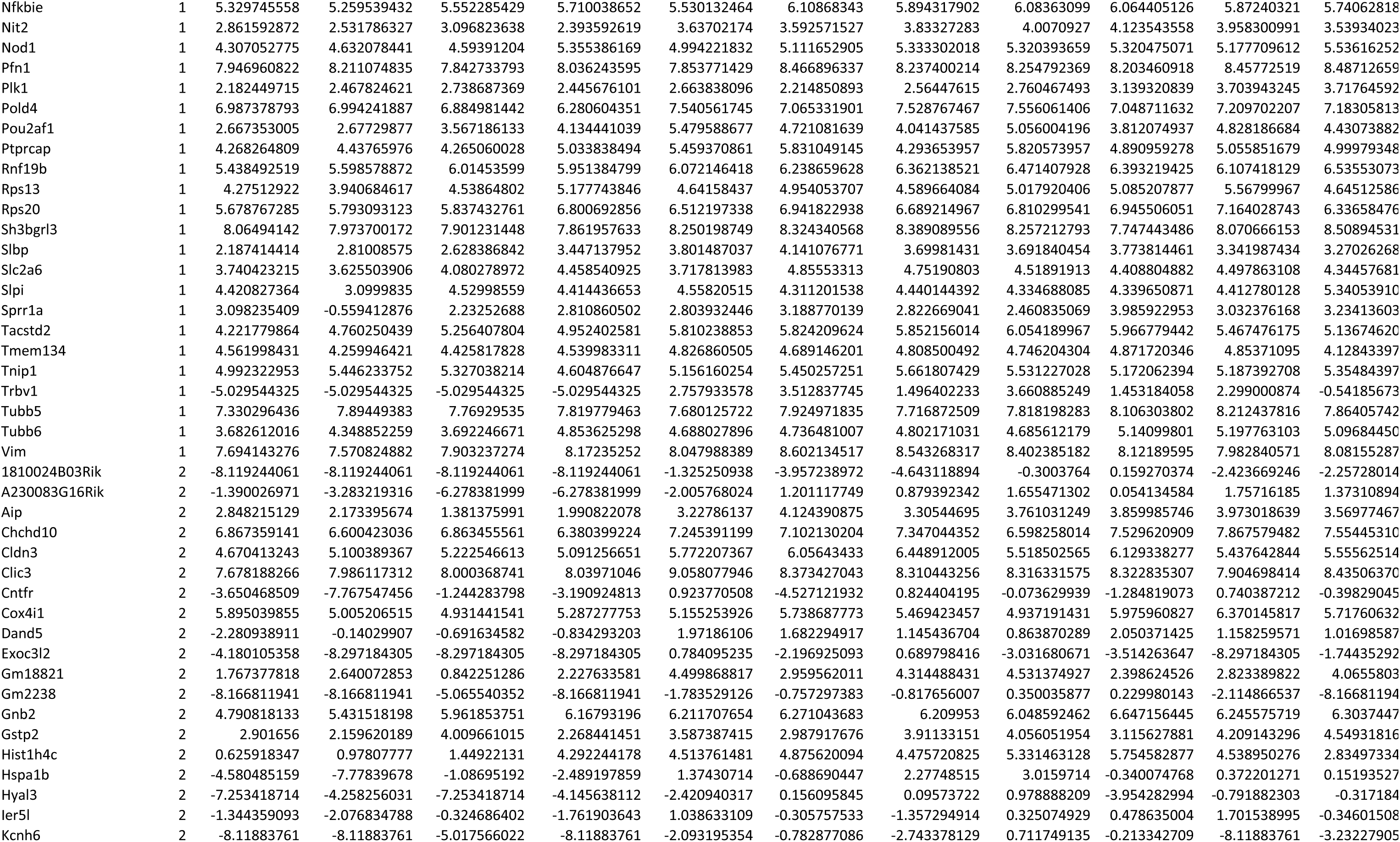
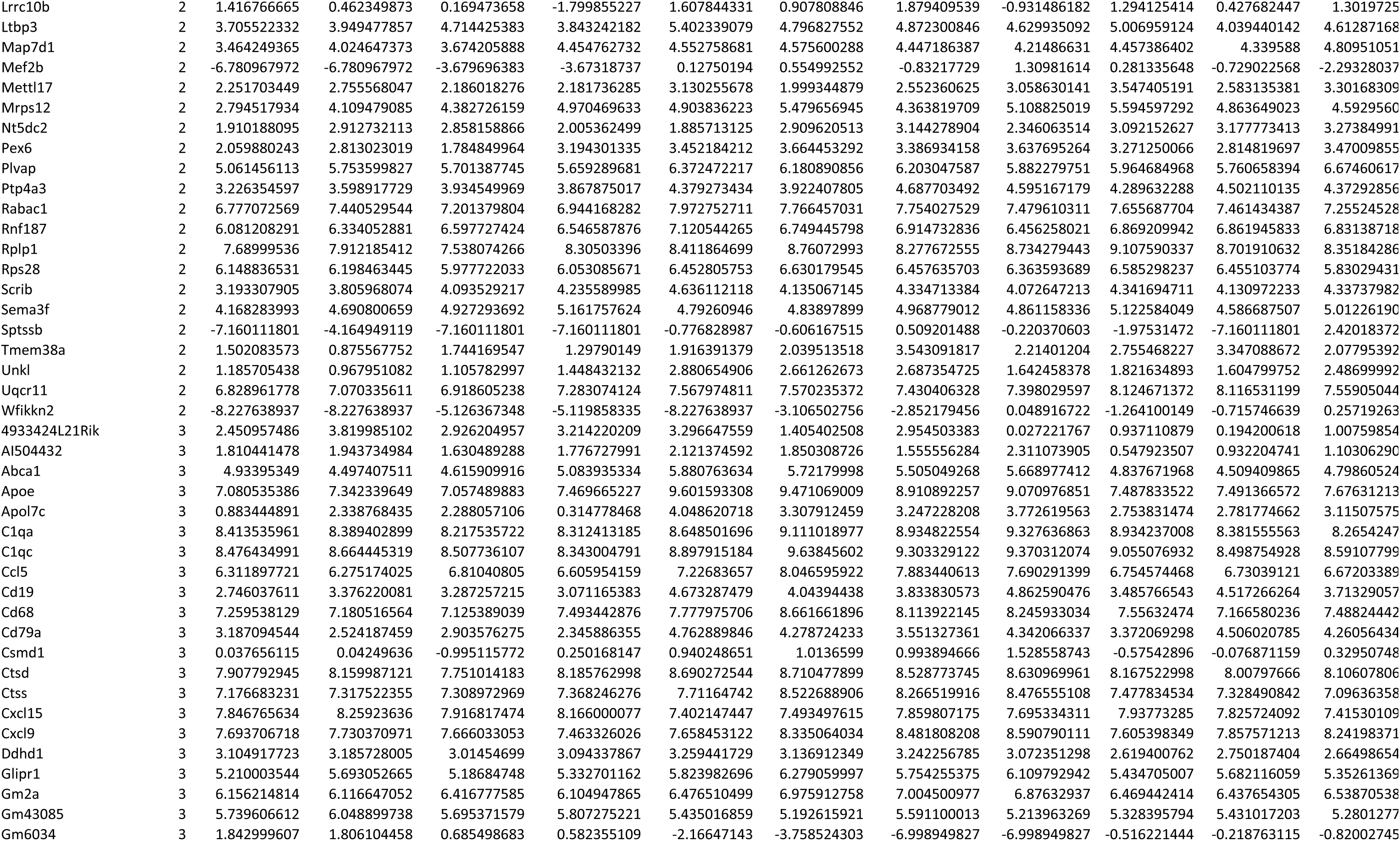
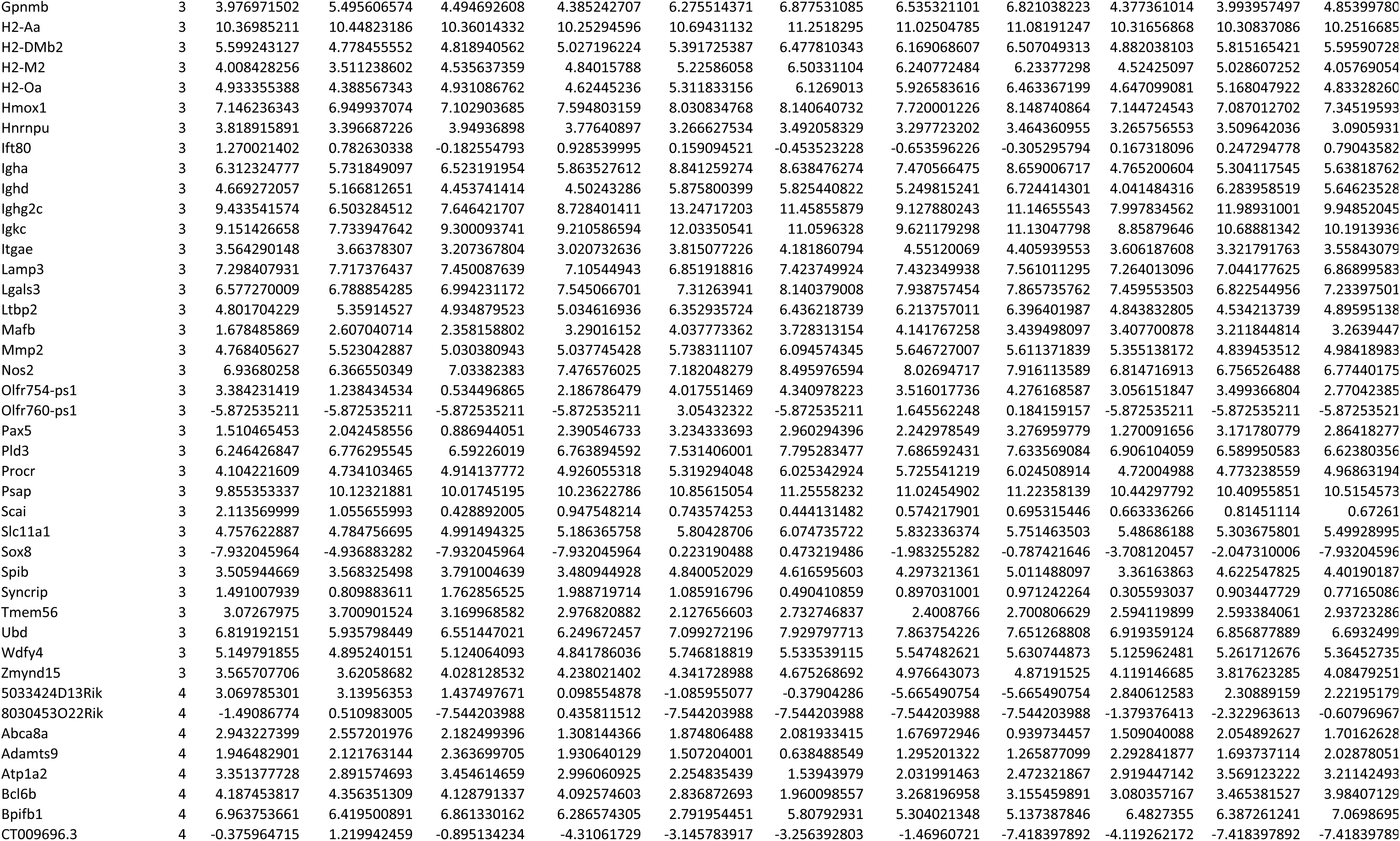
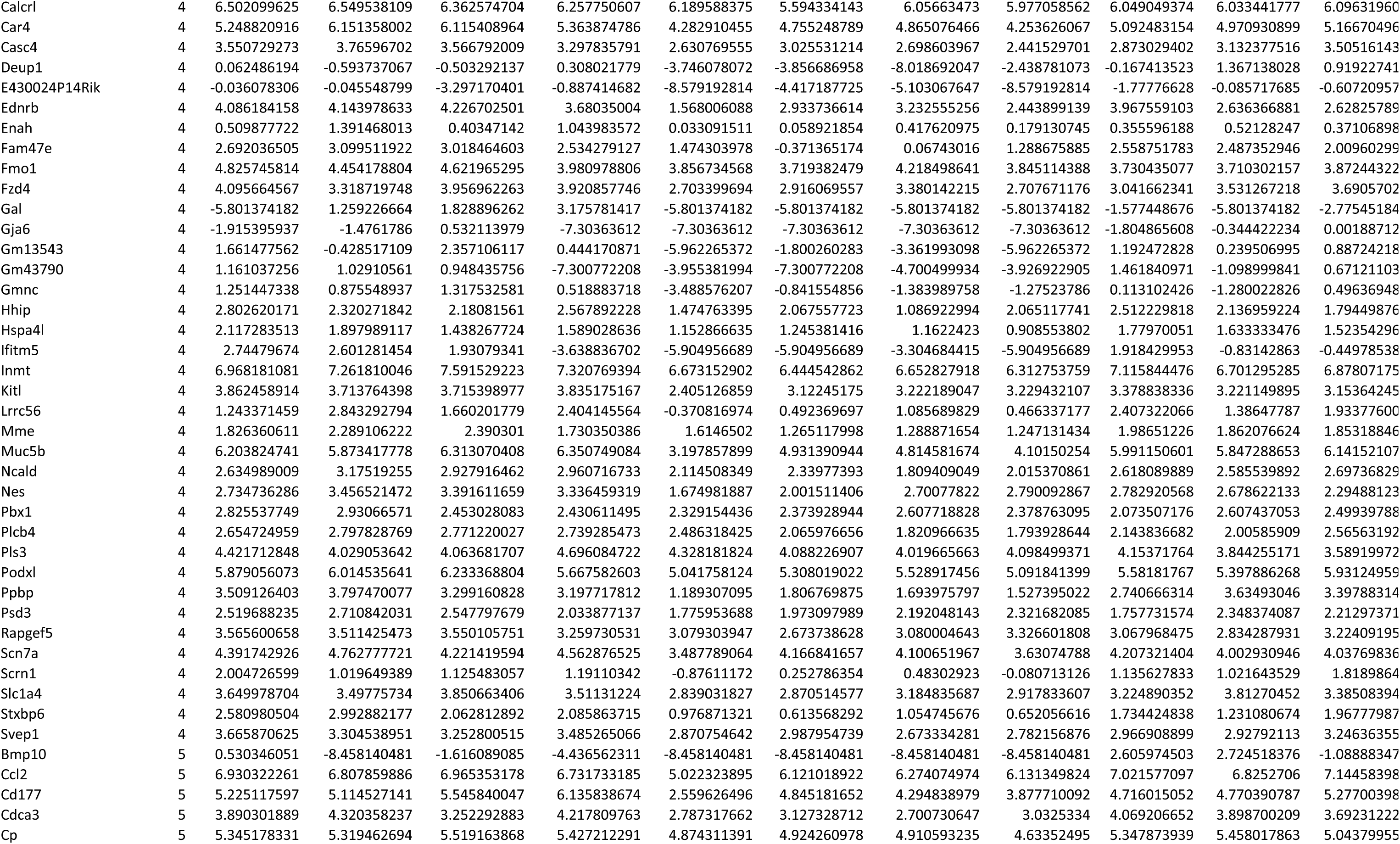
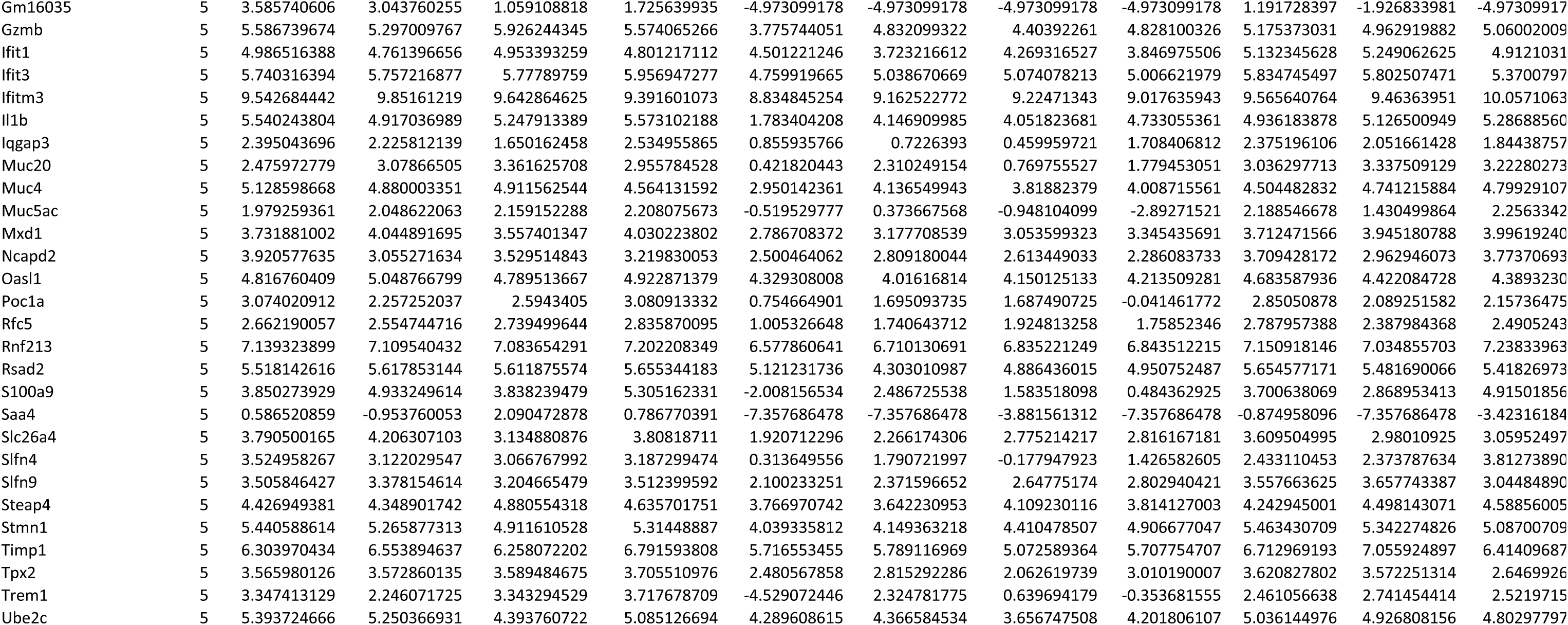
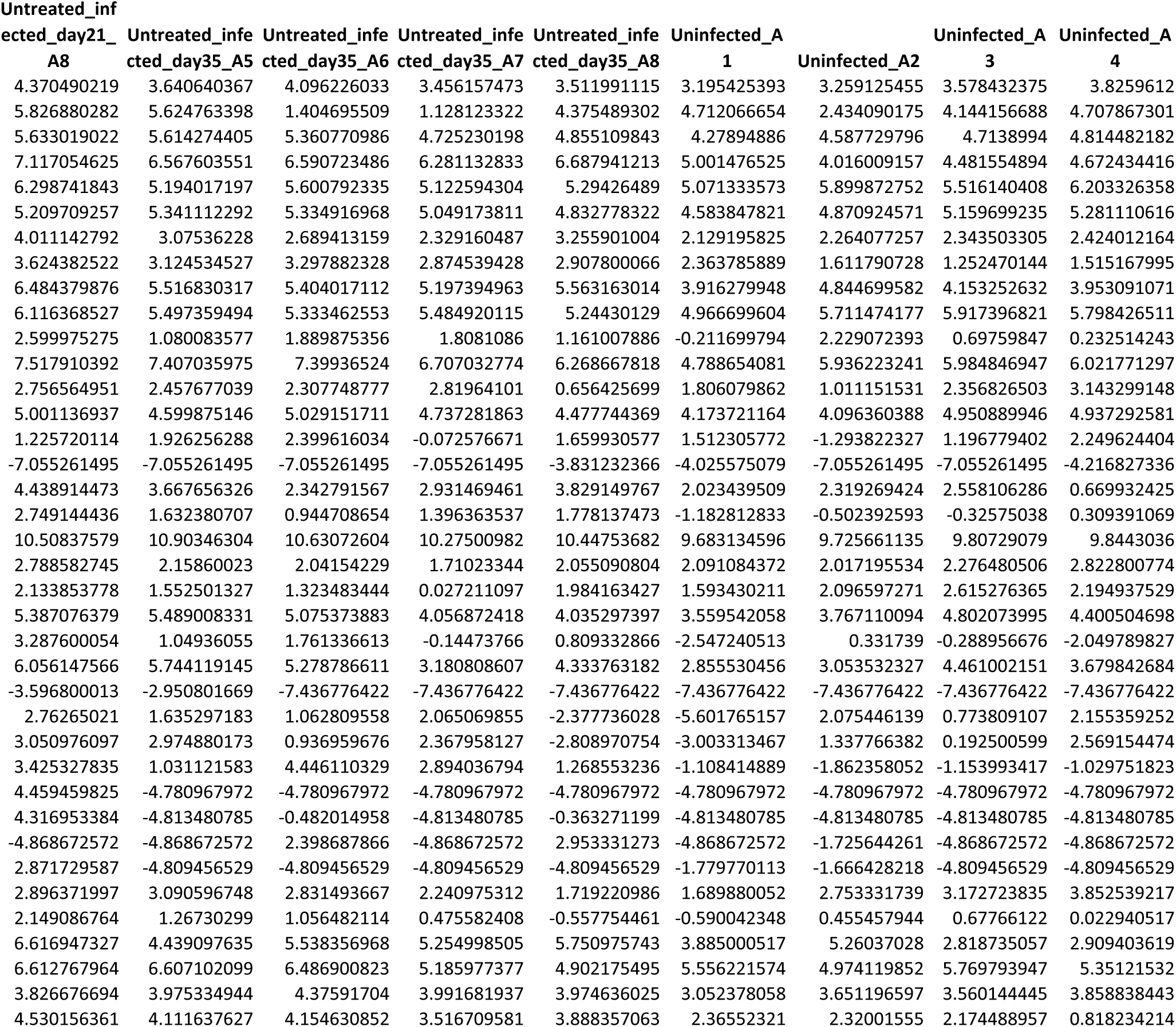
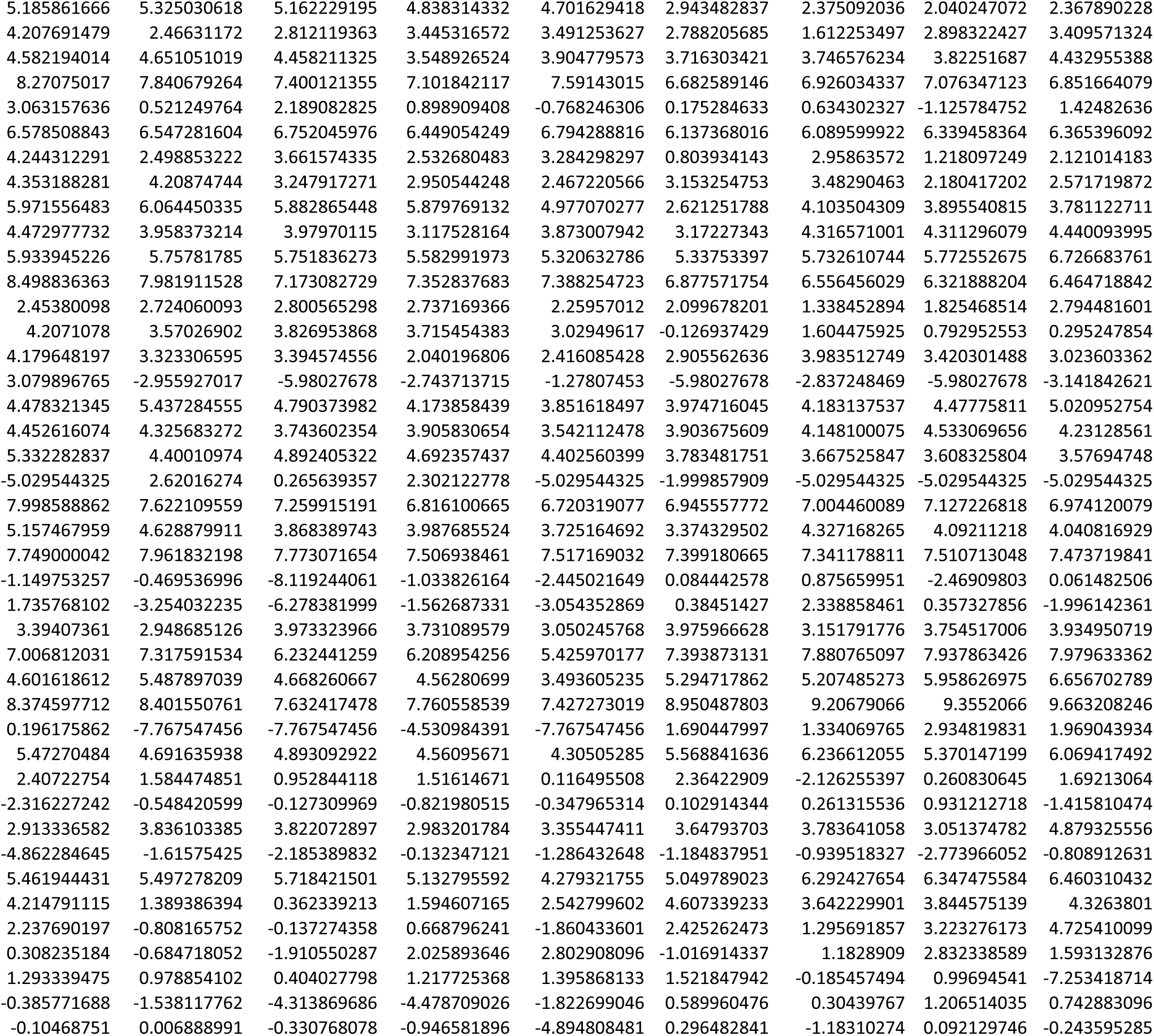
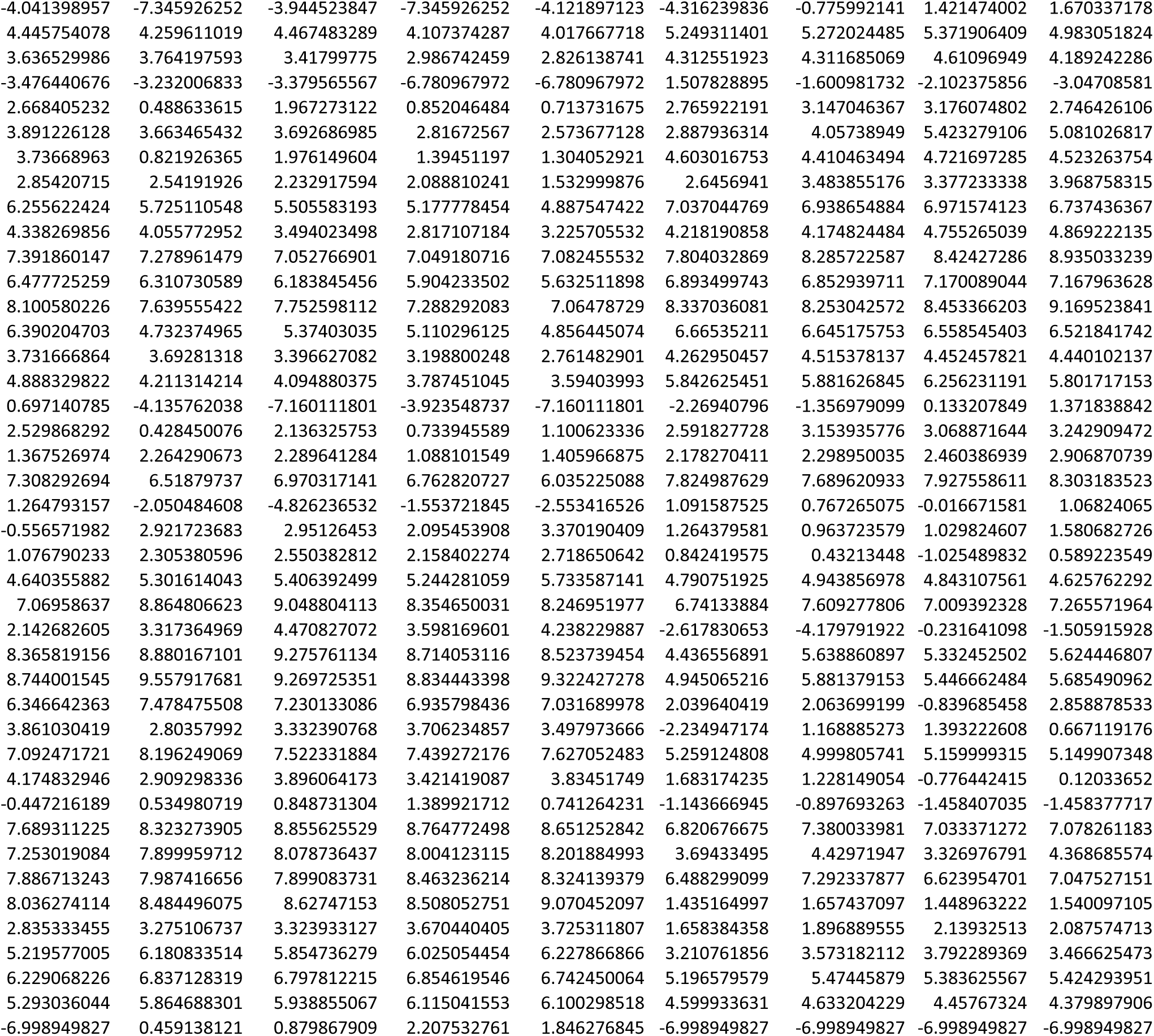
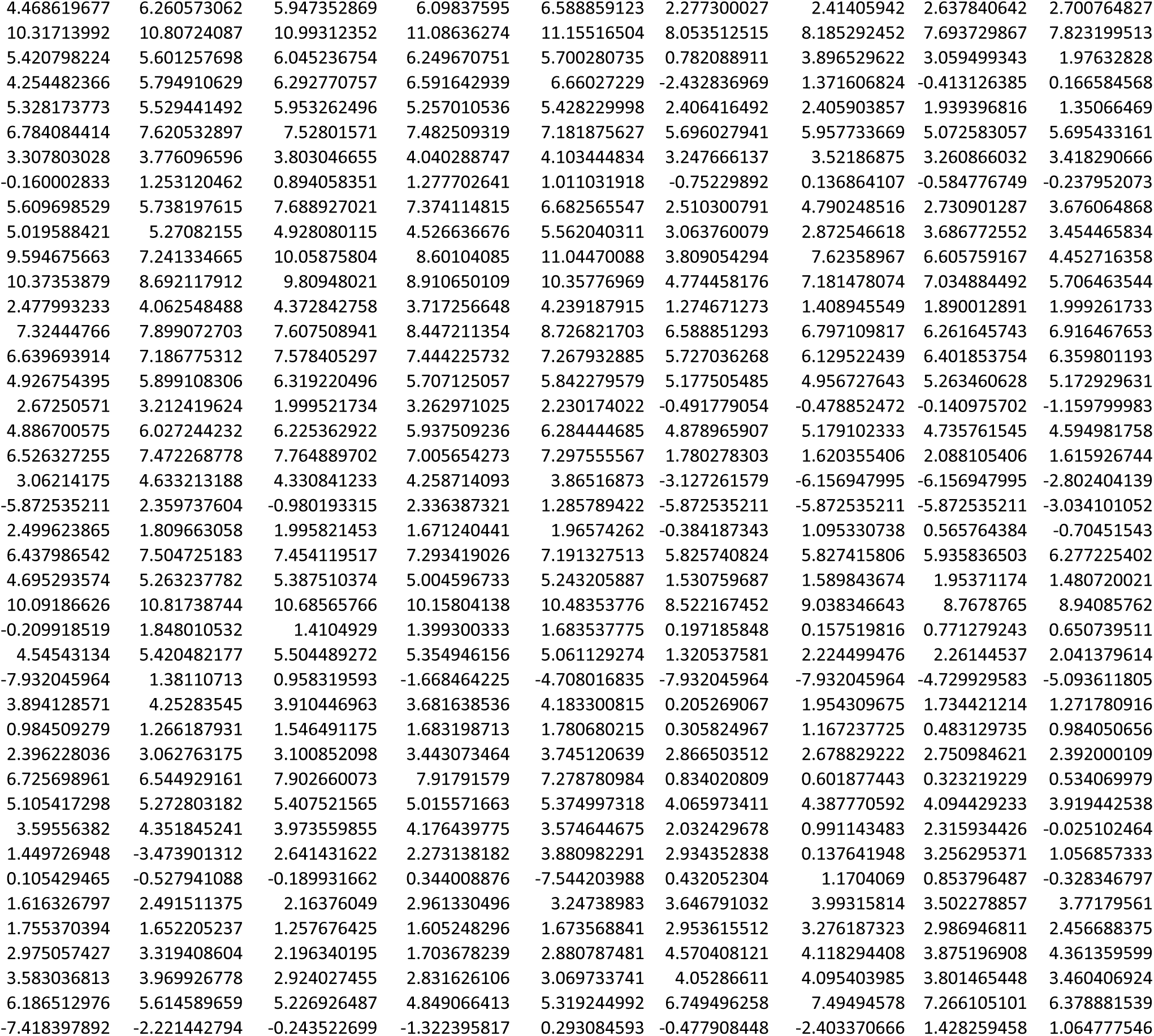
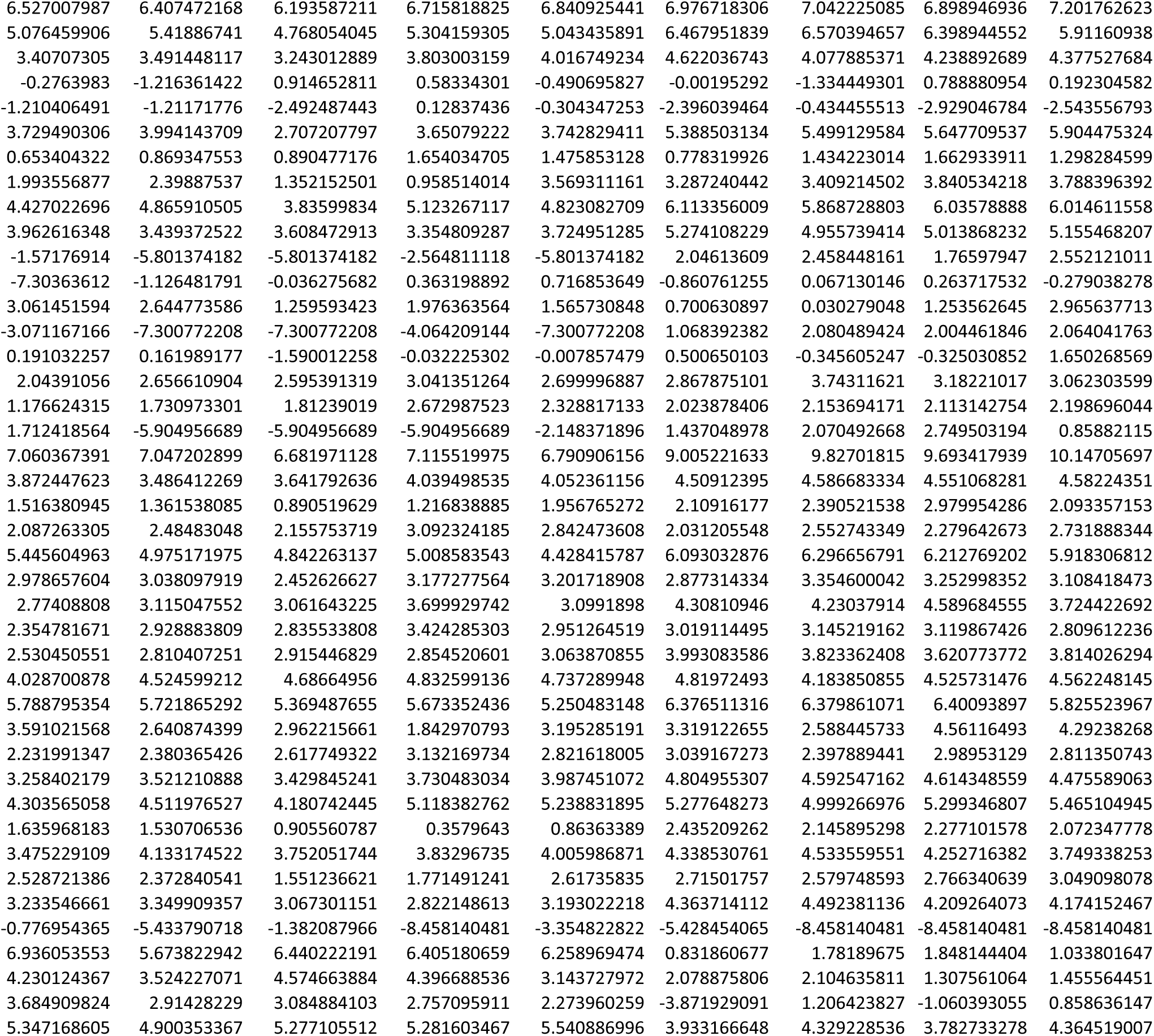
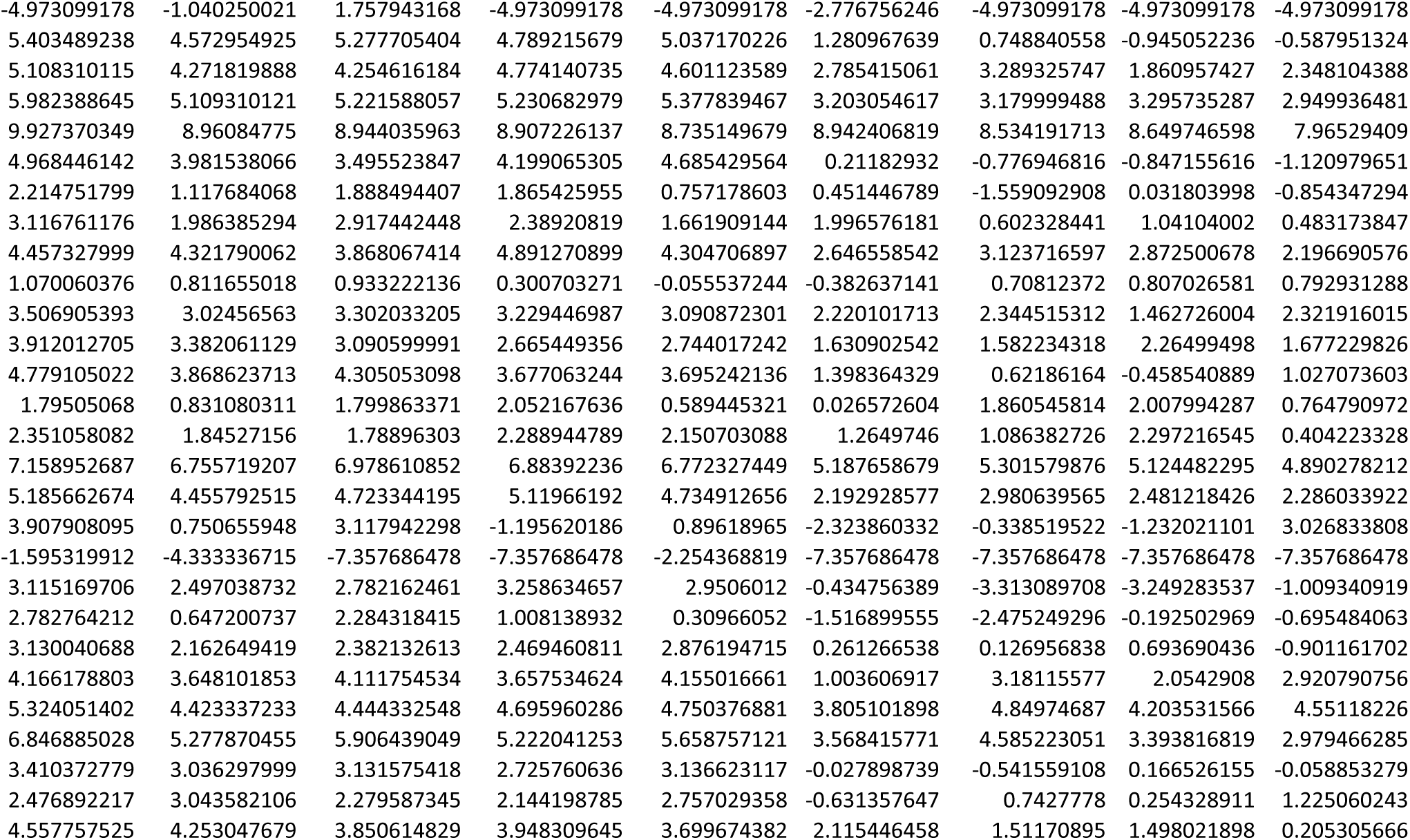
Genes modulated by time, infection and DIS treatment (genes identified as significantly changed during infection as well as being significantly changed during time or DIS treatment) are shown. Expression (log_2_ RPKM) of cluster specific genes (Figure 5A) is depicted for all groups.

